# An atlas of RNA-dependent proteins in cell division reveals the riboregulation of mitotic protein-protein interactions

**DOI:** 10.1101/2024.09.25.614981

**Authors:** Varshni Rajagopal, Jeanette Seiler, Isha Nasa, Simona Cantarella, Jana Theiss, Franziska Herget, Bianca Kaifer, Martin Schneider, Dominic Helm, Julian König, Kathi Zarnack, Sven Diederichs, Arminja N. Kettenbach, Maïwen Caudron-Herger

## Abstract

Ribonucleoprotein complexes are dynamic assemblies of RNA with RNA-binding proteins (RBPs), which can modulate the fate of the RNA molecules from transcription to degradation. Vice versa, RNA can regulate the interactions and functions of the associated proteins. Dysregulation of RBPs is linked to diseases such as cancer and neurological disorders. RNA and RBPs are present in mitotic structures like the centrosomes and spindle microtubules, but their influence on mitotic spindle integrity remains unknown. Thus, we applied the R-DeeP strategy for the proteome-wide identification of RNA-dependent proteins and complexes to cells synchronized in mitosis versus interphase. The resulting atlas of RNA-dependent proteins in cell division can be accessed through the R-DeeP 3.0 database (R-DeeP3.dkfz.de). It revealed key mitotic factors as RNA-dependent such as AURKA, KIFC1 and TPX2 that were linked to RNA despite their lack of canonical RNA-binding domains. KIFC1 was identified as a new interaction partner and phosphorylation substrate of AURKA at S^349^ and T^359^. In addition, KIFC1 interacted with both, AURKA and TPX2, in an RNA-dependent manner. Our data suggest a riboregulation of mitotic protein-protein interactions during spindle assembly, offering new perspectives on the control of cell division processes by RNA-protein complexes.

**Highlights:**

- Differential R-DeeP screens in mitosis and interphase are provided as a resource in a user-friendly database at R-DeeP3.dkfz.de
- An atlas of RNA-dependent proteins in cell division identifies a substantial number of unconventional RNA-binding proteins among mitotic factors
- Investigation of protein-protein interactions reveals KIFC1 as a new AURKA and TPX2 interaction partner during spindle assembly
- KIFC1, AURKA and TPX2 interact with each other in an RNA-dependent manner and directly bind to RNA
- AURKA phosphorylates KIFC1 at residues S^349^ and T^359^

## INTRODUCTION

From the onset of transcription in the nucleus until their degradation in the cytoplasm, RNA transcripts associate with RNA-binding proteins (RBPs) to form ribonucleoprotein (RNP) complexes (1). RNPs are dynamic macromolecular assemblies that regulate the fate of RNA molecules by coordinating all aspects of their post-transcriptional maturation and regulation such as splicing, modification, transport, translation and decay (2,3). Conversely, recent studies pointed to a more RNA-centric view of RNA-protein interactions, termed “riboregulation”, where RNA modulates RBP localization, conformation, interaction and function (4–8).

Due to their central role in various key cellular processes, dysregulation of RBP functions is implicated in the initiation and development of diseases such as neurological disorders, cancer and muscular atrophies (3,9,10). Therefore, RBPs have attracted increased interest in recent years, leading to the development of multiple strategies to establish comprehensive catalogs of RBPs in humans and in other species (11,12). Proteome-wide approaches include the identification of the mRNA-bound proteome via UV-crosslinking and subsequent oligo(dT) capture (13–18). Other techniques based on protease digestion (19,20), modified nucleotides (21,22) or organic phase separation (23) have been established to include RBPs that also bind to non-polyadenylated RNAs and orthogonal approaches include bioinformatic-based studies (24,25). R-DeeP is a complementary experimental strategy based on a sucrose density gradient and ultracentrifugation and thus independent of affinity or property-based protocols (26–28). R-DeeP adapts the concept of RNA dependence with RNA-dependent proteins being defined as proteins whose interactome depends on RNA and thus comprising proteins interacting directly or indirectly with RNA molecules. Using the R-DeeP strategy, more than 700 new RNA-dependent proteins have been recently identified in human cancer cells (28) and 545 in *Plasmodium falciparum* (29).

All these valuable resources support the view that canonical RBPs can bind to RNA through their structurally well-defined RNA-binding domains (RBDs) such as the RNA recognition motif (RRM) or the K-homology domain (KH) (12,30). However, they also point to a large number of unconventional RBPs that do not contain any canonical RBDs (5,31,32). Interestingly, the non-canonical or unconventional RBPs represent the large majority of RBPs detected in humans (76.2% of the RBPs) and other species (12). Unconventional RBPs usually perform primary functions in a great variety of key cellular pathways which are unrelated to RNA metabolism - and are thus prime candidates for proteins which do not regulate RNA, but are regulated by RNA interactions. Currently, their affinity for RNA and possible physiological relevance remains mostly uncharacterized. A few recent examples include the RNA-dependent stimulation of the transcription co-activator CBP to promote histone acetylation and regulation of enhancer function (33), the RNA-dependent differential regulation of the interferon-inducible protein kinase R (PKR) (6), the importance of the RNA-binding activity of ERα for tumor cell survival and therapeutic response (34) and the riboregulation of the glycolytic enzyme enolase 1, which alters cell metabolism and stem cell differentiation (7).

In the context of cell division, several studies have reported the presence of RNA, *i.e.* protein-coding (mRNAs) (35–37), non-coding RNAs (ncRNAs) (38,39) and ribosomal RNAs (rRNAs) (37,40), as well as various RBPs (41,42) within structures of the mitotic spindle apparatus such as the centrosomes (43,44), the kinetochores (44) and the spindle microtubules (36,37). Collectively, these works point to the possible importance of RNA and RBPs for the structural and functional integrity of the mitotic spindle. However, although RNase treatment (41,45) and inhibition of splicing and transcription (38) can disrupt the mitotic spindle apparatus in the *Xenopus laevis* egg extract system, the underlying mechanisms remain unknown. Therefore, we aim here to investigate RNA-dependent proteins involved in mitosis and how they might be affected by RNA.

The available proteome-wide RBP screens provide a highly valuable resource to further investigate the role and relevance of RBPs and RNA in key cellular processes. However, due to the low number of mitotic cells in non-synchronized cell populations, the differential protein expression levels throughout the cell cycle and the dynamics of post-translational modifications that may affect the affinity of proteins for RNA, these screens so far are not adapted to adequately reflect this important phase of the cell cycle. Therefore, we applied the R-DeeP screening strategy (26,27) in both mitotic and interphasic synchronized HeLa cells, to collect specific and quantitative data on RNA-dependent proteins in both cell cycle phases. This resource is now available online in the R-DeeP 3.0 database (https://R-DeeP3.dkfz.de).

Taking advantage of this new resource, we provide an atlas of RNA-dependent proteins in cell division, unraveling that a large number of well-known mitotic factors and key cell division players are linked to RNA, some also supported by the multiple studies that were integrated into the RBP2GO database (12). In particular, we identified and characterized the RNA dependence of a major mitotic regulator, Aurora kinase A (AURKA). We reveal that AURKA directly binds to RNA and that the presence of RNA is essential for the interaction of AURKA with multiple proteins, including several known targets such as TPX2 and a new target, the kinesin-like protein KIFC1 (KIFC1, also known as XCTK2 or HSET, from the Kinesin-14 family proteins). We show that AURKA phosphorylates KIFC1 at S^349^ and T^359^. Our data demonstrate that KIFC1, AURKA and TPX2 are unconventional RBPs, *i.e.*, they directly bind to RNA, without containing a known RNA-binding domain. The detailed analysis of the KIFC1-bound RNA reveals that KIFC1 interacts with both rRNA and protein-coding transcripts without apparent sequence specificity.

Altogether, our findings reveal an essential role for RNA in mediating RNA-dependent mitotic protein-protein interactions. Thus, our atlas of RNA-dependent proteins in cell division promises the exciting development of a research field integrating RNA-protein interactions as a new layer of regulation in our understanding of cell division processes.

## RESULTS

### Differential R-DeeP screens in mitosis and interphase provide cell cycle-specific data on RNA-dependent proteins

A gene ontology (GO) analysis of the RNA-dependent proteins identified in unsynchronized HeLa cells (26,27) revealed the significant enrichment of proteins with GO terms related to mitosis (Supplementary Figure S1). However, mitotic cells are underrepresented in an unsynchronized cell population. Therefore, to obtain a more comprehensive landscape of the RNA-dependent proteins in mitosis, we established the R-DeeP approach in HeLa cells synchronized in mitosis and in interphase (Figure 1A).

**Figure 1.**
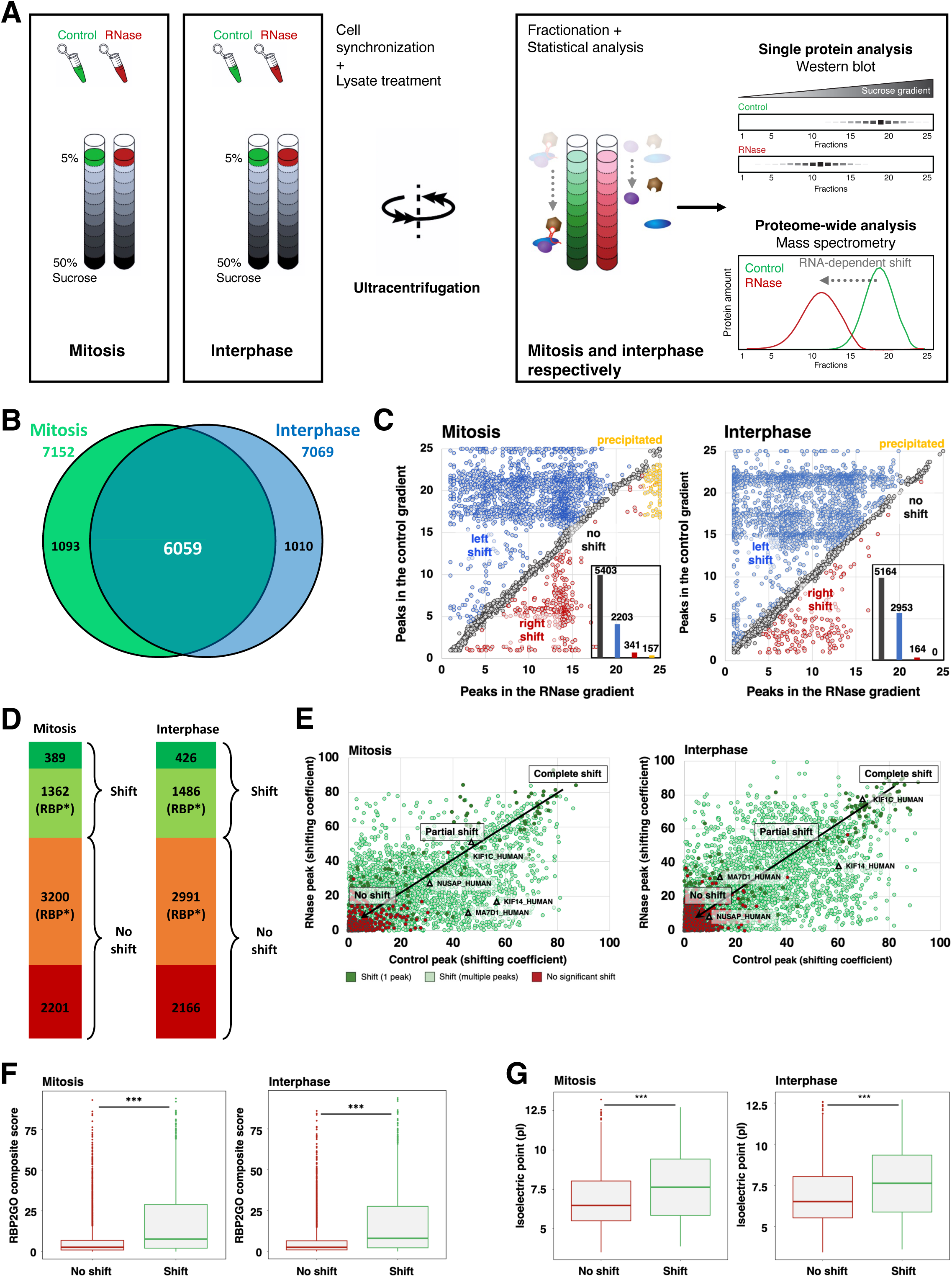
R-DeeP in HeLa mitotic and interphasic cells. **A**: HeLa mitotic or interphasic untreated (control) or RNase-treated cell lysates were prepared and loaded onto sucrose density gradients (5%–50%). Following ultracentrifugation, the gradients were fractionated into 25 fractions and subjected to either mass spectrometry (proteome-wide screen) or western blot analysis (individual protein analysis) for validation of the screens (N=3 in total). Adapted from (26). **B**: Venn diagram indicating the total number of proteins identified in each R-DeeP screen (mitosis: 7152 and interphase: 7069). 6059 proteins were commonly identified. **C**: Each graph (mitosis and interphase, respectively) indicates the position of maxima for each shift in the control and RNase-treated gradients according to the mean of three replicates. The inset bar graph shows the number of proteins in each sub-category (left shift: blue, right shift: red, no shift: gray and precipitated proteins: orange). **D**: Classification of the proteins analyzed in the R-DeeP mitotic and interphasic screens according to their shifting behavior and their prior classification in 43 proteome-wide human studies. RNA-dependent proteins not linked to RNA before - in mitosis screen: 389 and interphase screen: 426. RBP*: validated RBPs or RBP candidates. **E**: Graph depicting the shifting co-efficient of the proteins for each pair of control and RNase peaks. Red: No significant shift, dark green: significant shift from one control peak to one RNase peak, light green: shifts between multiple peaks. For shifts between multiple peaks, they are further categorized as complete shifts: total amount of protein shifting between control and RNase gradients (top right of the graph), partial shift: a of protein shifting between control and RNase gradients (middle) and no shift: no change in shifting pattern (bottom left of the graph). Representative microtubule-related proteins are highlighted on the graph. **F**: Boxplots representing the distribution of the RBP2GO composite score of the proteins classified as non-shifting (No shift) or shifting (Shift). The bar and the box indicate the lower, the median and the upper quartiles. The whiskers represent the range between the bottom of the first quartile and the top of the third quartile, excluding the outliers. The outliers are represented by dots and the p-value was calculated using a Wilcoxon test (*** *P*-value < 0.001). **G**: Boxplots representing the isoelectric point (pI) of the proteins classified as non-shifting (No shift) or shifting (Shift). The bar and the box indicate the lower, the median and the upper quartiles. The whiskers represent the range between the bottom of the first quartile and the top of the third quartile, excluding the outliers. The outliers are represented by dots and the p-value was calculated using a Wilcoxon test (*** *P*-value < 0.001).

The migration pattern of the proteins, *i.e.*, their distribution across the different fractions in the control as compared to the RNase-treated gradients revealed the RNA dependence or independence of the proteins (Figure 1A). We quantified a total of 7152 proteins in the mitotic gradient and 7060 proteins in the interphasic gradient, which included 6059 proteins that were commonly found in both screens (Figure 1B).

Using the established bioinformatic pipeline to automatically identify RNA-dependent proteins (26,27), Gaussian-fitted distributions were calculated for each protein profile, and shifts were characterized based on (i) amount of protein shifting in control vs. RNase-treated gradients: indicated by area under the curve, (ii) position of the peaks representing the apparent molecular weight of the complex, (iii) direction and distance of the shift, (iv) amplitude difference between control and RNase curves at each maxima and (v) statistical significance in the difference between amplitude maxima in the control and RNase-treated gradients. A protein was classified as RNA-dependent if the distance between the maxima in the control and RNase-treated gradients was at least greater than one fraction with statistically significant difference between the two gradients for at least one maximum. Next, the proteins were classified according to their shift and direction as follows: (i) left shifting proteins: proteins whose maxima were detected in lower fractions in RNase-treated gradients compared to control gradients indicating a lower apparent molecular weight upon RNase treatment, (ii) right shifting proteins: proteins whose maxima were detected in later fractions in RNase-treated gradients compared to control gradients, (iii) precipitated proteins: proteins shifting to greater fractions (fraction > 23) upon RNase treatment and (iv) non-shifting proteins: proteins whose maxima were detected in the same fraction in RNase-treated gradients compared to control gradients (Figure 1C and Supplementary Figure 2A). As commonly observed (26,28,29), RNA-dependent proteins were dominated by left-shifted proteins, *i.e.*, proteins losing interaction partners upon RNase treatment.

Out of the 7152 detected proteins in the mitosis screen, we identified 1751 proteins with at least one shift. Similarly, 1912 out of the 7060 proteins depicted at least one shift in the interphase screen (Supplementary Table S1). 948 common left-shifting proteins were identified in mitotic and interphasic screens, out of which 160 proteins were more highly expressed in mitosis and 52 proteins were more highly expressed in interphase (Supplementary Figure 2B and Supplementary Table S1). In addition to the 160 left-shifting proteins which were higher expressed in mitosis, our analysis revealed 500 left-shifting proteins detected only in mitosis. Expectedly, a GO enrichment analysis detected several terms related to microtubule and mitotic spindle activities (Supplementary Table S1). Importantly, we identified 389 and 426 RNA-dependent proteins in mitosis and interphase, respectively, (Figure 1D), which had never been linked to RNA before according to the RBP2GO database (12).

Taking advantage of the quantitative aspect of the R-DeeP screen, a shifting coefficient was calculated according to the amount of protein in each peak of the control and RNase-treated gradients (26). Based on the shifting coefficient, the proteins were further categorized as (i) partially RNA dependent (partial shift, *i.e.* only part of the protein amount shifts), (ii) completely RNA dependent (complete shift, *i.e.* the entire amount of the protein shifts) and (iii) RNA independent (no shift) (Figure 1E). To validate the RNA dependence of the newly identified RNA-dependent proteins, analyses on multiple parameters related to RBPs were performed by comparing the groups of shifting vs non-shifting proteins. For example, the recently defined RBP2GO composite score, which reflects the probability for a protein to be a true RBP (12), was consistently and significantly higher for shifting proteins as compared to non-shifting proteins in both mitotic and interphasic screens (Figure 1F). In addition, as expected from previous studies (14,24,26), mitotic and interphasic shifting proteins depicted higher isoelectric points (pI) than non-shifting proteins (Figure 1G).

In order to provide an open and easy access to the R-DeeP screen data, we compiled the R-DeeP 3.0 database (https://R-DeeP3.dkfz.de). R-DeeP 3.0 allows users to conveniently browse the present mitotic and interphasic R-DeeP screens, as well as the previously published screens that had been performed in unsynchronized HeLa (26) and A549 cells (28) to facilitate comparison of the results (Supplementary Figure S2C). The database provides various search options such as the single or advanced search option using gene name or UniProt ID to either obtain a summary of a single protein or to compare a protein between different cell lines and cell cycle stages. These multiple analysis options provide various output formats which can be downloaded for further custom analyses. Detailed instructions on how to use the database are provided via a comprehensive user guide, which can be accessed from the documentation section.

### An atlas of RNA-dependent proteins in cell division highlights well-known mitotic factors as RNA-dependent proteins

Mitosis is critical for the faithful segregation of the chromosomes to the daughter cells during cell division. This is mediated by a complex and dynamic macromolecular machinery that consists of polymerized microtubules, centrosomes and various specialized proteins (41). So far, most of the studies focused on the functional characterization of proteins and protein-protein interactions (46–48). Only a few studies have focused on the role of RNA in mitosis (38,40–42). The role of RNA at centrosomes has been discussed in the past (49) but not further explored. In sum, how RNA might be functionally involved during mitosis still requires further investigations. Our present R-DeeP screen in mitosis depicted an enrichment for proteins with microtubule-related functions (Supplementary Table S1). Taking advantage of these data together with the comprehensive knowledge from the RBP2GO database (11,12), we created a list of RNA-dependent proteins in mitosis. More specifically, combining our screening data with the use of the advanced search option from the RBP2GO database along with GO terms related to mitosis, we generated an atlas of 826 RNA-dependent proteins from of a total of 1472 mitotic proteins, meaning that 56.1% of the mitotic proteins were related to RNA (Figure 2A and Supplementary Table S2). 643 of these proteins (77.8%) did not contain any known RBD and were not linked to an RNA-related InterPro family ID (12). The information on the localization of these RBPs in various critical mitotic structures such as spindle poles, spindle midzone, kinetochores and chromosomes were included in the map, showing that nearly all mitotic substructures contained RNA-dependent proteins. Many of these proteins have well-known and characterized functions during cell division, *e.g.*, AURKA or TPX2, which are unrelated to RNA. Interestingly, the vast majority belong to the class of unconventional RBPs, awaiting further RNA-related functional characterization. Additionally, we categorized RNA-dependent proteins that are crucial for cell cycle transitions (Figure 2B). We also integrated the protein lists from the MiCroKITS database into our analysis (50), to provide orthogonal information on spindle, centrosome, kinetochore and midbody proteins and confirmed that >50% of the proteins from these lists were classified as RNA-dependent proteins, as well (Supplementary Table S2). Altogether, our atlas of RNA-dependent proteins in cell division highlights the extensive incidence of RNA-dependent proteins in cell division structures and events, including key mitotic factors, which strongly suggests RNA as a crucial co-factor.

**Figure 2.**
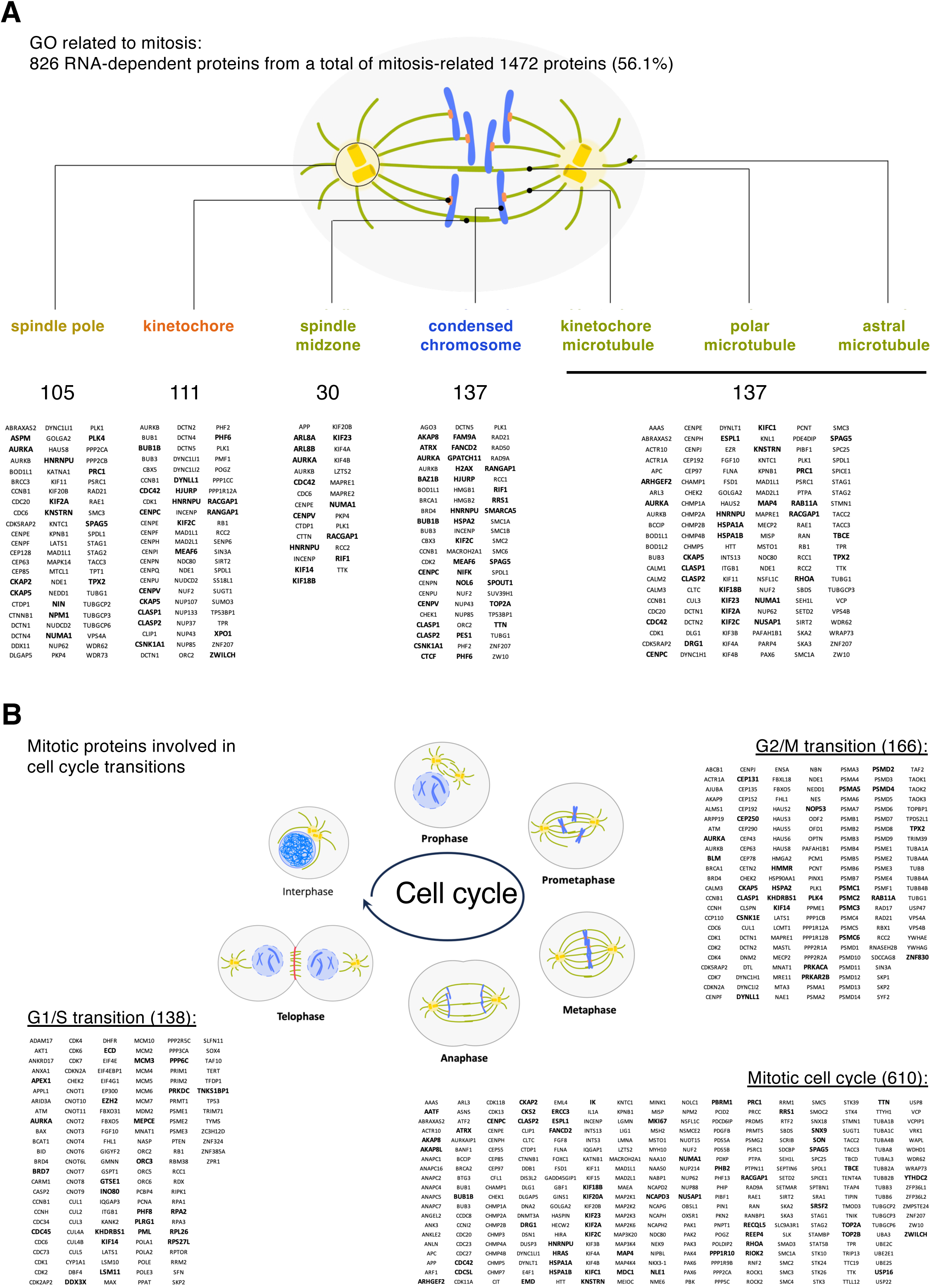
An atlas of RNA-dependent proteins in cell division. **A**: Schematic representation of a mitotic spindle showing the localization of 826 RNA-dependent proteins (out of 1472 proteins in total) to mitotic structures, based on their association with the respective GO terms. Data from the 43 proteome-wide human RBP screens as compiled in the RBP2GO database were used. The numbers indicate the number of proteins in each group. Only a random subset of the proteins are listed in this figure. The complete dataset can be found in Supplementary Table S2. **B**: Same as in A for mitotic proteins involved in different cell cycle transitions. Proteins indicated in bold represent the shifting proteins as identified in our present R-DeeP screen in mitosis. Their complete mitotic profile is available within the R-DeeP3 database (R-DeeP3.dkfz.de).

### Aurora Kinase A is an RNA-dependent protein

Since AURKA is an essential kinase for cell cycle regulation, we further investigated its RNA dependence. AURKA belongs to the highly-conserved serine/threonine kinase family and is essentially activated by autophosphorylation at the T^288^ amino acid residue (51). It is a cell cycle-regulated protein whose expression level increases from late S-phase and is degraded during mitotic exit (52). AURKA is first activated in late G2, where the protein interacts with PLK1 and CEP192 for centrosome maturation (48,53). Upon nuclear envelope breakdown and onset of a Ran-GTP gradient from the chromosomes, AURKA interacts with TPX2 (46,47). AURKA and TPX2 activate each other and interact with several other proteins such as NEDD1, RHAMM, and γ-Tubulin Ring Complexes to promote microtubule nucleation, spindle focusing and bundling (48,54). AURKA serves also as a therapeutic cancer target that is often overexpressed in several cancers (54,55).

In our present mitotic R-DeeP screen, the mass spectrometry analysis depicted a complete left shift in the AURKA profile upon RNase treatments from the control fraction 21 to the RNase fraction 5 (Figure 3A). The RNA-dependent shift of AURKA in mitosis was validated per Western blot analysis. The quantification graph of the Western blot bands showed a similar left shifting pattern as seen in the mass spectrometry quantification, strongly validating the RNA dependence of AURKA in mitosis and suggesting a loss of interaction partners upon RNase treatment (Figure 3B and 3C).

**Figure 3.**
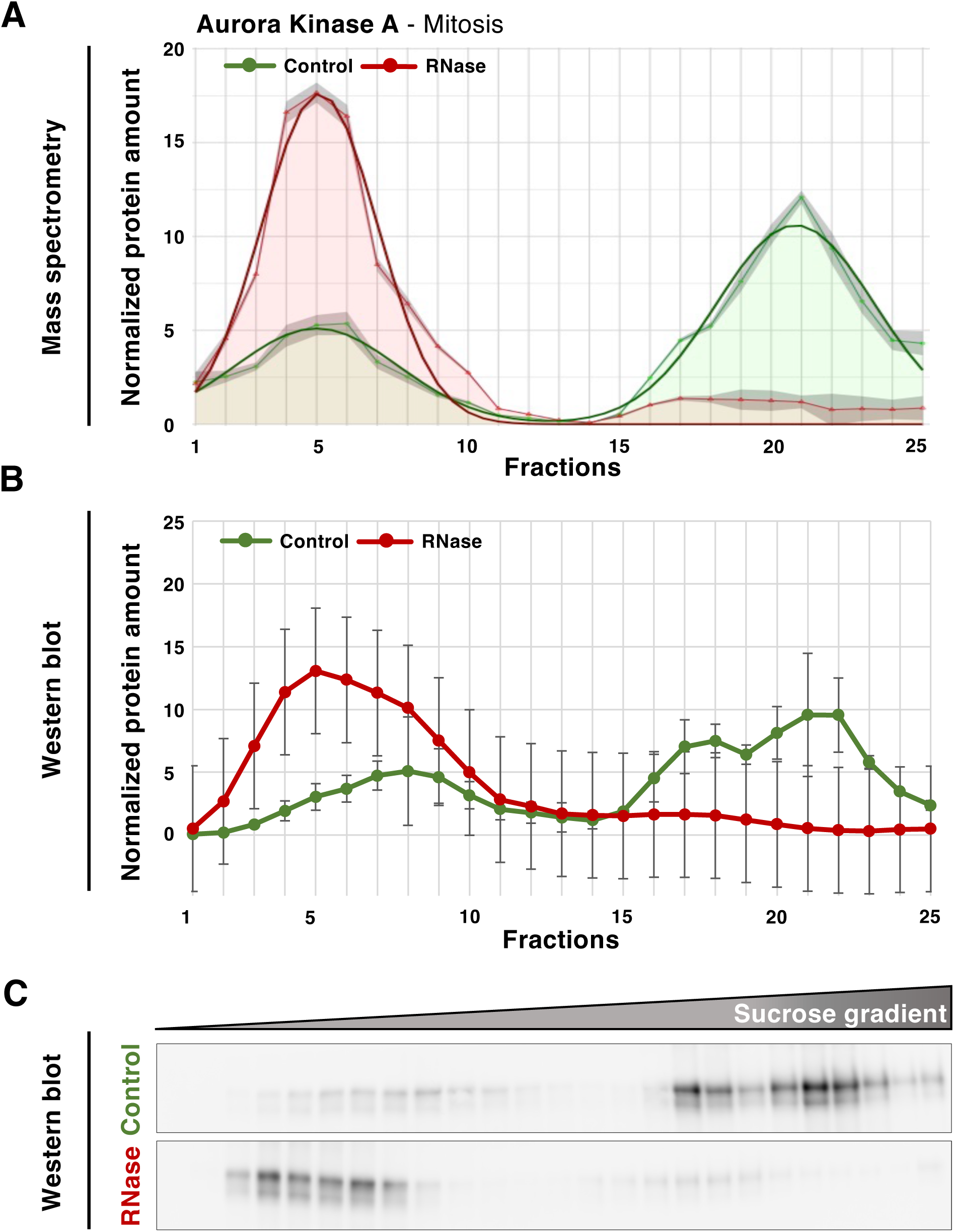
AURKA is an RNA-dependent protein in mitosis. **A**: Graphical representation of the protein amount in 25 different fractions of control (green) and RNase-treated (red) sucrose density gradients analyzed by mass spectrometry. Raw data (mean of three replicates) are depicted by line with markers. Smooth lines represent the respective Gaussian fit. The overall protein amount of the raw data was normalized to 100. **B**: Graph showing the quantitative analysis of western blot (WB) replicates depicted by the mean of three replicates with standard error of the mean (SEM, N=3). **C**: Western blot representing the distribution of AURKA in 25 different fractions in control and RNase-treated mitotic gradients.

### AURKA has both RNase-sensitive and RNase-insensitive interactions with other proteins

Protein-protein interactions of AURKA as a key mitotic factor were extensively studied, largely based on proteomic identification of its mitotic substrates (56,57). The loss of interaction partners upon RNase treatment motivated us to further investigate the RNA-dependent protein interactors of AURKA in mitosis. Therefore, we performed immunoprecipitation of AURKA in mitotic HeLa cell lysates, in the presence or absence of RNase treatment, followed by mass spectrometry analysis. Among the identified AURKA interacting proteins, 90.4% (1080 out of 1194) of the proteins were RNA-dependent according to the RBP2GO classification and our new mitotic R-DeeP screen (Figure 4A and Supplementary Table S3). We observed that some of the AURKA interactors including for example centrosomal proteins were not sensitive to RNase as their interaction with AURKA persisted after RNase treatment. Interestingly, well-known AURKA mitotic interactors such as TPX2, HMMR and CLASP1 interacted with AURKA in an RNase-sensitive manner (Figure 4B and Supplementary Table S3). We also identified KIFC1 as one of the most frequently detected RNase-sensitive candidates, whose interaction with AURKA had not been characterized before. KIFC1 is a minus end directed motor protein localized to the nucleus due to the presence of a nuclear localization signal (NLS) in its tail domain (58). Similar to TPX2, KIFC1 is bound by importins but released in a RanGTP-dependent manner to then function in spindle organization, microtubule focusing and crosslinking (59–61). In the following, we will focus on validating the RNA-dependent interactions of AURKA with TPX2 and the new interactor KIFC1.

**Figure 4.**
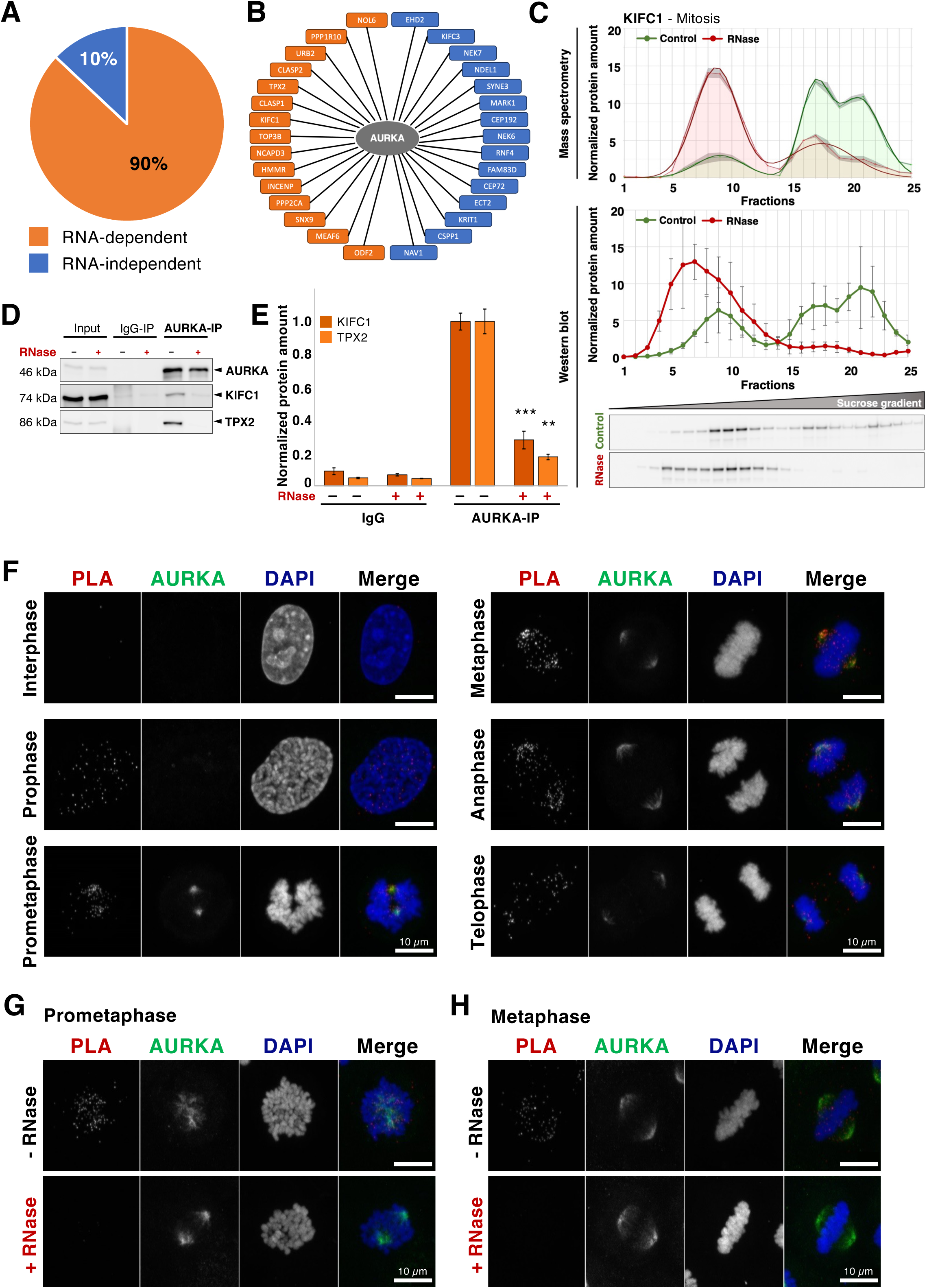
RNA-dependent protein interactors of AURKA. **A**: Pie chart showing the percentage of AURKA protein interactors that are RNA-dependent (orange) or - independent (blue), according to RBP2GO and our new mitotic R-DeeP screen (12), and identified by AURKA pulldown and mass spectrometry analysis in HeLa cells synchronized in mitosis (N=3). A complete list is available in Supplementary Table S3. **B**: Selected examples of AURKA RNase-sensitive mitotic factors (orange) and RNase-insensitive protein interactors (blue), as detected from the AURKA pulldown in presence or absence of RNase treatment. A complete list is available in Supplementary Table S3. **C**: R-DeeP profile of KIFC1 in mitosis and WB validation as for AURKA (see Figure 3). **D**: Western blot (WB) analysis showing the immunoprecipitation of AURKA in mitotic HeLa cells. AURKA pulldown was performed in the mitotic lysate treated with or without RNase I. Rabbit IgG was used as a negative control. KIFC1 (74 kDa) and TPX2 (86 kDa) were pulled down with AURKA (46 kDa) in control samples whereas their interaction was significantly reduced upon RNase treatment (reduction of the band intensity for KIFC1 and TPX2 in the last lane). **E**: Graph representing the amount of KIFC1 and TPX2 present in IgG and AURKA pulldown samples treated with or without RNase I. The intensity of the WB bands were quantified using Image J and represented in the bar graph with SEM (N=3). *P*-values were evaluated using a two-tailed, paired t-test (** *P*-value < 0.01, *** *P*-value < 0.001). **F**: Representative proximity ligation assay (PLA) images indicating the close proximity of AURKA and KIFC1 in HeLa cells across the cell cycle (interphase to telophase as indicated). The interaction is represented by dots (PLA channel, red dots in the merge channel), AURKA was stained per immunofluorescence (green) and DNA was stained using DAPI (blue). Controls of the PLA assay and quantifications are seen in Supplementary Figure S4 (part 1). Scale bar, 10 µm. **G** and **H**: PLA representative images showing the proximity of AURKA and KIFC1 in HeLa cells at prometaphase and metaphase, respectively, (PLA channel, red dots in the merge channel). AURKA is seen in green (immunostaining) and DNA was stained using DAPI (blue). The top images depict representative images in control cells. Bottom images depict representative images in RNase-treated cells. Scale bar, 10 µm.

The data from our mitotic R-DeeP analysis confirmed that KIFC1 and TPX2 were RNA-dependent proteins (Figure 4C, Supplementary Figure S3A). Similar to AURKA and TPX2, KIFC1 was differentially expressed throughout the cell cycle and highly expressed during mitosis (Supplementary Figure S3B-S3C, see R-DeeP3.dkfz.de for the comparative protein profiles in mitosis versus interphase, normalized to the protein amount in mitosis). Furthermore, AURKA, KIFC1 and TPX2 shifted from similar fractions in the control gradient with protein amount maxima at around fraction 21, indicating that these three proteins could be part of the same RNA-dependent protein complex (Figure 4C, Supplementary Figure 3A).

The mass spectrometry results of the AURKA pulldown were first validated using immunoprecipitation in presence and absence of RNase, followed by Western blot analysis. Upon RNase treatment and AURKA pulldown, the interaction with TPX2 and KIFC1 was significantly lost in both, HeLa and A549 mitotic cells (Figure 4D-4E, Supplementary Figure S3D-S3E). We also tested for the interaction of TPX2 with KIFC1, as our data indicated that they could be part of the same complex. Our TPX2 pulldown revealed the direct interaction between TPX2 and KIFC1. Additionally, we observed a decrease of the interaction of TPX2 with KIFC1 and AURKA upon RNase treatment and TPX2 pulldown in mitotic lysates (Supplementary Figure S3F-S3G). These observations indicated that the three proteins interacted with each other in an RNA-dependent manner and possibly function as a complex in mitosis.

To visualize the proximity of the proteins in the native cellular environment, we performed a proximity ligation assay (PLA). This technique supports the visualization of *in situ* interactions between two proteins, at endogenous levels, represented by signal in the form of dots resulting from a ligation followed by an amplification reaction (62). First, we monitored the interaction between AURKA, KIFC1 and TPX2 throughout mitosis. All tested interaction pairs, *i.e.* AURKA-KIFC1 (Figure 4F), TPX2-AURKA (Supplementary Figure S4A) and TPX2-KIFC1 (Supplementary Figure S4B) depicted strong PLA signals from prophase (within the centrosomal asters) through metaphase to telophase (at the spindle poles). This data indicated a colocalization between AURKA, KIFC1 and TPX2 during spindle assembly.

Furthermore, we repeated the PLA in presence or absence of RNase treatment. The resulting signal was quantified using an ImageJ-based analysis (Supplementary Figure S5A). We observed a clear PLA signal in the control cells in prometaphase and metaphase around the centrosomal asters and the spindles respectively, indicating the proximity of the proteins. In contrast, the PLA signal strongly decreased upon RNase treatment, reaching a level comparable to the background level produced by each antibody individually (Figure 4G-4H, Supplementary Figure S5B-S5E and Supplementary Figure S6A-S6H). Altogether, these observations further validated that AURKA, TPX2 and KIFC1 interactions were mediated by RNA.

In order to confirm that the loss of interaction between the proteins was due to the lack of RNA, and not due to the adverse effects of RNase itself, the interaction between α- and β-tubulin was used as a negative control. α-tubulin and β-tubulin are the constituting subunits of the microtubules. Thus, we reasoned that the α/β-tubulin interaction should persist as long as microtubules are present in the cell. The overall PLA signal between the control and the RNase-treated samples remained at a high level, confirming the persistence of the α/β-tubulin interaction upon RNase treatment (Supplementary Figure S7A-S7B). However, we noticed a modification of the α/β-tubulin PLA signal distribution, which seemed less homogeneously distributed after RNase treatment. To quantify the distribution of the α/β-tubulin PLA signal, the collected images were rotated and rescaled to finally be superposed, so that the variance of the signal could be evaluated pixel per pixel (Supplementary Figure S7C). This analysis revealed an increased variance of the α/β-tubulin PLA signal throughout the metaphasic spindle structure after exposure to RNase, in particular at the poles of the structure (dark orange, Supplementary Figure S7C). In addition to the PLA analysis, immunoprecipitation of β-tubulin was performed in mitotic lysates from cells that stably expressed GFP-α-tubulin. The Western blot analysis also confirmed the persistence of the interaction between β- and α-tubulin after RNase treatment (Supplementary Figure S7D-S7E).

Altogether, we demonstrated that both AURKA and TPX2 interacted with KIFC1 throughout mitosis and that the interactions between AURKA, KIFC1 and TPX2 were RNA-dependent. While α/β-tubulin interaction persisted upon RNase treatment, the distribution of the PLA signal became less homogeneous, indicating the introduction of some perturbations in the microtubule-based structure. Thus, our results indicated that RNA differentially regulates protein-protein interactions during mitosis.

### KIFC1-interacting RNAs do not contain sequence-specific binding sites

After demonstrating that the interactions between the AURKA, KIFC1 and TPX2 were RNA dependent, we performed an individual-nucleotide resolution UV cross-linking and immunoprecipitation (iCLIP) assay (63) to determine whether the proteins interacted directly with RNA transcripts. Under conditions that were adapted to each protein, we observed their direct interaction with RNA in HeLa and A549 cells synchronized in mitosis. Upon increasing dilution of the RNase 1 (1:5 to 1:1000 depending on the protein), the RNA signal migrated to higher molecular weight, corresponding to protein-RNA complexes with increasing ranges of transcript length (Figure 5A-5B and Supplementary Figure S8A-S8C).

**Figure 5.**
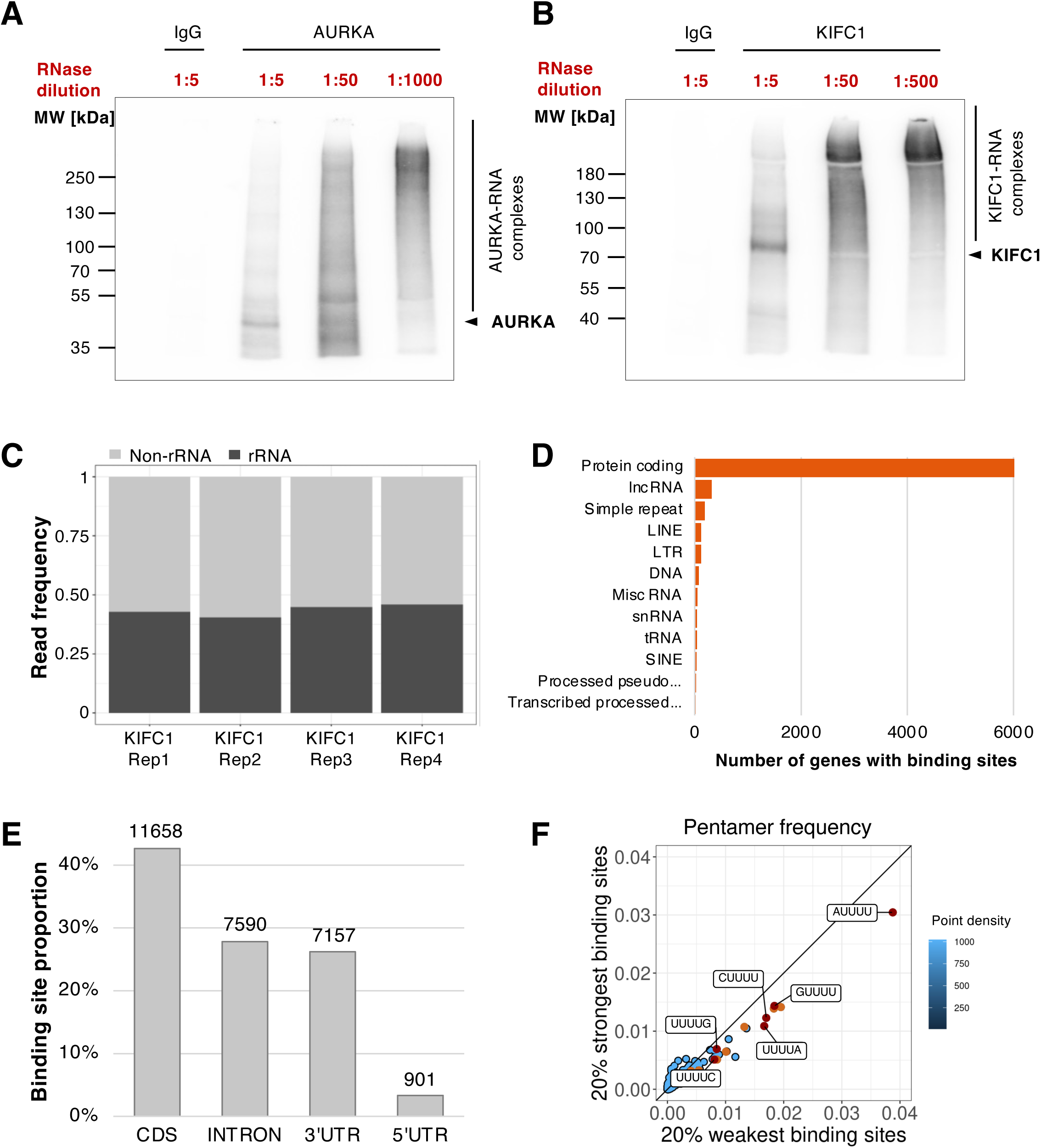
RNA mediating the interaction of AURKA, KIFC1 and TPX2. **A** and **B**: Autoradiography indicating the direct binding of AURKA and KIFC1, respectively, to RNA by iCLIP2 indicated by shifting of the radioactive RNA signal towards higher molecular weights with decreasing RNase I concentrations in HeLa prometaphase cells (representative images out of N=3 replicates are shown). **C**: Bar plot depicting the ribosomal RNA (rRNA) and non-ribosomal RNA (non-rRNA) read frequencies in individual KIFC1 iCLIP2-Seq replicates in HeLa prometaphase cells. **D**: Horizontal bar plot showing the KIFC1 target non-rRNA gene spectrum with number of genes identified in iCLIP2 in HeLa prometaphase cells in decreasing order. The genes are classified as protein coding, lncRNA, simple repeat, LINE, LTR, DNA, Misc RNA, snRNA, tRNA, SINE, processed pseudogene and transcribed processed pseudogene. **E**: Bar plot representing the proportion of binding sites in the respective transcript regions of protein coding genes. **F**: Scatter plot comparing the pentamer frequency within the 7-nt binding sites in the 20% strong binding sites vs. 20% weakest binding sites as defined by the PureCLIP score. The pentamers with the most extreme frequencies are colored in orange and red and contain U-stretches.

To further identify the interacting RNAs, we performed an iCLIP2 (63,64) for KIFC1, which presented a high radioactive signal of interacting RNAs (Figure 5B). About 80 million reads were obtained from each of the four replicates, which correlated well with each other (Supplementary Figure S8D). Following quality control and alignment, we noticed that up to 45% of the reads represented rRNA (Figure 5C). Furthermore, the analysis of the uniquely mapped reads showed that the vast majority of the binding sites (∼80%, 29465 out of 36934) were associated to protein-coding mRNA transcripts (Figure 5D, Supplementary Figure S8E and Supplementary Table S4). In total, KIFC1-associated binding sites were localized in 5687 genes (Supplementary Table S4), in particular in the coding, intronic and 3’UTR sequences (Figure 5E). The analysis of the pentamer frequency at the binding sites did not reveal a particular sequence specificity for the strong binding sites, which were defined according to the PureCLIP scoring system (65). The most enriched pentamers contained U-stretches, most likely reflecting the uridine bias of UV crosslinking (66), and were rather localized in the lower 20% binding sites (Figure 5D). In summary, RNA sequences binding to KIFC1 were predominantly originating from rRNA and protein-coding genes, and apparently lacking sequence specificity.

### AURKA phosphorylates KIFC1 at S^349^ and T^359^

Next, we reasoned that AURKA is one of the major mitotic kinases, whose main function is to phosphorylate several mitotic factors for their effective activation and promotion of cell division (48). Since the interaction between KIFC1 and AURKA was not characterized, we tested the hypothesis that KIFC1 could be a substrate for AURKA, as suggested by earlier large-scale quantitative phosphoproteomic analyses (57). Using the suggested consensus sequence of AURKA (57,67), we selected eight serine and threonine residues that were distributed across the KIFC1 sequence and localized in various domains such as disordered regions and the kinesin motor domain (Figure 6A-6B). Using KIFC1 pulldown from HeLa cell lysates overexpressing KIFC1 wild-type (WT) or KIFC1 mutants with non-phosphorylatable alanine, we performed an *in vitro* kinase assay, in the presence or absence of purified AURKA. The assay revealed two interesting residues in KIFC1: S^349^ and T^359^, whose mutation to alanine induced a strong reduction of phosphorylation by AURKA (Supplementary Figure S9A-S9B). Confirming these results, a mutant version of KIFC1, containing both mutations (S349A and T359A) depicted a >90% loss of phosphorylation as compared to the KIFC1 WT protein (Figure 6C-6D). A kinase-dead AURKA D274A confirmed the specificity of the obtained phosphorylation of KIFC1 b AURKA. Altogether, our results show that AURKA and KIFC1 interact in an RNA-dependent manner and that AURKA phosphorylates KIFC1 at residues S^349^ and T^359^.

**Figure 6.**
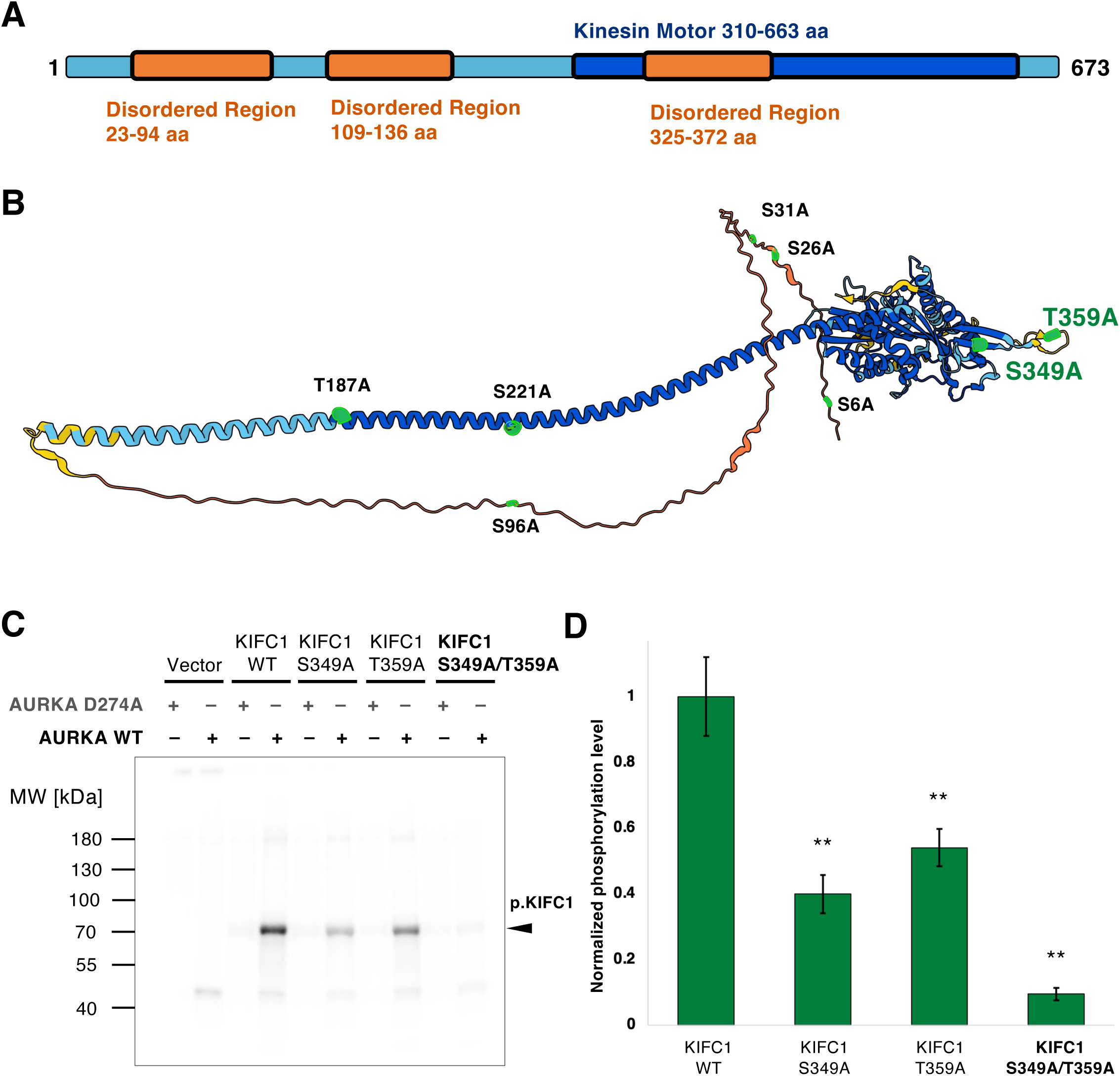
AURKA phosphorylates KIFC1 at S^349^ and T^359^ amino acid residues. **A**: Schematic representation of KIFC1 indicating the location of the kinase domain and the position of disordered regions across the KIFC1 structure. **B**: Schematic representation of KIFC1 indicating the position of eight potential phosphorylation sites (highlighted in green) that were identified based on the consensus sequence of AURKA (27). The schematic was generated using AlphaFold (alphafold.ebi.ac.uk). **C**: Autoradiography indicating the phosphorylation intensity of KIFC1 in wild-type (WT) and non-phosphorylatable KIFC1 mutants in the presence of purified AURKA WT or AURKA kinase-dead mutant (D274A). The *in vitro* kinase assay was performed using KIFC1 pulled down from HeLa prometaphase lysates overexpressing the WT or mutant KIFC1 proteins with an N-terminal Flag-HA tag. An empty vector was used as a negative control. **D**: Quantification of the autoradiography image (as in **C**), representing the KIFC1 phosphorylation signal in the form of a bar graph with SEM (N=3). *P*-values were calculated using two-tailed, paired t-test (** *P*-value < 0.01).

## DISCUSSION

The field of RNA biology is exposed to a number of exciting new challenges linked to the discovery of an increasing number of RBPs (11,13–23). On the one hand, unconventional RBPs, which interact with RNA in absence of canonical RBDs, represent a large majority of the RBPs in many species (12). Usually, the cellular functions of these unconventional RBPs are well-characterized, but their interaction with RNA was overlooked and thus remained for most of them unexplored. On the other hand, in a traditional view, RNA-protein interactions and dynamic RNP assemblies are mainly seen as a mean for proteins to control the fate of cellular RNAs, from transcription to degradation. However, this view is being challenged by the expanding concept of riboregulation, according to which RNA transcripts bind to RBPs and regulate their localization, conformation, interactions and function (5,7,8,68).

Here, we took advantage of the R-DeeP strategy to investigate RNA-dependent proteins in a cell cycle-dependent manner. A protein is termed “RNA dependent” if its interactome is dependent on the presence of RNA. We identified 7152 and 7069 RNA-dependent proteins in mitosis and interphase, respectively, including hundreds of proteins that had not been previously related to RNA. Our screens highlighted both similarities and differential behaviors of the proteins in mitosis and interphase, which can be explored within the R-DeeP 3.0 database (R-DeeP3.dkfz.de), as well as compared to previous datasets from R-DeeP screens in unsynchronized HeLa (26) and A549 cells (28). In addition, by integrating these results with the data from the RBP2GO database (11), we compiled an atlas of RNA-dependent proteins in cell division. The atlas reveals the localization of RBPs in various mitotic structures and their association with cell cycle transition phases. For example, our atlas reports a great number of midbody-associated proteins as RNA-dependent proteins, in agreement with a recent work describing midbodies as assembly sites of RNP granules (69). It also comprises a substantial number of centrosome-associated proteins, in line with previous debated studies reporting the presence and functional importance of RNA at the centrosomes. The potential role of RNA at the centrosome has been discussed (49) but not further experimentally studied. In addition, our atlas highlights RNA-dependent proteins that were identified in the mitotic R-DeeP screen whose quantitative gradient centrifugation profiles can be downloaded from the R-DeeP 3.0 database. Such an atlas could be easily adapted to any other cellular processes of interest, using selected GO terms in the advanced search option of the RBP2GO database (RBP2GO.dkfz.de). Altogether, this illustrates one of the utilities of integrating proteome-wide datasets into a central database, and to develop targeted tools that increase the functionality of the database as much as the impact of the included studies and datasets.

Over the past decades, we have been accustomed to the concept that a protein is regulated in terms of localization and functions through post-translational modifications that are mediated through protein-protein interactions. However, with the emergence of the concept of riboregulation, we have the opportunity to rethink this model and consider the possibility that proteins could also be regulated through interactions with RNA, especially in unexpected pathways such as cell division (5) - unexpected since one might not expect major activity of RNA and RBPs given the global transcriptional and translational repression during cell division (70–72). On the contrary, we identified multiple well-characterized major mitotic players such as AURKA, TPX2 and KIFC1 as RBPs, more specifically as unconventional RBPs. This raised the question of the functional meaning of their interactions with RNA. AURKA is one of the major regulators of the cell cycle, and an oncogene involved in tumorigenesis in several types of cancers that promotes centrosome amplification and tumor growth (51,55,73,74). It has evolved as an important anti-cancer target with several small molecule inhibitors being in clinical trials (75,76). Here, we investigated the RNA dependence of AURKA and its interactions. While the interaction of some interactors to AURKA was RNase-insensitive, other interactions were lost upon RNase treatment. Also, we revealed that 90% of AURKA interactors were classified as RNA-dependent proteins themselves. We demonstrated that the interaction of AURKA with TPX2, an essential mitotic interactor, was RNA dependent. Most importantly, we uncovered the interaction of both AURKA and TPX2 with KIFC1, which had not been studied before. These interactions were also RNA-dependent. Interestingly, the mitotic R-DeeP profile of AURKA, TPX2 and KIFC1 depicted similar peaks around fraction 21 of the control gradient, indicating that they co-migrated to the same fraction as one complex in the presence of RNA. Together with the proximity labeling assays, our findings strongly suggest that the three proteins are part of the same complex, which relies on RNA to be assembled. RNA is essential for such close proximity of the proteins, which is in turn important for their respective phosphorylation and activation (46,47), including the phosphorylation of KIFC1 at S^349^ and T^359^ by AURKA.

A fundamental step in understanding the molecular and cellular functions of RBPs is to identify their interacting RNAs. After demonstrating that AURKA, TPX2 and KIFC1 directly bound to RNA, we analyzed per iCLIP2 the KIFC1-interacting RNAs and found predominantly rRNA and protein-coding transcripts, lacking specific binding site sequences though. Large amounts of rRNA can be interpreted as contamination, a reason why rRNA is very often omitted from further analyses. However, recent publications have pointed to the enrichment of rRNA around the mitotic chromosomes and highlighted their role in mediating chromosome clustering (77–79). Furthermore, rRNAs are also associated with the microtubules and mitotic spindles (37,40). These data indicate an essential role of rRNA in spindle assembly and mitotic progression, although the molecular mechanisms at play remain unclear. Similarly, mRNAs are attached to the spindle, often mRNAs of mitotic factors, consistent with spindle-localized protein synthesis (35,36,80,81). In addition, a study also pointed to the role of RNAs in regulating protein localization to the mitotic spindle (82). Altogether, those previous findings confirm that rRNA and mRNA species are found at the mitotic microtubules and spindle and can effectively interact with mitotic factors, as indicated from our KIFC1 iCLIP2 data.

AURKA, TPX2 and KIFC1 belong to the class of unconventional RBPs, because they lack known RBDs. Currently, the RNA-binding characteristics of such proteins are not well understood, but a recent study reported the lack of sequence specificity for the great majority of the 492 investigated unconventional RBPs (83) - well in line with our findings for KIFC1. The same study also claimed that these unconventional RBPs were often highly abundant proteins and suggested that their identification in RNA interactome studies could occur via weak non-sequence-specific interactions with RNA. Thus, it might not be surprising that we did not identify any sequence specificity in the binding sites of the KIFC1-interacting RNAs. In addition, AURKA, KIFC1 and TPX2 all have high isoelectric points due to their positively charged amino acids and contain intrinsically disordered regions. Both aspects facilitate their binding to RNA. Thus, one can speculate that unspecific interaction with RNA possibly stabilizes their structure via a disorder-to-order transition, which in turn promotes interactions with other molecules, such as proteins or RNAs (84,85). However, further targeted investigations of the RNA-protein interactions, *e.g.* for selected unconventional mitotic RBPs, will certainly help understanding the driving mechanisms, that might not be simply limited to sequence characteristics.

For nuclear proteins such as TPX2 and KIFC1 (59,86), the proteins can already interact with nuclear-retained mRNAs throughout interphase. However, at the onset of cell division, they are suddenly exposed to the highly abundant cytoplasmic pool of both mRNA and rRNA transcripts (87,88) upon nuclear envelope breakdown. In the context of riboregulation, it is conceivable that multiple RNA transcripts collectively “crowd-control” AURKA, KIFC1 and TPX2 interactions. This hypothesis may be also expanded to other RBPs in mitosis and may provide the starting point for a more general concept of crowd-controlled riboregulation upon nuclear envelope breakdown in mitosis to be further investigated in the future. This idea also fits very well to the concept of self-organization, defined as the emergence of an ordered pattern in space and time, as the result of the collective interactions of individual constituents. Noteworthy, it is well-established, that such physical principle is the basis of the “self-organized” mitotic spindle (89) and other key cellular structures (90).

In sum, our study provides an atlas of RNA-dependent mitotic factors, suggesting an important role of RNA in orchestrating multiple aspects of cell division. Our detailed analysis of the RNA dependence of AURKA, TPX2 and KIFC1 reveals the importance of RNA in riboregulating mitotic protein-protein interactions, and suggests functional implications in many substructures of the dividing cells. This view is supported by past studies but still a matter of debate (40–42,49,82). To address these challenging questions within the native cellular environment, dedicated tools and new technologies will be needed. Thus, a completely new complexity level including mitotic protein-RNA interactions promises to advance our understanding of the molecular mechanisms regulating cell division.

## Supporting information

Supplementary Tables and R-DeeP3 User Guide

## Supplementary information

Supplementary information consists in the following files:

- Supplementary Figures S1 to S9
- Supplementary Tables S1 to S5
- User Guide for the R-DeeP 3.0 database

## Data availability

The R-DeeP 3.0 database is available online at https://R-DeeP3.dkfz.de.

The accession number for the R-DeeP screen proteomic dataset is PXD050038 at ProteomeXchange.

The accession number for the AURKA interactor analysis proteomic dataset is PXD056233 at ProteomeXchange.

The accession number for the KIFC1 iCLIP2 datasets (4 replicates) is E-MTAB-14472 at ArrayExpress.

## Acknowledgements

The authors sincerely acknowledge the IT core facility of the German Cancer Research Center (DKFZ) for their precious support and help in deploying the R-DeeP 3.0 database. This study would not have been realized without the support of the microscopy core facility of the DKFZ (Felix Bestvater, Manuela Brom and Damir Krunic) and the support of all the lab members especially Malte Hermes. We sincerely thank Anna Orekhova for her support with the iCLIP protocol, and Melina Klostermann and Mirko Brüggemann for their support and advice with the analysis of the iCLIP datasets. We also thank the high-throughput sequencing core facility of the DKFZ, the Protein Expression and Purification Core Facility at the EMBL, and our collaborators for the feedback on the database, experimental design, figures and manuscript.

## Author contributions

M.C.-H. conceived the study. M.C.-H. and S.D. supervised the R-DeeP screens, the statistical analysis, and the analyses of the RNA dependence of the proteins. A.N.K. supervised the mass spectrometry analyses. J.K. and K.Z. supported the implementation of the iCLIP protocol and downstream bioinformatic analysis. M.S. and D.H. performed mass spectrometry for AURKA immunoprecipitation and provided strong support for the downstream analysis. V.R., J.S., I.N., S.C., J.T. and F.H. performed the experiments. B.K. integrated the datasets into the R-DeeP 3.0 database. V.R. wrote the initial manuscript. M.C.-H. and S.D. edited the manuscript including the contributions of the co-authors. All the authors have seen and approved the final version of the manuscript.

## Detailed Methods Section

For catalog numbers and further details such as sequences and URL please refer to Supplementary Table S5.

### Gene ontology analysis

A gene ontology (GO) enrichment analysis was performed using the GO enrichment analysis tool from the Gene Ontology Resource (91) on RNA-dependent proteins as identified in the R-DeeP screen performed in HeLa cells (1751 proteins in total) (26).

### Cell culture

Hela WT cells and HeLa WT cells stably expressing GFP-α-tubulin (92) were grown in DMEM high glucose medium supplemented with 10% FBS and A549 WT cells were grown in RPMI 1640 medium supplemented with 10% FBS in a humidified incubator at 37°C with 5% CO2. Both cell lines were obtained from the lab of Prof. Diederichs, authenticated and frequently tested for the absence of mycoplasma contamination.

### Cell synchronization

On day 1, 2.5-3 million cells were seeded for synchronization in prometaphase and 2 million cells were seeded for synchronization at interphase in 15 cm dishes. On day 2, thymidine was added to the cells (2 mM final concentration) for 16 h to arrest the cells in S-phase. The cells were washed once with warm PBS and released with fresh media for 9 h. On day 3, thymidine was added again to the cells (2 mM final concentration) for 16 h to arrest the cells in S-phase (double thymidine block). To further synchronize the cells in prometaphase, the cells were washed once with PBS and fresh media was added to that contained 100 ng/ml nocodazole. Finally, after 12 h, the cells were harvested, flash frozen and stored at -80°C until lysate preparation.

To synchronize cells at metaphase, 5 million cells were seeded on a 15 cm plate on day 1. On day 2, thymidine was added to the cells (2 mM final concentration) for 24 h to arrest the cells in S-phase. On day 3, cells were washed once with warm PBS and further incubated with fresh medium containing 40 ng/ml nocodazole for 12 h. Cells were washed three times with warm PBS and released with fresh media for 40 min. Finally, the cells were harvested, flash frozen and stored at -80°C until lysate preparation.

### R-DeeP screen

*Sucrose density gradients*: the cells were cultured and synchronized in prometaphase and interphase as stated above. The gradients, cell lysate preparation, RNase treatment, ultracentrifugation and fractionation were performed as previously published (26,27). Briefly, the cell lysates remained either untreated (control gradients) or treated with an RNase cocktail (RNase-treated gradients), loaded on the sucrose gradients and subjected to ultracentrifugation. The gradients were fractionated to 25 different fractions, and further analyzed either via mass spectrometry (proteome-wide analysis on a Fusion Orbitrap Lumos mass spectrometer) or Western blot analysis (individual candidate analysis).

*Bioinformatic analysis*: mass spectrometry datasets were analyzed as explained in detailed in our published protocol (27).

### SDS-PAGE and Western blot analysis

Sodium dodecyl sulphate–polyacrylamide gel electrophoresis (SDS-PAGE) was performed to separate proteins based on their molecular weight. The samples in 1× SDS sample buffer (30% (v/v) glycerol, 12% (w/v) SDS, 3.6 M DTT, 0.012% (w/v) bromophenol blue, and 500 mM Tris-HCl (pH 6.8)) or 1x LDS buffer were boiled at 95°C for 5 min or 70°C for 10 min respectively, and briefly spun down. Next, the samples were loaded onto appropriate Bio-Rad pre-cast protein gels and run at 120 V in the electrophoresis chamber containing 1x SDS Running buffer (25 mM Tris base, 192 mM glycine, 0.1% (w/v) SDS).

Western blot analysis was performed on nitrocellulose membrane in a Trans blot turbo wet transfer system using 1x Trans-Blot Turbo Transfer Buffer containing 20% ethanol (mixed molecular weight program). The membrane was blocked with 5% milk in Tris-buffered saline (blocking solution: 24.7 mM Tris-HCl (pH 7.4), 137 mM NaCl, 2.7 mM KCl) containing 0.05% Tween-20 (TBST) for 1 h at room temperature (RT). Furthermore, the membrane was incubated at 4°C overnight with the respective antibodies. The following day, the membrane was washed three times with TBST for 5 min with at RT and incubated with the appropriate HRP-conjugated secondary antibody at 1:5000 dilution in blocking solution for 1 h at RT. The membrane was washed three times with TBST for 5 min at RT. Finally, the membrane was incubated with ECL reagent for 5 min and the blots were imaged using an INTAS ECL Chemocam imager. Quantitative analysis of Western blot images was performed using Fiji (Image J) software.

### AURKA Immunoprecipitation (IP) followed by LC-MS/MS-based protein analysis

*AURKA immunoprecipitation*: Hela cells synchronized in prometaphase were used for immunoprecipitation. The cell pellets were lysed in three pellet volume of lysis buffer (50 mM HEPES-KOH, pH-7.5, 150 mM KCl, 0.5% NP-40, 2 mM EDTA, 1 mM NaF, 0.5 mM DTT and 1x complete EDTA free protease inhibitor cocktail), incubated on ice for 30 min and centrifuged at 17,000 g for 20 min at 4°C. The supernatant was transferred to fresh tubes and centrifuged again for 17,000 g for 20 min at 4°C. The supernatant was stored in a fresh tube on ice until the beads were prepared for the pre-clearing step (see below).

30 µl of pierce ChIP-grade protein A/G magnetic beads were used per sample in the IP. The beads were aliquoted to a fresh 1.5 ml tube and washed three times with 1 ml lysis buffer. Next, the lysate was added on the beads and incubated for 1 h at 4°C on a rotator to remove unspecific binding to the beads. After 1 h, the beads were removed from the lysate on a magnetic stand and the pre-cleared lysate was transferred to fresh 1.5 ml tubes. A BCA assay was performed to measure the protein concentration.

The lysate was split into 2 different samples (containing 4 mg total lysate each) for overnight protein-antibody complex formation at 4°C on a rotator. Here, 0.8 µg AURKA antibody was used for AURKA IP and rabbit IgG was used as a negative control.

On the next day, beads were prepared by washing them three times in 1 ml lysis buffer. The washed beads were split in 2 tubes, each lysate-antibody mix was added to the beads and was incubated for 2 h at 4°C on a rotator. After incubation, the beads were removed from the lysate on a magnetic stand and the flow through (FT) was discarded. The beads-antibody-protein complexes were washed three times with 1 ml wash buffer I (50 mM HEPES-KOH, pH-7.5, 150 mM KCl, 0.5% NP-40, 0.5 mM DTT and 1x complete EDTA free protease inhibitor cocktail). Within the last wash step, each tube was split into two tubes for control and RNase treatment (total 4 tubes). Using a magnetic stand, the supernatant was discarded and the beads were resuspended with 100 µl wash buffer I.

For the control tubes, 10 µl wash buffer I was added to the sample. For the RNase-treated samples, 10 µl RNase cocktail was added (RNase cocktail: equal volume of RNase A, RNase I, RNase III, RNase H and RNase T1) and incubated for 1 h at 4°C on a rotator. After incubation, the beads were captured using a magnetic stand and FT were collected for LC-MS/MS-based protein analysis. The beads were washed three times with wash buffer II (50 mM HEPES-KOH, pH-7.5, 300 mM KCl, 0.5% NP-40, 0.5 mM DTT and 1x complete EDTA free protease inhibitor cocktail). Finally, the protein complexes were eluted using 30 µl 1x LDS containing 100 mM DTT and were analyzed using SDS-PAGE/Western blot and LC-MS/MS-based protein analysis at Proteomics Core Facility (Mass spectrometry-based protein analysis unit).

*Protein digestion of AURKA IP samples for LC-MS/MS analysis*: IP eluates were run for 0.5 cm into an SDS-PAGE and the entire piece was cut out and digested using trypsin according to Shevchenko *et al.* (93) adapted on a DigestPro MSi robotic system (INTAVIS Bioanalytical Instruments AG).

*LC-MS/MS analysis of AURKA IP*: The LC-MS/MS analysis was carried out on an Ultimate 3000 UPLC system (Thermo Fisher Scientific) directly connected to an Orbitrap Exploris 480 mass spectrometer for a total of 120 min. Peptides were online desalted on a trapping cartridge (Acclaim PepMap300 C18, 5 µm, 300 Å wide pore; Thermo Fisher Scientific) for 3 min using 30 µl/min flow of 0.05% TFA in water. The analytical multistep gradient (300 nl/min) was performed using a nanoEase MZ Peptide analytical column (300 Å, 1.7 µm, 75 µm x 200 mm, Waters) using solvent A (0.1% formic acid in water) and solvent B (0.1% formic acid in acetonitrile). For 102 min the concentration of B was linearly ramped from 4% to 30%, followed by a quick ramp to 78%, after two minutes the concentration of B was lowered to 2% and a 10 min equilibration step appended. Eluting peptides were analyzed in the mass spectrometer using data dependent acquisition (DDA) mode. A full scan at 120k resolution (380-1400 m/z, 300% AGC target, 45 ms maxIT) was followed by up to 2 seconds of MS/MS scans. Peptide features were isolated with a window of 1.4 m/z, fragmented using 26% NCE. Fragment spectra were recorded at 15k resolution (100% AGC target, 54 ms maxIT). Unassigned and singly charged eluting features were excluded from fragmentation and dynamic exclusion was set to 30 s. Each sample was followed by a wash run (40 min) to minimize carry-over between samples. Instrument performance throughout the course of the measurement was monitored by regular (approx. one per 48 h) injections of a standard sample and an in-house shiny application.

*Data analysis of LC-MS/MS data*: Data analysis was carried out by MaxQuant (94) using an organism specific database extracted from Uniprot.org (human reference database, containing 74,811 unique entries from 27^th^ February 2020). Settings were left at default with the following adaptions. Match between runs (MBR) was enabled to transfer peptide identifications across RAW files based on accurate retention time and m/z. Fractions were set in a way that MBR was only performed within replicates. Label free quantification (LFQ) was enabled with default settings. The iBAQ-value (95) generation was enabled. Peptides from AURKA interactors were analyzed using the log2(iBAQ) values from four replicates. Replicate four was excluded from the analysis due to unproper clustering pattern as compared to the other three replicates. Three conditions were analyzed (IgG IP, AURKA IP, AURKA IP RNase treatment in three replicates, 9 samples in total). Interactors were filtered and only further analyzed if they were detected in at least 70% of the samples. Missing values were imputed using random values based on a gaussian distribution centered around the median of the sample and outliers were adjusted based on the mean method. Ratios between the (AURKA IP)/(IgG) samples and (AURKA IP)/(AURKA IP RNase treatment), *i.e.* differences of the log2(iBAQ) values were calculated for each replicate and adjusted p-values were computed by applying a t-test, corrected for multiple testing (FDR method). AURKA interactors were selected based on an at least two-fold increased (AURKA IP)/(IgG) ratio (adjusted p-values < 0.05). RNase sensitive AURKA interactors were identified based on an at least two-fold increased (AURKA IP)/(AURKA IP RNase treatment) ratio (adjusted p-values < 0.05).

### AURKA Immunoprecipitation (IP)

HeLa and A549 cells synchronized in prometaphase or metaphase as explained above were used for immunoprecipitation followed by Western blot analysis. Each cell pellets were lysed in 2 ml lysis buffer (50 mM HEPES-KOH, pH-7.5, 150 mM KCl, 0.5% NP-40, 2 mM EDTA, 1 mM NaF, 0.5 mM DTT and 1x complete EDTA free protease inhibitor cocktail) as explained in the previous section. BCA assay was performed to measure the protein concentration in the lysate. The lysate was diluted to 2 mg/ml with lysis buffer and the total lysate was split into 4 tubes containing 2 mg lysate each.

2 µl of turbo DNase were added to each tube. 10 µl of lysis buffer was added to control samples while, 10 µl RNase I was added to RNase-treatment samples and were incubated at 37°C for 3 min at 1100 rpm in a thermomixer to digest the DNA and RNA in the lysates. The samples were cooled down by incubating on ice for 3 min. Further, the samples were centrifuged at 17,000 g for 20 min at 4°C and the supernatants were transferred to fresh tubes. The lysates were filtered through a proteus clarification mini spin column by centrifuging at 16,000 g for 2 min at 4°C. The filtered lysates were then transferred to fresh 2 ml tubes and kept on ice until the beads were prepared for the pre-clearing step (see previous section).

The lysates were incubated with the respective antibodies overnight at 4°C on a rotator to form protein-antibody complexes. For AURKA IP: 0.4 µg AURKA antibody per IP and rabbit IgG was used as a negative control. For TPX2 IP: 1.5 µg TPX2 antibody per IP and mouse IgG1 was used as a negative control.

On the next day, the beads were prepared by washing three times in 1 ml lysis buffer. The washed beads were added to the lysate-antibody mix and incubated for 2 h at 4°C on a rotator. After incubation, the beads were removed from the lysate on a magnetic stand and the FT was discarded.

For AURKA IP: the beads-antibody-protein complexes were washed three times with 1 ml wash buffer (HeLa: 50 mM HEPES-KOH, pH-7.5, 150 mM KCl, 0.5% NP-40, 0.5 mM DTT and 1x complete EDTA free protease inhibitor cocktail, A549: 50 mM HEPES-KOH, pH-7.5, 15 mM KCl, 0.5% NP-40, 0.5 mM DTT and 1x complete EDTA free protease inhibitor cocktail). Later, the beads were resuspended in 20 µl lysis buffer, 2 µl RNase I and incubated at 37°C for 3 min at 1100 rpm in a thermomixer. Lastly, the samples were eluted by adding 7.5 µl of 4x LDS (with 200 mM DTT) and boiling at 70°C for 10 min.

For TPX2 IP: the beads-antibody-protein complexes were washed three times with 1 ml wash buffer (50 mM HEPES-KOH, pH-7.5, 300 mM KCl, 0.5% NP-40, 0.5 mM DTT and 1x complete EDTA free protease inhibitor cocktail). Finally, the samples were eluted by adding 30 µl of 1x LDS (with 100 mM DTT) and boiling at 70°C for 10 min.

The samples were stored in fresh tubes at -20°C until SDS-PAGE/Western blot analysis.

### Proximity Ligation Assay (PLA)

All the buffers and solutions were provided in the Duolink® in-situ PLA kit.

120,000 cells/well were seeded on a coverslip in a 12-well plate and were allowed to grow overnight. On the following day, cell medium was aspirated, washed once with warm PBS and fixed with either methanol or 4% PFA (depending on the antibody) for 10 min or 15 min respectively and RT. After fixation, the cells were permeabilized in 0.25% Triton in PBS for 10 min at RT. After permeabilization, cells were washed twice with warm PBS and blocked with 40 μl Duolink® blocking solution for 1 h at 37°C in a heated humidity chamber. Meanwhile, primary antibodies were diluted in the Duolink® antibody diluent to appropriate concentrations. After blocking, the cells were incubated with primary antibody overnight at 4°C in a humidity chamber. All subsequent steps were performed following the instructions as provided with the Duolink® in-situ PLA kit. Mouse or rabbit secondary antibody Alexa Fluor 488 was added to the amplification mix in a 1:500 dilution.

For PLA +/- RNase: 200,000 cells/well were seeded on a coverslip in a 12-well plate. The cells were synchronized in metaphase based on the protocol described above. Following the synchronization, the media was removed, and the cells were first treated with 0.1% Triton in PBS with RNase and without RNase for control slides for 30 s at RT. Further, the cells were fixed with methanol at RT for 10 min, washed once with warm PBS and permeabilized with 0.25% Triton for 10 min at RT. After the permeabilization, the cells were washed twice with warm PBS and the rest of the PLA protocol from blocking step was followed as described above.

*Imaging*: images were acquired on a Zeiss LSM 980 Airyscan NIR, in confocal acquisition mode (best signal setting) or on a Zeiss LSM 710 ConfoCor 3, both equipped with diodes for the excitation of DAPI (405 nm), Alexa 488 (488 nm) and Alexa 594 (561 nm/555nm) fluorophores. Samples within one replicate were all acquired with the same settings. Z-stacks were acquired and maximal projection images (512 x 512 pixels, 8 bits) were analyze using Fiji (ImageJ) software. For intensity calculation, background pixel values up to a value of 10 were removed.

### Individual-nucleotide resolution UV Cross-linking and Immunoprecipitation (iCLIP2)

Frozen pellets from HeLa and A549 cells synchronized in prometaphase (AURKA and KIFC1) or metaphase (TPX2) were used for the iCLIP2 assay.

*Beads-antibody preparation*: the antibodies were conjugated to beads first. Dynabeads Protein A for AURKA/KIFC1 and Dynabeads protein G beads for TPX2 iCLIP2 (100 µl per IP) were washed three times with 1 ml lysis buffer. After the last wash the beads were re-suspended in 500 µl (AURKA/KIFC1 iCLIP2) or 400 µl (TPX2 iCLIP2) lysis buffer and split into two tubes, one for the IgG control (100 µl) and one for the protein of interest AURKA/KIFC1/TPX2 (400 µl/300 µl), and further incubated with the antibodies (IgG/AURKA/KIFC1: 2 µg per IP and IgG_1_/TPX2: 8 µg per IP) for 1 h at RT on a rotator (10 rpm). The bead–antibody complexes were captured on a magnetic rack and washed once with 1 ml high-salt wash buffer (50 mM Tris-HCl pH 7.4, 1.5 M NaCl, 1 mM EDTA pH 8.0, 1% Igepal CA-630, 0.1% SDS, 0.5% sodium deoxycholate) and twice with 1 ml lysis buffer. The beads were then resuspended in 100 µl lysis buffer for IgG or 400 µl/ 300 µl lysis buffer for AURKA, KIFC1 and TPX2 pulldown.

*Cell lysis*: UV cross-linked (254 nm, 200 mJ/cm^2^) and non-crosslinked cells were lysed in 2 ml lysis buffer per cell pellet (50 mM Tris-HCl pH 7.4, 100 mM NaCl, 1% Igepal (CA-630), 0.1% SDS, 0.5% Sodium deoxycholate, 1x protease inhibitor cocktail). BCA assay was performed to measure the protein concentration and the lysate was diluted to 2 mg/ml (AURKA/KIFC1) and 5 mg/ml (TPX2). The lysate was distributed into different 1.5 ml low-bind tubes containing 1 ml total lysate. The lysates were treated with 4 µl turbo DNase and different RNase I dilutions ranging from 1:5 to 1:1000 for AURKA, 1:5 to 1:500 for KIFC1 and 1:5 to 1:50 for TPX2. The RNase and DNase treatments were performed at 37°C at 1100 rpm on a thermomixer for 3 min. Next, the samples were incubated on ice for 3 min and centrifuged at 17,000 g for 20 min at 4°C. The supernatant was collected into a fresh 2 ml low-bind tube and filtered through proteus clarification mini spin column by centrifuging at 16,000 g for 2 min at 4°C. Further, the filtered lysates were transferred to a fresh 2 ml low-bind tubes and kept on ice.

*Pulldown/Immunoprecipitation*: 100 µl of the resuspended beads were added to the respective tubes containing cleared lysate and incubated for 2 h rotating at 4°C for immunoprecipitation. After 2 h, the complex was captured on a magnetic rack. The FT was removed and the beads were washed twice with 1 ml high-salt wash buffer with rotation at 10 rpm at 4°C for 1 min and then washed twice with 1 ml PNK wash buffer (KIFC1:20 mM Tris-HCl pH 7.4, 10 mM MgCl_2_, 0.2% Tween-20, AURKA/ TPX2: 20 mM Tris-HCl pH 7.4, 10 mM MgCl_2_, 0.2% Tween-20, PhosStop). During the last wash, the beads were transferred to fresh 1.5 ml low-bind tubes and stored at 4°C.

On the following day, for AURKA/TPX2 iCLIP2: the samples were placed on magnetic stand, PNK buffer containing PhosStop was removed and the beads were resuspended in 1ml PNK buffer (20 mM Tris-HCl pH 7.4, 10 mM MgCl_2_, 0.2% Tween-20) without PhosStop.

Further, 100 µl of beads were transferred to a fresh tube for Western blot and 900 µl were used for labelling the RNA. For Western blot, beads were captured, the supernatant was removed, and the protein complexes were eluted using 1× LDS buffer containing 50 mM DTT at 70°C for 10 min. The eluate was collected and stored at −20°C to check for immunoprecipitation efficiency using Western blot analysis. The SDS-PAGE and Western blot analysis was performed as described earlier.

*RNA labelling with radioactive ^32^P*: RNA labelling was performed using the remaining 900 µl sample. The radioactive labelling of RNA using ^32^P was performed using a master mix containing 11.85 µl nuclease free water, 0.75 µl T4 PNK enzyme, 1.5 µl 10× PNK buffer, and 0.9 µl ^32^P-γ-ATP per sample. Beads were captured on ice, the supernatant was removed and the beads were resuspended in 15 µl PNK mix. The samples were incubated on a thermomixer at 37°C for 5 min at 1100 rpm for labelling the RNA. Later, the samples were washed twice with 1 ml PNK wash buffer to get rid of excess radioactivity and eluted in 25 µl 1× LDS buffer containing 50 mM DTT on a thermomixer at 70°C for 10 min at 1100 rpm.

To visualize the RNA-labelling, SDS-PAGE and Western blot analysis were performed. The samples were loaded in a 7.5% Mini-PROTEAN® TGX™ precast protein gel and run 120 V in a vertical electrophoresis chamber filled with 1× SDS running buffer (25 mM Tris base, 192 mM glycine, 0.1% (w/v) SDS). Western blot was performed using 0.45 µm nitrocellulose membrane with a wet transfer system with transfer buffer (25 mM Tris base, 192 mM glycine) containing 20% methanol for 1.5 h at 120 V in an ice bath. Finally, the membrane was washed once in nuclease-free water, covered with plastic wrap and exposed to a phosphor imager screen and imaged using a Typhoon laser scanner phosphor imager at 200 µm, high speed and intensity 3.

### iCLIP2 library preparation and sequence analysis

The iCLIP2 library preparation was performed based on the publication “Improved library preparation with the new iCLIP2 protocol” (63). For adapter, barcodes or primer sequences refer to Supplementary Table S5.

For KIFC1 iCLIP2 library preparation, UV-crosslinked prometaphase cells synchronized and harvested on four different dates were used and IgG was used as a negative control. Here, RNase treatment was performed on the lysate at 1:10 dilution. All the steps from lysate preparation, beads preparation, and pulldown were performed as described in the above section (Individual Nucleotide Resolution and UV Cross-linked Immunoprecipitation (iCLIP2).

*Dephosphorylation*: 3’ dephosphorylation of RNA was performed using the master mix containing 1x PNK buffer pH-6.5 (350 mM Tris-HCl, pH 6.5, 50 mM MgCl_2_, 5 mM DTT), 0.5 µl SUPERase-In, 0.5 µl of T4 PNK enzyme) in a total volume of 15 µl per sample. The PNK buffer was removed from the samples, and the beads were resuspended in 20 µl of the dephosphorylation master mix and incubated for 20 min at 37°C at 1100 rpm on a thermomixer. After the incubation, the beads were washed once with 1 ml PNK wash buffer (20 mM Tris-HCl pH 7.4, 10 mM MgCl_2_, 0.2% Tween-20), twice with 1 ml high-salt wash buffer (50 mM Tris-HCl pH 7.4, 1.5 M NaCl, 1 mM EDTA pH 8.0, 1% Igepal CA-630, 0.1% SDS, 0.5% Sodium deoxycholate) for 2 min at 4°C on a rotator and twice again with 1 ml PNK wash buffer (20 mM Tris-HCl pH 7.4, 10 mM MgCl_2_, 0.2% Tween-20).

*3’ adapter ligation*: adapter ligation was performed using the following ligation mix containing (4x ligation buffer (200 mM Tris-HCl, pH 7.8, 40 mM MgCl_2_, 4 mM DTT, PEG 400, 3 µl of L3-App-Fluo adapter (10 µM) (/rApp/AGATCGGAAGAGCGGTTCAG/ddC/), 0.5 µl SUPERase-In, 1 µl of T4 ligase in a total volume of 6.5 µl per sample). The PNK buffer was removed from the beads and the beads were resuspended in 20 µl ligation mix and incubated overnight at 16°C at 1100 rpm on a thermomixer. The following day, the beads were washed once with 0.5 ml PNK wash buffer (20 mM Tris-HCl pH 7.4, 10 mM MgCl_2_, 0.2% Tween-20), twice with 1 ml high-salt wash buffer (50 mM Tris-HCl pH 7.4, 1.5 M NaCl, 1 mM EDTA pH 8.0, 1% Igepal CA-630, 0.1% SDS, 0.5% sodium deoxycholate) for 2 min at 4°C on a rotator and once again with 0.5 ml PNK wash buffer. During the last wash, the samples were transferred to a fresh 1.5 ml low-bind tubes and the beads were re-suspended in 1ml PNK wash buffer. 100 µl of the sample were aliquoted for Western blot analysis. The remaining 900 µl of the samples were used for RNA extraction. The samples were placed on a magnetic stand, buffer was removed and the samples were eluted using 35 µl 1x LDS buffer, boiled at 70°C for 10 min. The samples were loaded 7.5% Mini-PROTEAN® TGX™ precast protein gel and run at 120 V in a vertical electrophoresis chamber filled with 1× SDS running buffer (25 mM Tris base, 192 mM glycine, 0.1% (w/v) SDS). Western blot was performed using 0.45 µm nitrocellulose membrane using a wet transfer system with transfer buffer (25 mM Tris base, 192 mM glycine) containing 20% methanol for 1.5 h at 120 V on an ice bath.

*Proteinase K digestion*: Following the Western blot, the membrane was cut at 90 kDa-150 kDa to 4-5 small pieces and transferred into a 2 ml low-bind tube. Further the master mix (2x proteinase K buffer, 1mg proteinase K) for proteinase K digestion was prepared. 400 µl of the mix was added to each tube with cut membrane, vortexed for 20 seconds and incubated for 1 h 30 min at 55°C at 1000 rpm on a thermomixer.

*RNA extraction*: For RNA extraction, 2 volumes of acidic phenol-chloroform-IAA (pH 6.5-6.9) was directly added to the proteinase K digested samples, mixed by inverting for 15 s and incubated at RT for 5 min. Meanwhile, phaselock gel heavy tubes were prepared by spinning them at 12,000 g for 30 s. The supernatant except the membrane pieces were transferred to the prepared phaselock gel heavy tubes, incubated at RT for 5 min at 1200 rpm on a thermomixer and centrifuged at 17,000 g for 15 min at RT. The aqueous layer was transferred to new 2 ml low-bind tubes. Further, 2 volumes of RNA binding buffer from the Nucleospin plasmid isolation kit and equal volume of 100% ethanol were added and mixed well. The mixed samples were then transferred to Zymo-spin column, centrifuged at 15,000 g for 30 seconds at RT and the flow through was discarded. This step was repeated until all the samples were loaded and spun through the column. 400 µl RNA prep buffer was added to the column, centrifuged at 15,000 g for 30 seconds at RT and the flow through was discarded. Later, 700 µl of RNA wash buffer was added to the column, centrifuged at 15,000 g for 30 seconds at RT and the flow through was discarded. This step was repeated again with 400 ml RNA wash buffer with additional centrifugation for 2 min to get rid of excess RNA wash buffer. Finally, the RNA was eluted on fresh low-bind tube with nuclease free water by centrifuging at 15,000 g for 1 min at RT. The RNA was stored at -80°C until reverse transcription.

*Reverse transcription*: dNTPs and RT oligo mix (2 µl of RT oligo (1 µM), 1 µl dNTPs (10 mM each), and nuclease free water in a total volume of 5 µl per sample) were prepared and added to each RNA extracted from the membrane. The samples were mixes, briefly centrifuged and incubated in a thermomixer for 5 min at 65°C and then on ice for at least 1 min. 5x superscript IV buffer (SSIV) was vortexed, briefly centrifuge and the reverse transcription (RT) reaction mix was prepared (4 µl of 5x SSIV buffer, 1 µl 0.1 M DTT, 1 µl RNase OUT, 1 µl superscript IV in a total volume of 7 µl per sample). 7 µl of the RT reaction mix was added to each tube containing the RNA and incubated in a thermomixer at 25°C for 5 min, 42°C for 20 min, 50°C for 10 min, 80°C for 5 min and hold at 4°C. Later, 1 µl RNase H was added to each tube and incubated at 37°C for 20 min to cleave the RNA in the DNA-RNA hybrid.

*Cleanup I*: MyONE beads were used for the cleanup of the cDNA. The MyONE beads were mixed by vortexing and 10 µl beads were used per sample. The beads were washed once with 500 µl RLT buffer, resuspended in 125 µl RLT buffer and added to each sample and finally transferred to new 1.5 ml low-bind tube and mixed well by pipetting. 150 µl of 100% ice cold ethanol was added to the cDNA-beads complex, mixed well and incubated at RT for 5 min. The samples were further incubated for 5 min at RT after mixing them once again by pipetting up and down. Next, the samples were placed on a magnetic stand, the supernatant was discarded, and the beads were re-suspended in 900 µl of freshly prepared 80% ice cold ethanol and mixed by pipetting. The mix was transferred to a new low-bind tube, the supernatant was discarded and the above step was repeated twice. The tubes were spun briefly to remove as much ethanol as possible, air dried at RT. Finally, the beads were re-suspended in 5 µl nuclease free water.

*Adapter ligation*: 2 µl adapters L02clip2.0 (IgG), L02clip2.0 (KIFC1 replicate 1), L05clip2.0 (KIFC1 replicate 2), L10clip2.0 (KIFC1 replicate 3), L19clip2.0 (KIFC1 replicate 4), and L02clip2.0 (nuclease free water) from 10 µm stock were used from the publication Buchbender et al. (63). 1 µl of 100% DMSO was added to each tube, mixed well and heated on a thermomixer for 2 min at 75°C and the samples were immediately kept on ice for less than 1 min. The ligase mix (2 µl 10x NEB RNA ligase buffer with 10 mM DTT, 0.2 µl 100 mM ATP, 9 µl 50% PEG 8000, 0.5 µl high conc. RNA ligase in a total volume of 12 µl per sample) were prepared and added to the tubes containing the beads and the adapters samples and mixed well. Additionally, 1 µl of high conc. RNA ligase was added to each sample, mixed well and the samples were incubated overnight at RT (20°C) at 1100 rpm on a thermomixer.

*Cleanup II*: MyONE beads were used for the second cleanup procedure and steps were followed as mentioned in the cleanup I section (see above). At the end, the beads were resuspended in 23 µl nuclease free water, incubated at RT for 5 min, and the supernatant without the beads were transferred to new PCR tubes.

*cDNA pre-amplification*: Phusion master mix (2.5 µl P3Solexa_s and P5Solexa_s mix, 10 µM each, 25 µl 2x Phusion HF PCR master mix to a total volume of 27.5 µl per sample) was added to 22.5 µl cDNA and PCR amplification (98°C for 30 s, 98°C for 10 s, 65°C for 30 s, 72°C for 15 s, 72°C for 3 min) was performed for 6 cycles. The amplified cDNA was then size selected using the ProNex beads to reduce the primer-dimers formed during the PCR reaction (see below).

*ProNex size selection I*: To discard fragments less than 55 nt and to retain fragments longer than 75 nt, size selection using ProNex beads was performed. Ultra-low range (ULR) ladder was used for reference (1 µl ULR ladder, 49 µl water) and for size selection (1 µl ULR ladder, 25 µl 2x Phusion HF PCR master mix in a total volume of 50 µl). First, the beads were equilibrated to RT for 30 min on a rotator. 145 µl ProNex beads were added per sample (beads to sample ratio: 1:2.9), mixed well by pipetting and incubated at RT for 10 min. The supernatant was discarded, 300 µl ProNex buffer was added to the beads, incubated for 30-60 s and the supernatant was discarded. This step was repeated once more, the beads were air-dried and the beads were resuspended in 23 µl nuclease free water and the ULR ladder for size selection was resuspended in 50 µl nuclease free water and incubated at RT for 5 min. The samples were placed on a magnetic stand and the eluted cDNA was carefully transferred to a fresh PCR tube. The size was checked using ULR reference ladder and ULR ladder for size selection using sensitivity D1000 tape station kit.

*PCR cycle optimization*: 1 µl size selected cDNA was used for PCR cycle optimization. Phusion master mix (0.5 µl P3Solexa_s and P5Solexa_s mix, 10 µM each, 5 µl 2x Phusion HF PCR master mix to a total volume of 9 µl per sample) were added to 1 µl cDNA and PCR amplification (98°C for 30 s, 98°C for 10 s, 65°C for 30 s, 72°C for 30 s, 72°C for 3 min) was performed for 7 and 10 cycles. Depending on the amount of cDNA obtained and to limit the amplification within 10 cycles to minimize the PCR duplicates, for this library we decided to continue with PCR cycle 8 and 9 for preparative PCR.

*Preparative PCR*: Preparative PCR was performed with 10 µl cDNA for 8 and 9 cycles. Phusion master mix (2 µl P3Solexa_s and P5Solexa_s mix, 10 µM each, 20 µl 2x Phusion HF PCR master mix to a total volume of 30 µl per sample) was added to 10 µl cDNA and amplification was performed. 2 µl of the amplified library was used for run with a High Sensitivity D1000 tape station kit. The library from 8 and 9 cycles were combined and all four samples (KIFC1 replicates 1-4) were multiplexed. Further, a second size selection using ProNex beads was preformed to remove residual primer-dimers from the PCR amplification.

*ProNex size selection II*: ProNex selection was performed as described in the above section ProNex size selection I with samples to beads ratio of 1:2.2. Finally, the library samples were eluted in 63 µl nuclease free water and the concentration was measured using a Qubit device. The samples were sequenced using Illumina Inc., NextSeq 550 high output v2.5, 150 cycles, 320 million reads platform at the DKFZ high-throughput core facility.

*Sequence analysis*: The 150 nt long reads were mapped to hg38 human genome (GENCODE v39) using STAR 2.5.3a (96) and further analysis was performed using uniquely mapped reads, after evaluation of the amount of rRNA sequences as previously described (65,97). Binding sites were defined with BindingSiteFinder v2.0.0 as described in Busch *et al.* (64). The binding site width was set to 7 nt. 36934 binding sites were in common for at least 3 out of 4 replicates. Each binding site was assigned the maximum of the PureCLIP score from the replicates and the reads were overlapped with gene annotations from GENCODE on human genome (GENCODE v43). The binding sites with the 5% lowest and highest scores were removed and after assigning the binding sites to distinct genes and gene regions, the target spectrum of KIFC1 was generated (33240 binding sites remaining). The pentamer frequencies were evaluated in the top 20% binding site as compared to the bottom 20% binding sites.

### Cloning

For primer sequences and catalog numbers, please refer to Supplementary Table S5.

KIFC1 cDNA ORF clone in cloning vector was purchased from Sino biologicals. KIFC1 WT pENTRY clone was cloned into pDONOR221 plasmid by PCR (25 µl 2x Phusion high-fidelity PCR master mix, 1 µl DNA (10 ng in total), 2.5 µl forward and reverse primer (10 µM), 1.5 µl DMSO in a total volume 50 µl per reaction) amplification at 98°C for 1 min, 98°C for 10 s, 55°C for 30 s, 72°C for 1 min, 72°C for 10 min and hold at 4°C) for 40 cycles. The amplicon was run on 0.8% agarose gel, the band was cut out, purified using the GeneJET gel extraction kit according to the manufacturer’s instructions and finally eluted in 50 µl nuclease free water. Further, BP reaction (1 µl PCR product (∼50ng/µl), 1 µl pDONOR221 (150 ng/µl), 6 µl TE buffer pH 8, 2 µl 5x BP clonase enzyme mix in a total volume of 10 µl) was carried out at 25°C overnight in a PCR machine. Finally, 1 µl proteinase K was added, incubated at 37°C for 10 min in a PCR machine to terminate the reaction.

*Transformation*: One shot top 10 chemically competent *E. coli* cells were transformed with 1.5 µl BP reaction mix by incubating on ice for 30 min, heat shock at 42°C for 2 min, incubated on ice again for 2 min. Further, 250 µl LB medium was added to the cells and incubated for 1 h at 37°C with mild shaking. Later, 100 µl cells were plated out on LB agarose plates containing 10 µg/ml kanamycin and incubated overnight at 37°C for the bacteria to grow.

The following day, 2 colonies were picked per plate, plasmid isolation was performed using the Nucleospin plasmid isolation kit based on manufacturers protocol and were sequenced prior to use.

The KIFC1 WT sequences were shuttled into pFRT-Flag-HA-ΔCmR-ΔccdB vector using gateway cloning system for further use.

Serine and threonine residues were mutated to alanine-an amino acid residue which cannot be phosphorylated (KIFC1-S6A, KIFC1-S26A, KIFC1-S31A, KIFC1-S96A, KIFC1-T187A, KIFC1-S221A, KIFC1-S349A, KIFC1-T359A, KIFC1-S349A/T359A). The mutation was performed using the Q5 Site-Directed Mutagenesis Kit from NEB. The primers were ordered to perform single or double base substitutions using PCR amplification(1µg DNA template, 12.5 µl Q5 Hot Start High-Fidelity 2x Master Mix, 1.25 µl 10 µM forward and reverse primer each, in a total volume of 25 µl per sample) at (Initial denaturation 98°C for 30 seconds, denaturation at 98°C for 10 seconds, annealing at 61-72°C for 30 seconds depending on the individual primers, extension at 72°C for 3 min, final extension at 72°C for 2 min and hold at 4°C for 25 cycles). The mutations were performed on the KIFC1 WT sequence on pFRT-Flag-HA-ΔCmR-ΔccdB plasmid backbone. 10% of the PCR amplified product was run on 0.8% agarose gel to check of the amplicon size and integrity.

Transformation: Kinase, ligase and DpnI (KLD) treatment (PCR product 1 µl, 5 µl 2x KLD reaction buffer, 1 µl 10x KLD enzyme were added to a total volume of 10 µl per reaction) was performed prior to transformation for 5 min at RT. Following the treatment, 5 µl of the KLD reaction mixture was directly added to 50 µl NEB^®^ 5-alpha Competent *E. coli* (High Efficiency) cells, incubated for 30 min on ice, heat shock at 42°C for 30 seconds, followed by incubation on ice for 5 min. Further, 950 µl SOC media was added to the cells and incubated at 37°C for 1 h with gentle shaking. Later, 50-100 µl of the cells were plated on LB agar plate with appropriate selection antibiotics and incubated overnight at 37°C for the bacteria to grow.

The following day, 2 colonies were picked per plate/mutant, plasmid isolation was performed using the Nucleospin plasmid isolation kit (Macherey Nagel) based on manufacturers protocol and sequences were verified to use.

### *In vitro* kinase assay

*Transfection*: 5 million HeLa cells were plated out on a 15 cm dish on day 1. On day 2, they were transfected with plasmids (pFRT-Flag-HA-ΔCmR-ΔccdB as empty vector control, FlagHA-KIFC1_WT, FlagHA-KIFC1_S6A, FlagHA-KIFC1_S26A, FlagHA-KIFC1_S31A, FlagHA-KIFC1_S96A, FlagHA-KIFC1_T187A, FlagHA-KIFC1_S221A, FlagHA-KIFC1_S349A, FlagHA-KIFC1_T359A) using lipofectamine 2000 at a DNA:lipofectamine ratio of 1:2.5, based on the manufacturer’s instructions. After 6 h, the plates were washed once with warm PBS and fresh media was added to the cells. 24 h post transfection, the cells were harvested, the pellets were flash frozen and stored at -80°C until further use.

*Cell lysis*: Each cell pellet was lysed in 1 ml lysis buffer (50 mM Tris-HCl pH 7.4, 150 mM NaCl, 1 mM EDTA, 1 mM NaF, 1% triton X-100, 1x complete EDTA free protease inhibitor cocktail), incubated on ice for 30 min and centrifuged at 17,000 g for 20 min at 4°C. The supernatant was collected into a fresh 2 ml low-bind tube and filtered through proteus clarification mini spin column by centrifuging at 16,000 g for 2 min at 4°C. BCA assay was performed to determine the protein concentration and the lysates were diluted to 1.5 mg/ml using lysis buffer.

*Beads preparation and immunoprecipitation*: Anti-DYKDDDDK magnetic agarose beads were mixed by gentle vortexing and 37.5 µl beads were used per sample containing 1.5 mg lysate. The beads were first washed three times with 1 ml lysis buffer. After the last wash, the beads were transferred to a fresh 1.5 ml tube, collected on a magnetic stand, the supernatant was discarded and lysate was added to the beads. The lysate-beads mix were incubated for 2 h at 4°C on a rotator. Further, the samples were placed on a magnetic stand, the supernatant was removed and the beads were washed 4 times with 1 ml wash buffer (50 mM Tris-HCl pH 7.4, 350 mM NaCl, 1% triton X-100, 1x complete EDTA free protease inhibitor cocktail) for 5 min at 4°C on a rotating wheel. During the last wash, the beads were transferred to a fresh 1.5 ml tube, the supernatant was discarded and the beads were resuspended in 1 ml PNK buffer (20 mM Tris-HCl pH 7.4, 10 mM MgCl_2_, 0.2% tween-20, 1x complete EDTA free protease inhibitor cocktail) and stored at 4°C overnight. The following day, 100 µl of the samples were used for Western blot analysis to check for IP efficiency.

*In vitro kinase assay for single mutants*: 1x kinase assay buffer was prepared from 10 x kinase assay buffer (500 mM Tris-HCl pH 7.5, 100 mM NaCl, 100 mM MgCl_2,_ 10 mM DTT, 1x complete EDTA free protease inhibitor cocktail). The insect cell purified AURKA protein was diluted to 0.5 mg/ml in 1x kinase assay buffer and the kinase assay master mix was prepared (negative control without purified AURKA protein: 2 µl 1x kinase assay buffer, 2 µl 10x kinase assay buffer, 2 µl 1 mM cold ATP and 0.5 µl ^32^P-γ-ATP; kinase assay with purified AURKA protein: 2 µl purified AURKA protein 0.5 mg/ml, 2 µl 10x kinase assay buffer, 2 µl 1mM cold ATP and 0.5 µl ^32^P-γ-ATP in a total volume of 6.5 µl per sample). Further, supernatant was removed from the beads, 6.5 µl of the master mix was added to the samples and incubated for 30 min at 30°C in a thermomixer at 800 rpm. After the incubation, the samples were eluted by adding 5 µl 4x LDS supplemented with 200 mM DTT, boiled at 70°C for 10 min.

To visualize the phosphorylation signal, SDS-PAGE analysis was performed. The samples were loaded in a 7.5% Mini-PROTEAN® TGX™ precast protein gel and run at 120 V in a vertical electrophoresis chamber filled with 1× SDS running buffer (25 mM Tris base, 192 mM glycine, 0.1% (w/v) SDS). Later, the gel was fixed with 15 ml fixation solution (50% methanol, 10% acetic acid) for 1 h at RT with slow rocking, washed three times with nuclease free water for 5 min and dried for 1-1.5 h at 80°C. The dried gel was then exposed to phosphor imager screen and was scanned after appropriate amount of time using Typhoon laser scanner phosphor imager at 200 µm, high speed and intensity 3.

*In-vitro kinase assay for double mutant*: 5 million HeLa cells were seeded on a 15 cm dish. On the next day, the cells were transfected with plasmids (pFRT-Flag-HA-ΔCmR-ΔccdB (empty vector control), FlagHA-KIFC1_WT, FlagHA-KIFC1_S349A, FlagHA-KIFC1_T359A, FlagHA-KIFC1_S349A/T359A) using lipofectamine 2000 at DNA:lipofectamine ratio of 1:2.5, based on the manufacturer’s instructions. After 6 h, the plates were washed once with warm PBS and fresh media was added to the cells. 24 h post transfection, the cells were harvested, the pellets were flash frozen and stored at -80°C until further use.

The kinase assay was performed just as described above in this section. During the kinase reaction, insect cell purified AURKA kinase dead mutant D274A was used as a negative control and AURKA WT protein was used for the kinase activity to demonstrate the phosphorylation of KIFC1 WT vs the phosphorylation mutants. Further, after fixing the gel, the gel was first stained with Coomassie stain (to visualize protein bands) overnight at RT on a shaker, washed with nuclease free water for 1 h at RT, imaged and then dried and exposed to phosphor imager screen.

## Supplementary Figures

**Supplementary Figure S1.**
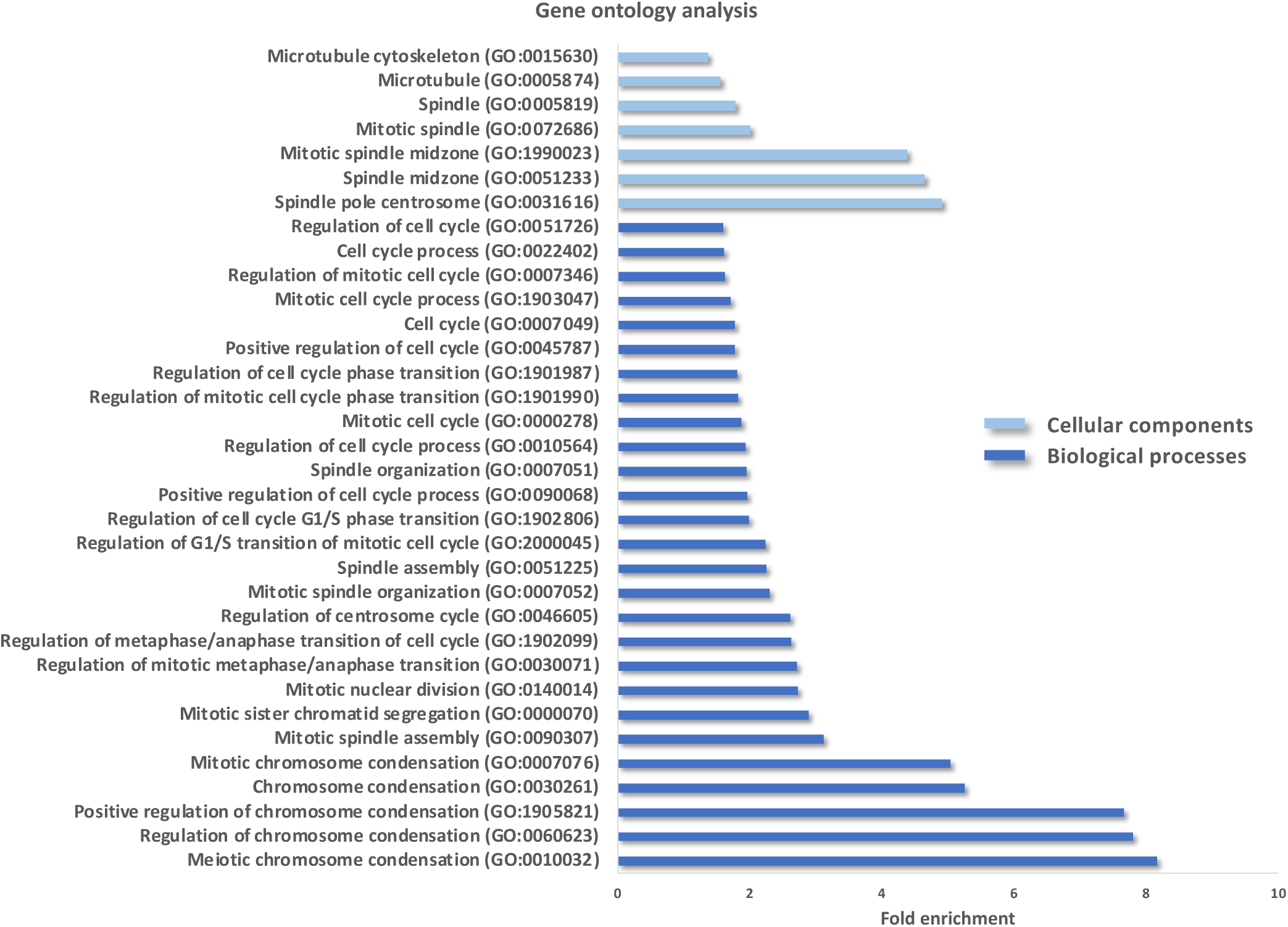
RNA-dependent proteins are enriched in mitotic factors. A gene ontology (GO) analysis was performed based on the list of RNA-dependent protein from the R-DeeP screen (27). An at least 2-fold significant enrichment was observed for various mitosis-related terms such as cell cycle regulation, spindle and microtubule (adjusted *P*-value < 0.05 according to a Fisher’s exact test and FDR correction for multiple testing). Bars in light blue indicate GO analysis for cellular components and bars in dark blue represent GO analysis for biological processes.

**Supplementary Figure S2.**
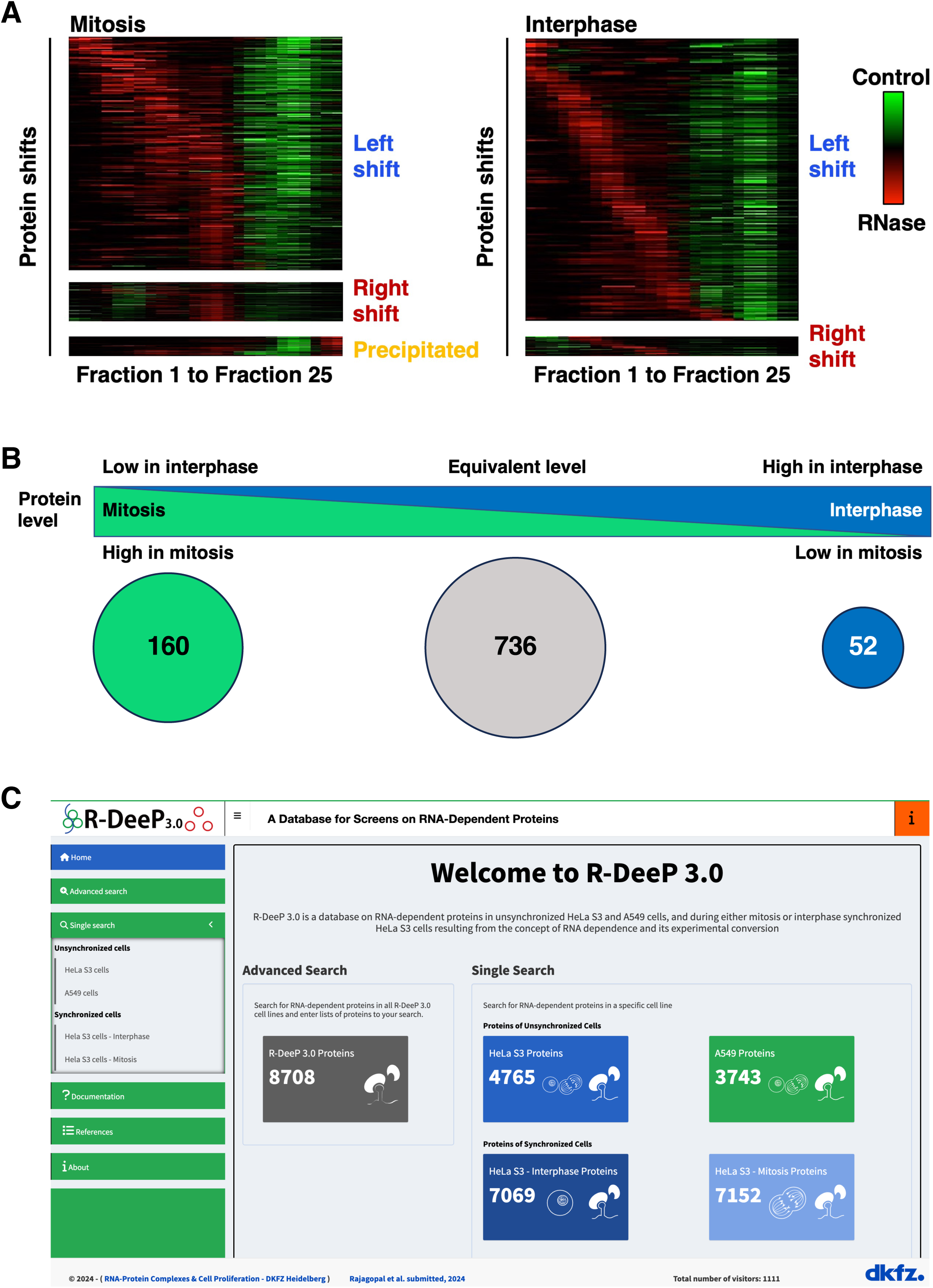
R-DeeP screens in mitosis and interphase. **A**: Heatmaps of the sub-categories for shifting proteins (left shift, right shift, or precipitated), representing the enrichment in the control (green) or in the RNase (red) fractions, for the mitotic and interphasic screen, respectively. **B**: Schematic representation of the differential protein levels of the common left shifting proteins in mitotic and interphasic R-DeeP screens that are performed in HeLa cells. While 736 proteins depict equal expression levels in both cell cycle phases, 160 proteins are higher expressed in mitosis and 52 proteins are preferentially expressed in interphase and thus show differential expression in a cell cycle dependent manner. **C**: The R-DeeP 3 database (https://R-DeeP3.dkfz.de) integrates comprehensive information on proteins that are detected in the R-DeeP screens in synchronized HeLa cells at mitosis and interphase, in addition to the knowledge from the two previous screens in unsynchronized HeLa and A549 cells. It provides detailed information on each protein in terms of shift, maxima and offers multiple search and download options. The database is linked to external resources for complementary knowledge on protein structure, sequence and interaction partners.

**Supplementary Figure S3.**
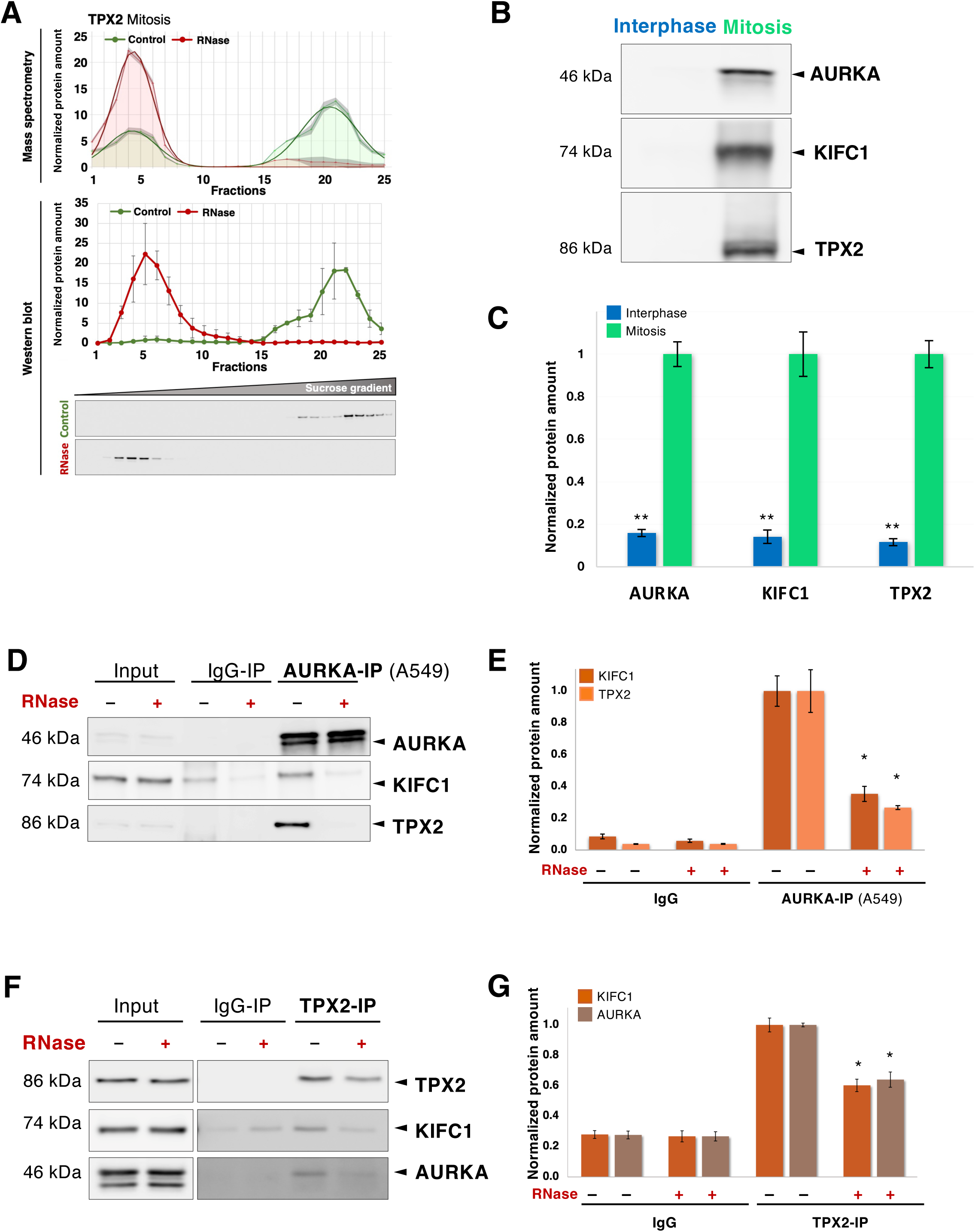
AURKA protein-protein interactors that are RNA-dependent in mitosis. **A**: R-DeeP profile of TPX2 in mitosis and WB validation (as for AURKA, see Figure 3, N=3). **B**: WB analysis showing the differential expression of AURKA, KIFC1 and TPX2 in interphasic and mitotic HeLa cell lysates (20 µg total lysate per lane). **C**: Bar graph representing the total protein amount in interphase and mitotic Hela cell lysates with SEM (N=3). The *P*-value was calculated using a two-tailed, paired t-test (** *P*-value <0.01). **D**: WB analysis showing the immunoprecipitation of AURKA in mitotic A549 cells. AURKA pulldown was performed in the mitotic lysate treated with or without RNase I. Mouse IgG was used as a negative control. KIFC1 (74 kDa) and TPX2 (86 kDa) were pulled down with AURKA (46 kDa) in control samples whereas their interaction was significantly reduced upon RNase treatment (reduction of the band intensity for KIFC1 and TPX2 in the last lane). **E**: Graph representing the amount of KIFC1 and TPX2 present in IgG and AURKA pulldown samples treated with or without RNase I. The intensities of the WB bands were quantified using ImageJ and represented in the bar graph with SEM (N=3). *P*-values were evaluated using a two-tailed, paired t-test (* *P*-value <0.05). **F** and **G**: same as in D and E, respectively, for TPX2 pulldown in mitotic HeLa cells synchronized in prometaphase. *P*-value was calculated using two-tailed, paired t-test (* *P*-value <0.05, N=3).

**Supplementary Figure S4.**
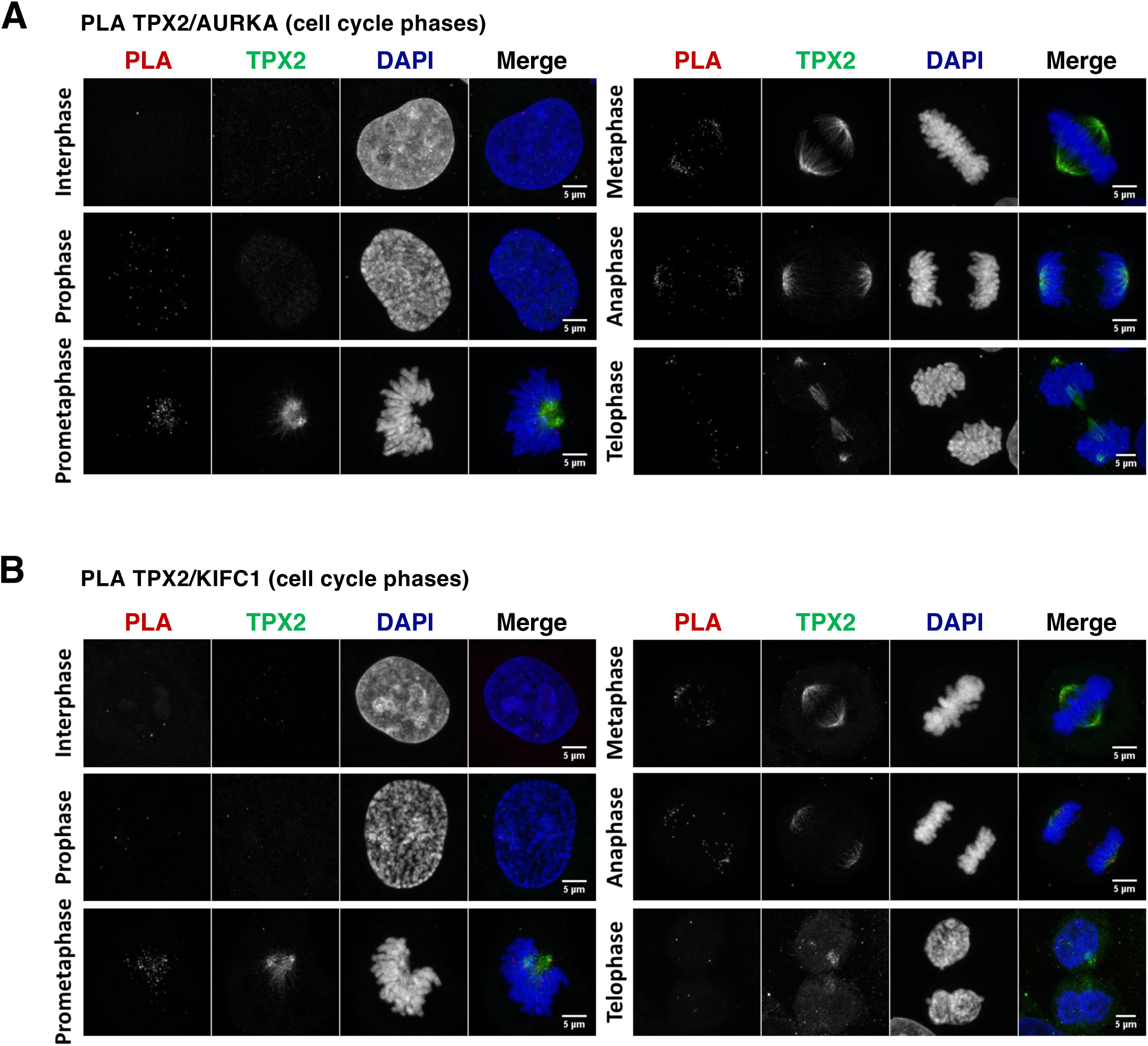
Protein-protein interactions in mitosis throughout the cell cycle. **A**: Representative PLA images indicating the close proximity of TPX2 and AURKA in HeLa cells across the cell cycle (interphase to telophase as indicated). The interaction is represented by dots (PLA channel, red dots in the merge channel), TPX2 was stained per immunofluorescence (green) and DNA was stained using DAPI (blue). Scale bar, 5 µm. **B**: same as in B for TPX2 and KIFC1.

**Supplementary Figure S5.**
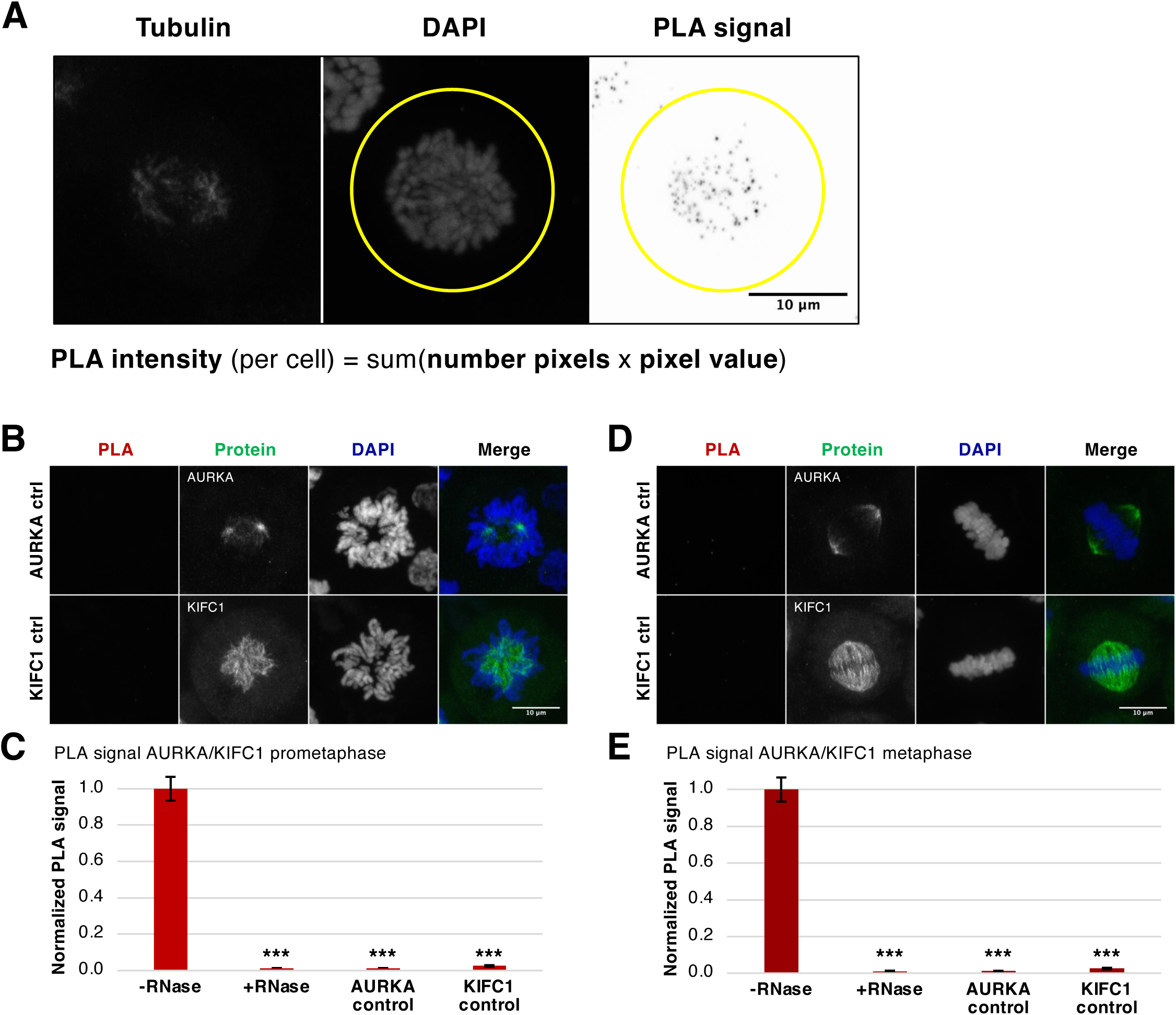
RNA-dependent protein interactions in mitosis demonstrated by Proximity Ligation Assay (PLA) **A**: Principle of the analysis of the PLA signal intensity. For each image series, a common circular region of interest (ROI) was placed around the chromosomes. After background signal depletion, the signal intensity (pixel values) of the PLA image was collected and the overall signal intensity inside the ROI was calculated as the sum of each pixel value times the number of pixel with this value. Scale bar, 10 µm. **B**: Representative PLA images of the PLA assay control samples in HeLa cells synchronized in prometaphase. The aim of these images is to estimate the amount of background PLA signal generated by the antibody alone and not via the proximity of the two proteins. Here, the cells were probed with only one antibody against AURKA or KIFC1 (in green) and further processed for the PLA assay. It appears, that each antibody alone did not produced any strong PLA. Scale bars, 10 µm. **C**: Bar graph of the quantification of the PLA signal (as explained in **A**) for the proximity of AURKA and KIFC1 in HeLa prometaphase cells in absence (-RNase) or presence (+RNase) of RNase treatment (see Figure 4G), as well as in the individual antibody control PLA assays. The signal intensities in each sample was normalized to the signal intensity of the -RNase sample. The error bars indicate the SEM (N=3). The *P*-value were calculated using a two-tailed, unpaired t-test (*** *P*-value <0.001). Related to Figure 4G **D** and **E**: same as in **B** and **C** respectively, for HeLa cells synchronized in metaphase (N=3, related to Figure 4H).

**Supplementary Figure S6.**
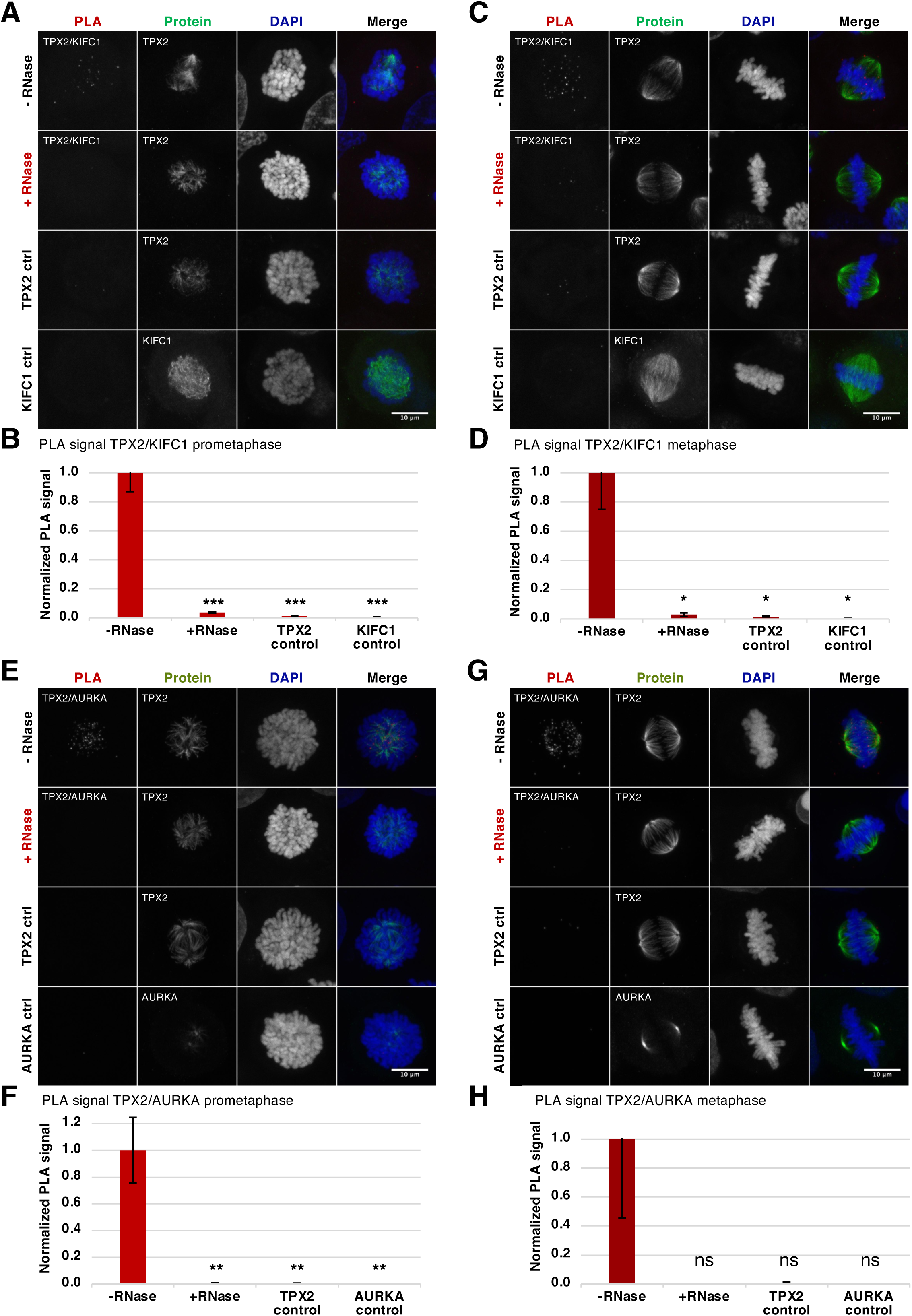
RNA-dependent protein interactions in mitosis demonstrated by Proximity Ligation Assay (PLA) **A**: PLA images showing the proximity of TPX2 and KIFC1 in HeLa cells at prometaphase (PLA channel, red dots in the merge channel). The protein (TPX2 or KIFC1) is seen in green (immunostaining) and DNA was stained using DAPI (blue). The two top images depict representative images in control cells and RNase-treated cells. The two bottom images depict representative images of the individual antibodies PLA controls. Scale bars, 10 µm. **B**: Bar graph of the quantification of the PLA signal for the proximity of TPX2 and KIFC1 in HeLa prometaphase cells in absence (-RNase) or presence (+RNase) of RNase treatment, as well as in the individual antibody control PLA assays. The signal intensities in each sample were normalized to the signal intensity of the -RNase sample. The error bars indicate the SEM (N=3). The *P*-value were calculated using a two-tailed, unpaired t-test (* *P*-value <0.05, ** *P*-value <0.01, *** *P*-value <0.001, ns not significant). **C** and **D**: same as in **A** and **B** respectively, at metaphase (N=3). **E** and **F**: same as in **A** and **B**, for TPX2 and AURKA in HeLa cells synchronized in prometaphase (N=3). **G** and **H**: same as in **A** and **B**, for TPX2 and AURKA in HeLa cells synchronized in metaphase (N=3). In panels B, D and H, the upper error bar, which have the same size as the lower error bars, were removed to represent all “-RNase” reference bars with the same height for better comparison.

**Supplementary Figure S7.**
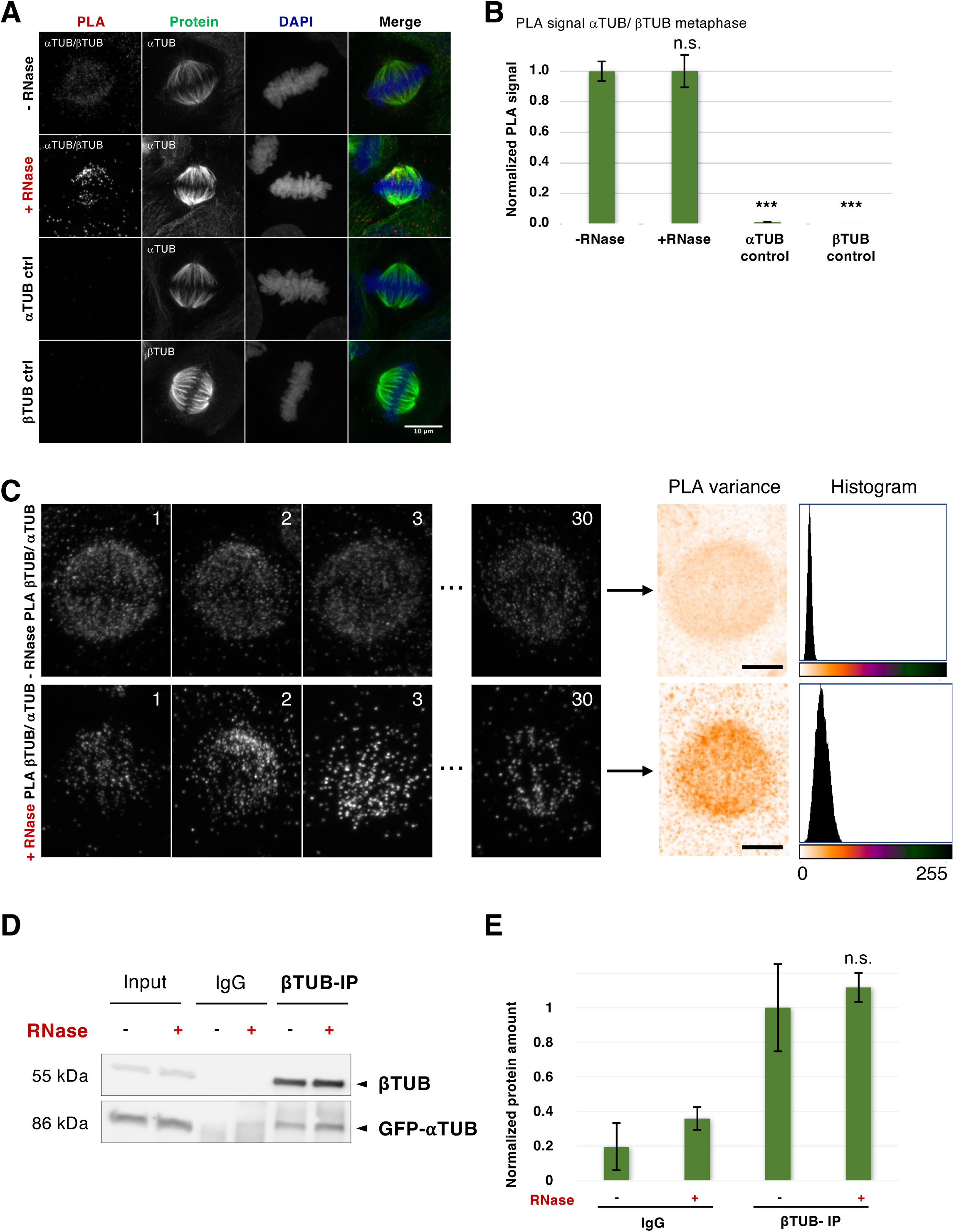
Interaction between α– and β– tubulin upon RNase treatment. **A:** PLA images showing the proximity of α– and β– tubulin in HeLa cells synchronized in prometaphase (PLA channel, red dots in the merge channel). The protein (the indicated tubulin) is seen in green (immunostaining) and DNA was stained using DAPI (blue). The two top images depict representative images in control cells and RNase-treated cells. The two bottom images depict representative images of the individual antibodies PLA controls. Scale bars, 10 µm. **B**: Bar graph of the quantification of the PLA signal for the proximity of α– and β– tubulin in HeLa metaphase cells in absence (-RNase) or presence (+RNase) of RNase treatment, as well as in the individual antibody control PLA assays. The signal intensities in each sample were normalized to the signal intensity of the -RNase sample. The error bars indicate the SEM (N=3). The *P*-value were calculated using a t-test (*** *P*-value <0.001, ns not significant). The PLA signal for the proximity of α– and β– tubulin is not significantly decreasing upon RNase treatment, as compared to the untreated (-RNase) sample. **C**: Differential analysis of the PLA signal for the proximity of α– and β– tubulin. Left panel: images of three replicates (3 x 10 images) were rotated, scaled to the average pole-to-pole distance, cropped and superposed to compute an image (orange) representing the variance of the PLA signal in the two series of images (right panel). As seen from the PLA images, the PLA variances and the histograms, the PLA signal for the proximity of α- and β-tubulin after RNase treatment is remarkably inhomogeneously distributed, especially throughout the spindle area (greater variance values). Scale bar, 5 µm. **D**: WB analysis of the pulldown of β-tubulin (βTUB) and co-pulldown of α-tubulin tagged with GFP (GFP-αTUB) in HeLa prometaphase cell lysates treated with or without RNase. IgG was used as a negative control. αTUB was detected using an anti-GFP antibody. **E**: Bar plot showing the quantification of the pulldown of β-tubulin (βTUB) and co-pulldown of α-tubulin tagged with GFP (GFP-αTUB) in HeLa prometaphase cell lysates treated with or without RNase, as in D. The error bars represent the SEM (N=3). The *P*-value was calculated using a two-tailed, unpaired t-test (n.s., not significant).

**Supplementary Figure S8.**
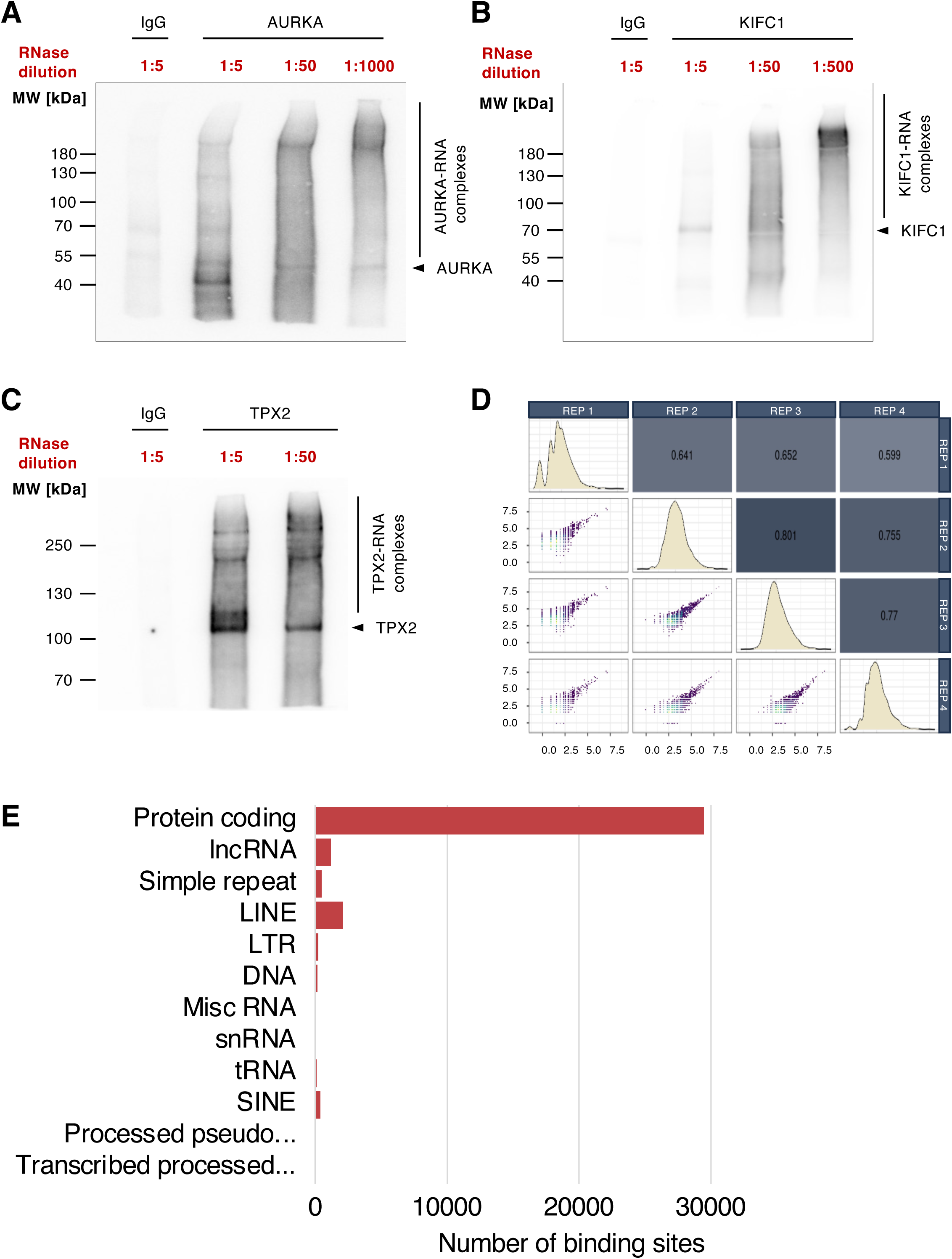
KIFC1 and AURKA preferentially bind protein coding RNAs in prometaphase. **A** and **B**: Autoradiography indicating the direct binding of AURKA and KIFC1, respectively, to RNA by iCLIP2 as indicated by shifting of the radioactive RNA signal towards higher molecular weights with decreasing RNase I concentrations in A549 prometaphase cells (representative images out of N=3 replicates are shown). **C**: Composite graph representing the pairwise correlations between the KIFC1 iCLIP2-Seq replicates as scatter plot (bottom left), the Pearson correlation coefficient (upper right) and the coverage distribution as density (diagonal). **D**: Horizontal bar plot indicating the number of binding sites within the various categories of non-rRNA target genes from the KIFC1 iCLIP2-Seq in HeLa prometaphase cells. The genes are classified as protein coding, lncRNA, simple repeat, LINE, LTR, DNA, Misc RNA, snRNA, tRNA, SINE, processed pseudogene and transcribed processed pseudogene.

**Supplementary Figure S9.**
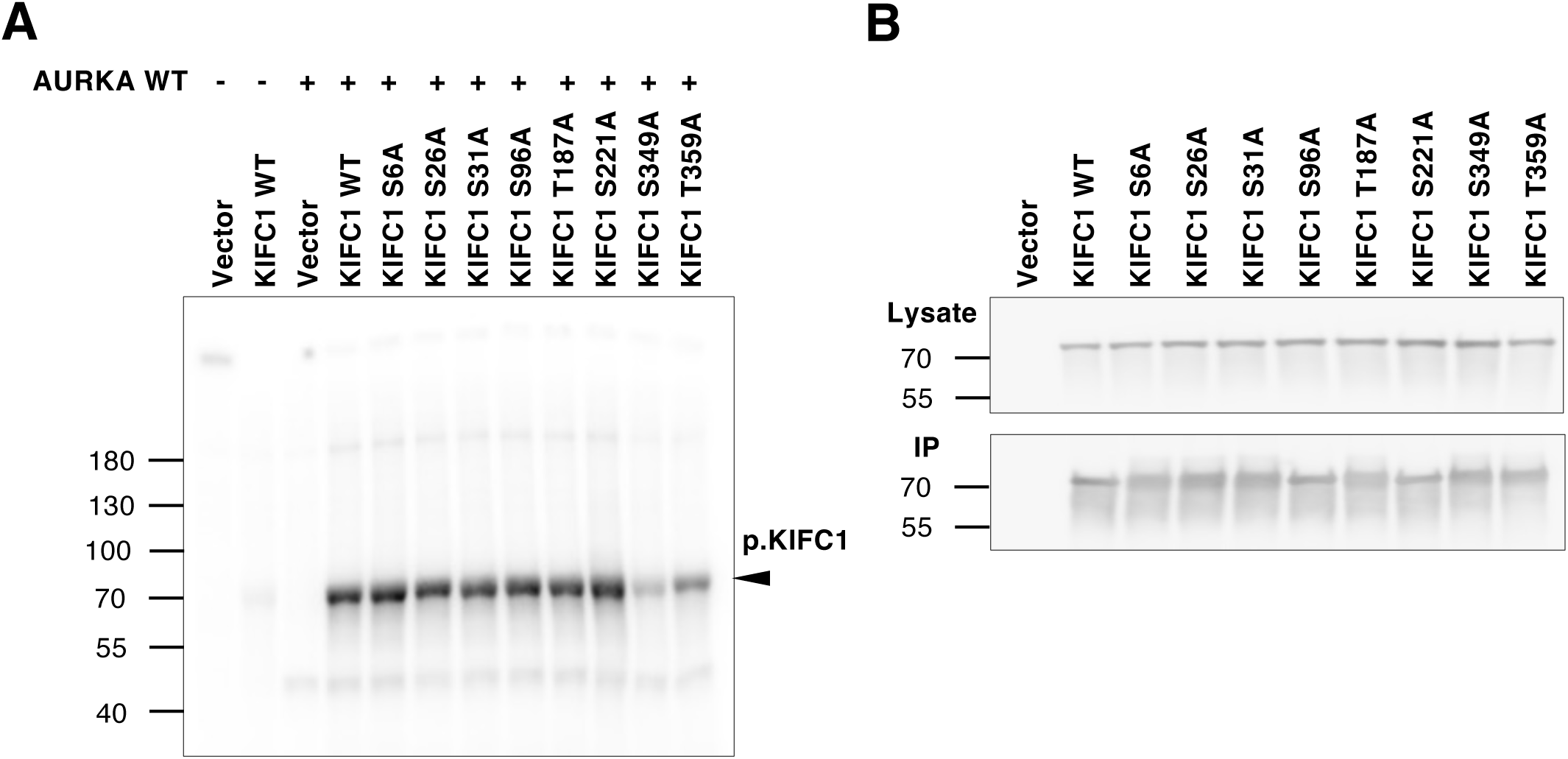
AURKA phosphorylates KIFC1 at S349 and T359. **A**: Autoradiography indicating the phosphorylation intensity of KIFC1 in wild-type (WT) and non-phosphorylatable KIFC1 mutants in the presence of purified AURKA WT. The *in vitro* kinase assay was performed using KIFC1 pulled down from HeLa prometaphase lysates overexpressing the WT or mutant KIFC1 proteins with an N-terminal Flag-HA tag. An empty vector was used as a negative control. The single mutations of the residues are indicated above each lane. **B**: WB analysis of the expression and pulldown efficiency of Flag-HA-tagged KIFC1 WT and mutants (as in A) using anti-DYKDDDDK magnetic agarose beads for *in vitro* kinase assay in HeLa cell lysates.

