## Supplementary Tables and R-DeeP3 User Guide for "An atlas of RNA-dependent proteins in cell division reveals the riboregulation of mitotic protein-protein interactions": Rajagopal_et_al_Supplementary_Table_S5.pdf

### Supplementary Table S5: List of reagents

| Experimental models & Cell lines | Catalogue number | Company |
|---|---|---|
| HeLa | CCL-2 | ATCC |
| A549 | CCL-185 | ATCC |

| Reagents or Resources | Catalogue number | Lot number | Company |
|---|---|---|---|
| DMEM High Glucose medium | D65796 | N/A | Sigma Aldrich |
| RPMI 1640 medium | 11875093 | N/A | Gibco, Thermo Fisher Scientific |
| Dulbecco's Phosphate Buffered Saline (PBS) | D8537 | N/A | Sigma Aldrich |
| Luria Broth Base (LB) | 12795-027 | 200806062408 | Invitrogen |
| Human KIFC1 ORF | HG15958-G | N/A | Sino Biologicals |
| NEB 5-alpha Competent E. coli (High Efficiency) | C2987H | N/A | New England Biolabs (NEB) |
| One Shot TOP10 Chemically Competent cells E. coli | C404003 | N/A | Thermo Fisher Scientific |
| Q5 Site-Directed Mutagenesis Kit | E0554S | N/A | New England Biolabs (NEB) |
| E. coli Ribonuclease H (RNase H), 2 U/µl | 8021014 | | Thermo Fisher Scientific |
| Nucleospin plasmid isolation kit | 740588.250 | 1707/002 | Macherey Nagel |
| GeneJET gel extraction kit | K0692 | 00439790 | Thermo Scientific |
| Gateway™ BP Clonase™ II Enzyme-Mix | 11789100 | N/A | Thermo Fisher Scientific |
| Gateway TM LR Clonase TM II Enzyme Mix | 11791020 | N/A | Thermo Fisher Scientific |
| Proteinase K | AM2546 | | Ambion |
| 2x Phusion High-Fidelity PCR MasterMix | M0531S | N/A | New England Biolabs (NEB) |
| Tissue culture dish 146 x 21 mm (15 cm) | 93150 | N/A | Techno Plastic Products |
| Thymidine cell culture tested | T1895-1G | N/A | Sigma Aldrich |
| Methyl-(5-(2-thienylcarbonyl)-1H-*benzim (Nocodazole) | M1404-2MG | N/A | Sigma Aldrich |

| | | | |
|---|---|---|---|
| 4–20% Criterion™ TGX Stain-Free™ Protein Gel, 26 well, 15 µl | 5678095 | N/A | BioRad |
| 7.5% precast Mini-Protean-TGX gel, 10 wells | 456-1024 | N/A | BioRad |
| 7.5% precast Mini-Protean-TGX gel, 12 wells | 456-1025 | N/A | BioRad |
| 7.5% precast Mini-Protean-TGX gel, 15 wells | 456-1026 | N/A | BioRad |
| NuPAGE LDS Sample Buffer | NP0007 | 2538211 | Thermo Fisher Scientific |
| Page Ruler Prestained Protein Ladder | 26616 | N/A | Thermo Fisher Scientific |
| Pager Ruler Plus Prestained Protein Ladder | 26619 | N/A | Thermo Fisher Scientific |
| Bovine Serum Albumin (BSA) | A1470 | 10042749 | Sigma-Aldrich |
| Bicinchoninic Acid solution | B9643-1L | N/A | Sigma-Aldrich |
| Pierce™ BCA Protein Assay Reagent B | 23224 | TA260103 | Thermo Fisher Scientific |
| Amersham™ Protran® Western blotting membranes, nitrocellulose | 10600002 | N/A | Sigma-Aldrich |
| 5x Trans-Blot Turbo Transfer Buffer | 10026938 | N/A | Bio-Rad |
| HRP-conjugated Goat anti-mouse secondary antibody | 115-035-003 | 108284 | Dianova |
| HRP-conjugated Goat anti-rabbit secondary antibody | 111-035-144 | 149084 | Dianova |
| ECL prime Western blotting system | RPN2232 | N/A | Cytiva |
| Proteus clarification mini spin column | 42225.01 | N/A | Serva |
| Pierce ChIP-grade Protein A/G Magnetic Beads | 26162 | XH353638 | Thermo Fisher Scientific |
| Dynabeads Protein A | 10002D | 01268693 | Thermo Fisher Scientific |
| Dynabeads Protein G | 10004D | 01265132 | Thermo Fisher Scientific |

| | | | |
|---|---|---|---|
| DynaMagTM-2 magnet | 12321D | N/A | Thermo Fisher Scientific |
| Complete, EDTA-free Protease Inhibitor Cocktail | 4693132 | 73791000 | Sigma-Aldrich |
| PhosStop | 4906845001 | 73124700 | Sigma-Aldrich |
| TURBO DNase | AM2238 | 01341010 | Thermo Fisher Scientific |
| RNase I 100 U/µl | AM2295 | 01317147 | Thermo Fisher Scientific |
| Normal rabbit IgG | 12-370 | N/A | Millipore |
| Aurora A (D3E4Q) Rabbit monoclonal antibody | 14475 | Lot2 | Cell Signaling Technology |
| KIFC1 antibody (11445), Rb monoclonal antibody | 172620 | GR3259790-3 | Abcam |
| Anti-TPX2, Clone TPX2-01 | SAB4701065 | 539147 | Sigma-Aldrich |
| Anti-TPX2 (18D5) | 628002 | B256728 | Biolegend |
| Cell culture plate, 12-well | 92412 | N/A | Techno Plastic Products |
| Microscope cover glasses, 12 mm, Nr. 1.5 | 0112520 | 51687 | Neolab |
| Duolink® In Situ Detection Reagents Red | DUO92008 | 0000326527 | Sigma-Aldrich |
| Duolink® In Situ Wash Buffers, Fluorescence | DUO82049 | N/A | Sigma-Aldrich |
| Duolink® In Situ PLA® Probe Anti-Mouse MINUS, Affinity purified Donkey anti-Mouse IgG (H+L) | DUO92004 | 0000337233 | Sigma-Aldrich |
| Duolink® In Situ PLA® Probe Anti-Rabbit PLUS, Affinity purified Donkey anti-Rabbit IgG (H+L) | DUO92002 | SLCR1898 | Sigma-Aldrich |
| Duolink In Situ Mounting medium | DUO82040-5ML | 0000339465 | Sigma-Aldrich |
| Alexa Fluor 488 goat anti-rabbit IgG | A11034 | VG302077 | Thermo Fisher Scientific |
| Alexa Fluor 488 goat anti-mouse IgG | A32723 | VH309036 | Thermo Fisher Scientific |
| T4 PNK 10U/µl 500 µl | M0201 | 10129472 | New England Biolabs (NEB) |
| ATP 10 mM 1 ml | P0756 | 10109057 | New England Biolabs (NEB) |

| | | | |
|---|---|---|---|
| [gamma-P32] Adenosine 5'-triphosphate (ATP) /9,25 MBq | SRP-501 | | Hartmann Analytic |
| SUPERase-In RNase Inhibitor (20 U/µL)-10,000 units | AM2696 | | Thermo Fisher Scientific |
| Phenol-chloroform-Isoamylalkohol pH 6.5-6.9 | P3803 | | Sigma-Aldrich |
| Heavy Phase Lock Gel tubes | 733-2478 | | SERVA |
| T4 RNA Ligase 1 (ssRNA Ligase), High Concentration | M0437M | | NEB |
| Superscript IV reverse transcriptase | 18090050 | | Thermo Fisher Scientific |
| MyONE silane beads | 37002D | | Thermo Fisher Scientific |
| RLT Buffer | 79216 | | Qiagen |
| RNaseOUT, 40 U/µl | 10777-019 | | |
| ProNex® Size-Selective Purification System | NG2001 | | Promega |
| RNA Clean & Concentrator™-5 (50 Preps) w/ Zymo-Spin™ IC Columns (Capped) | R1015 | | Zymo Research |
| Ribolock RNase Inhibitor (40 U/µL) | EO0384 | 00965329 | Thermo Fisher Scientific |
| Lipofectamine 2000 Transfection Reagent | 11668500 | | Thermo Fisher Scientific |
| Opti-MEM(R) I Reduced Serum Medium | 31985047 | | Thermo Fisher Scientific |
| Pierce™ Anti-DYKDDDDK Magnetic Agarose | A36797 | YD368599 | Thermo Scientific |
| Human AURKA protein | N/A | N/A | EMBL Protein Expression and Purification Core Facility |
| Human AURKA-D274A protein | N/A | N/A | EMBL Protein Expression and Purification Core Facility |

| **Others** | **Catalogue Number** | **Company** |
|---|---|---|
| Imager ECL Chemo Cam CC5569 | N/A | INTAS |
| Stratalinker 2400 | N/A | Stratagene |
| SW 40 Ti Swinging-Bucket Rotor | 331302 | Beckman Coulter |

| | | |
|---|---|---|
| Ultra-Clear Tube | 344060 | Beckman Coulter |
| Orbitrap Fusion LC-MS/MS platform | N/A | Thermo Fisher Scientific |
| Fusion Orbitrap Lumos mass spectrometer | N/A | Thermo Fisher Scientific |
| Discovery 90SE Ultracentrifuge | N/A | Sorvall |
| Sequencing Grade Modified Trypsin | V5113 | Promega |
| TMTsixplex™ Isobaric Label Reagent Set | 90066 | Thermo Fisher Scientific |

| **Softwares** | | |
|---|---|---|
| Bioconductor | Gentleman et al., 2004 | www.bioconductor.org |
| Comet | Eng et al., 2013 | http://comet-ms.sourceforge.net |
| ImageJ | Schneider et al., 2012 | https://imagej.net/Downloads |
| LabImage 1D 2006 | Kapelan Bio-Imaging GmbH | www.labimage.com |
| Microsoft Excel | Microsoft | www.microsoft.com |
| Primer Blast | NCBI | https://www.ncbi.nlm.nih.gov/tools/primer-blast/ |
| R programming | The R project | www.r-project.org |
| Shiny | R Studio | https://shiny.rstudio.com |
| GO Enrichment Analysis | Thomas PD et al., 2022 | https://geneontology.org/ |
| Serial cloner | Serial Basics | http://serialbasics.free.fr/Serial_Cloner.html |
| GIMP | The GNU Project | https://www.gimp.org |
| Ensemble | Harrison et al., 2024 | https://www.ensembl.org/index.html |

| **Primers and other sequences** | | | |
|---|---|---|---|
| **Cloning primers** | **Sequence** | | **Company** |
| KIFC1 Forward | GGGGACAAGTTTGTACAAAAAAGCAGGCT TCATGGATCCGCAGAGGTCCCCCTATTG | Primers to generate Entry clone for Gateway cloning. | Sigma-Aldrich |
| KIFC1 Reverse | GGGGACCACTTTGTACAAGAAAGCTGGGTT CTATCACTTCCTGTTGGCCTGAGCAGTACC | | Sigma-Aldrich |
| S6A_FP | TCCGCAGAGGGCCCCCTATTGG | | Sigma-Aldrich |
| S6A_RP | TCCATTCACTTCCTGTTGGCCTGAGC | | Sigma-Aldrich |
| S26A_FP | TAAGGCCCCTGCCCAGCTGCCTC | | Sigma-Aldrich |
| S26A_RP | ATCAGAGGTCTCTTCAGTTCTATGTTCCCCT TTAC | | Sigma-Aldrich |
| S31A_FP | GCTGCCTCTCGCAGGAAGCAGAC | | Sigma-Aldrich |
| S31A_RP | TGGGAAGGGGCCTTAATCAG | | Sigma-Aldrich |
| S349A_FP | AACCCGCCTTGCGCTCTCCCGGTCTGAC | | Sigma-Aldrich |
| S349A_RP | GGAGGATCAGAGGGCCCA | | Sigma-Aldrich |
| T359A_FP | GCGGCGTGGGGCCCTGAGTGGGG | | Sigma-Aldrich |
| T359A_RP | TCGTCAGACCGGGAGAGGCTAAGGCG | | Sigma-Aldrich |
| pFastBac_KIFC1-FP | ATGGATCCGGAATTCAAAG | Primer for NEBuilder Cloning to generate KIFC1 for expression in insect cells | Sigma-Aldrich |
| pFastBac_KIFC1-RP | GGCGCCCTGAAAATACAG | | Sigma-Aldrich |
| KIFC1-FP | ACCTGTATTTTCAGGGCGCCATGGATCCGC AGAGGTCC | | Sigma-Aldrich |
| KIFC1-RP | CCTTTGAATTCCGGATCCATTCACTTCCTGT TGGCCTG | | Sigma-Aldrich |
| pFastBac_AURKA-FP | ACAGTCTTAGTAATCAGCCATACCACATTTG | Primer for NEBuilder Cloning to generate AURKA for expression in insect cells | Sigma-Aldrich |
| pFastBac_AURKA-RP | ATCGGTCCATGGCGCCCTGAAAATACAG | | Sigma-Aldrich |
| AURKA-FP | TCAGGGCGCCATGGACCGATCTAAAGAAA AC | | Sigma-Aldrich |
| AURKA-RP | TGGCTGATTACTAAGACTGTTTGCTAGC | | Sigma-Aldrich |
| D274A_FP | TAAAATTGCAGCTTTTGGGTGGTC | | Sigma-Aldrich |
| D274A_RP | AGCTCTCCAGCTGATCCA | | Sigma-Aldrich |

| iCLIP2 primers, adapters or barcode sequences | | | |
|---|---|---|---|
| **Adapters or barcodes or primers** | **Sequence** | | **Company** |
| L3-App | /5rApp/AG ATC GGA AGA GCG GTT CAG /3ddC/ | | IDT |
| RT oligo | GGATCCTGAACCGCT | | Sigma-Aldrich |
| P3Solexa_s | CACGACGCTCTTCCGATCT | | Sigma-Aldrich |
| P5Solexa_s | CTGAACCGCTCTTCCGATCT | | Sigma-Aldrich |
| P3Solexa | AATGATACGGCGACCACCGAGATCTACACT CTTTCCCTACACGACGCTCTTCCGATCT | | Sigma-Aldrich |
| P5Solexa | CAAGCAGAAGACGGCATACGAGATCGGTC TCGGCATTCCTGCTGAACCGCTCTTCCGATC T | | Sigma-Aldrich |
| L02clip2.0 | /5Phos/NN NNC GAT GTN NNN NAG ATC GGA AGA GCG TCG TG/3ddC/ | | IDT |
| L05clip2.0 | /5Phos/NN NNA CAG TGN NNN NAG ATC GGA AGA GCG TCG TG/3ddC/ | | IDT |
| L10clip2.0 | /5Phos/NN NNT AGC TTN NNN NAG ATC GGA AGA GCG TCG TG/3ddC/ | | IDT |
| L19clip2.0 | /5Phos/NN NNG TGA AAN NNN NAG ATC GGA AGA GCG TCG TG/3ddC/ | | IDT |
