## Supplementary Tables and R-DeeP3 User Guide for "An atlas of RNA-dependent proteins in cell division reveals the riboregulation of mitotic protein-protein interactions": Rajagopal_et_al_UserGuide_RDeeP3.pdf

#### R-DeeP 3.0 - Database for RNA-Dependent Proteins (https://R-DeeP3.dkfz.de)

R-DeeP 3.0 is a database for RNA-dependent proteins based on the concept of "RNA dependence" and its direct experimental application.

A protein is defined as "RNA-dependent" if its interactome depends on RNA without necessarily directly binding to RNA as described previously [1][2]. Briefly, RNA-dependent proteins and complexes migrate to different positions in a sucrose density gradient in the control versus the RNase-treated lysates, according to their apparent molecular weight (i.e., according to the size of the complex of which they are part of). This allows the specific and quantitative discovery of RNA-dependent proteins [1][3]. R-DeeP 3.0 is the result of the statistical analysis of proteome-wide mass spectrometry data to determine the behavior of proteins in a sucrose density gradient in presence of RNA molecules and after RNase treatment analyzed in human unsynchronized HeLa S3 and A549 cells (previous datasets) and HeLa S3 cells synchronized in mitosis or interphase (new datasets).

R-DeeP 3.0 provides various search and download options for in-depth analyses, which are further described in this User Guide (**Illustration 1**). In addition, R-DeeP 3.0 offers a summary of multiple RNA-binding protein resources with access to the corresponding publication webpage.

The database R-DeeP 3.0 has been optimized for use with Firefox or Safari browsers.

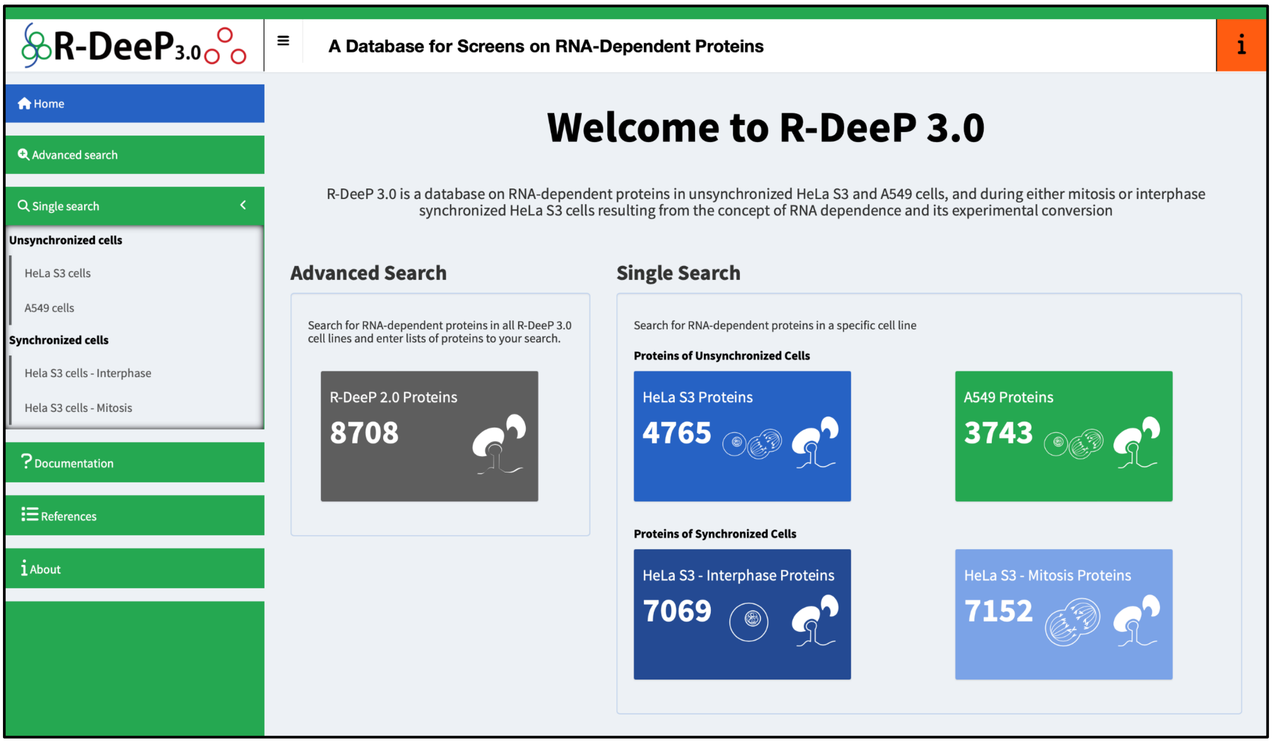


**Illustration 1**. R-DeeP 3.0 Homepage at https://r-deep3.dkfz.de/. The info boxes for all datasets are clickable and bring the user directly to the Advanced Search or respective Single Search option for each dataset and cell line (see below).

### 1. Advanced Search

The Advanced search option allows comparing a protein directly between the four datasets. In addition, it provides support for a list of proteins as an input format. Such a list can be copied and pasted into the search field (**Illustration 2**).

The protein names, gene names, Uniprot IDs or other aliases such as accession numbers from HGNC and MIM databases can be entered into the search field and the search is started when the "Enter" key or the "Submit" button is pressed (**Illustration 3**). As soon as the "Submit" button or "Enter" key is pressed, the sidebar is closed to provide more space to display the results. Pressing on the "Sidebar" button will re-open the sidebar. Alternatively, the sidebar can be activated using the small icon right from the R-DeeP 3.0 image at the top left side of the page.

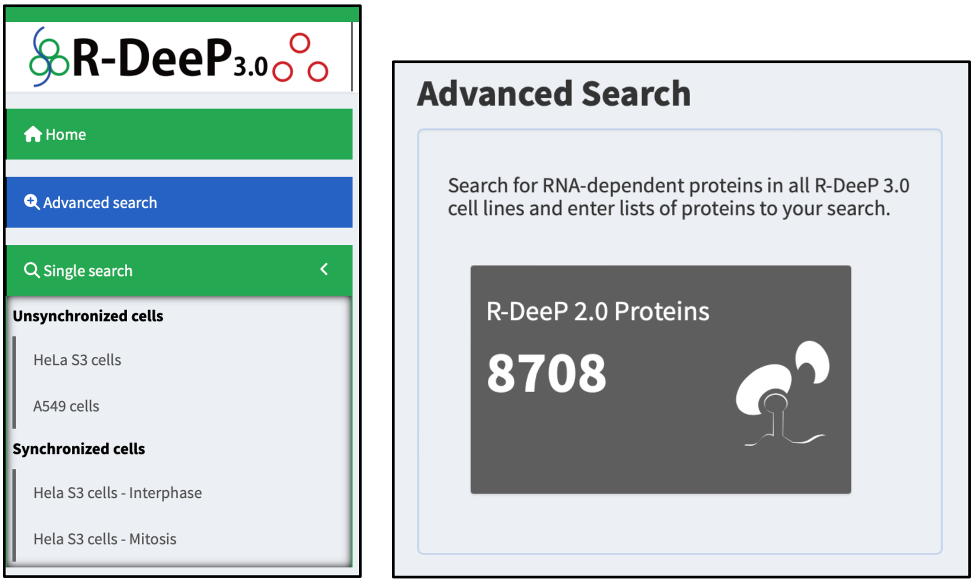


**Illustration 2.** Advanced search selection from the sidebar (left) or from the info box (right).

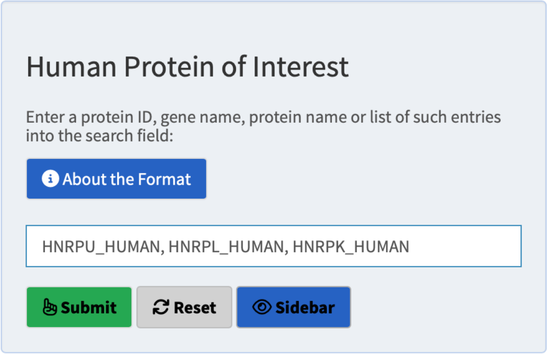


**Illustration 3**. Advanced search input field.

In the “About the Format” pop-up window, more detailed information about the input format is provided (**Illustration 4**). The pop-up window contains a link to the UniProt database, that could be helpful to find the proper protein or gene name in case they are not known. It also provides more information about the format requirements of a protein list. Besides the regular option to search for a protein or gene name, it is possible to copy and paste a list of proteins from a table. The format of the list needs to strictly adhere to the formatting instructions mentioned in the “About the format” panel (**Illustration 4**).

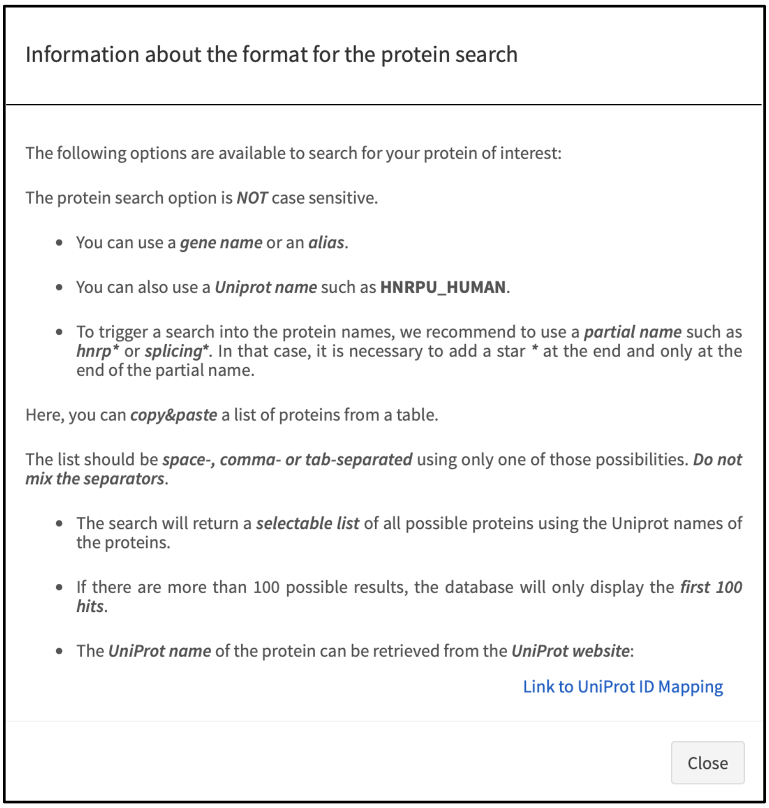


**Illustration 4**. Information about the search format.

In case of a successful search, a new panel with search results appears with either a unique result (**Illustration 5**) or a list of suggested hits (**Illustration 6**). The search result already indicates whether data is available for this protein and whether the protein is shifting (green comment next to the protein name, clickable button), whether there is data available but the protein does not shift (yellow comment, clickable button) or whether there is no data available, *i.e.* the protein has not been detected by mass spectrometry (red comment, no clickable button, **Illustration 7**). The protein of interest, if available, can be selected to obtain the corresponding comprehensive R-DeeP 3.0 analysis.

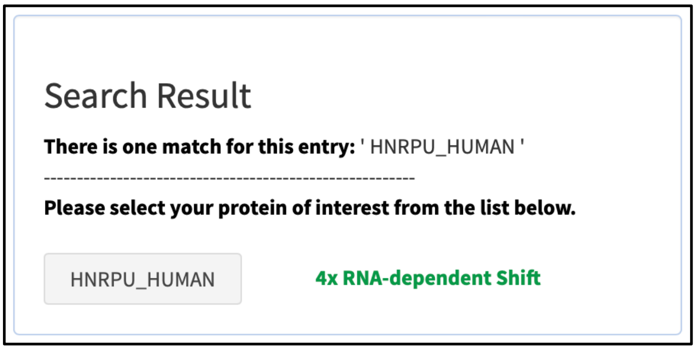


**Illustration 5.** Advanced search for a single protein.

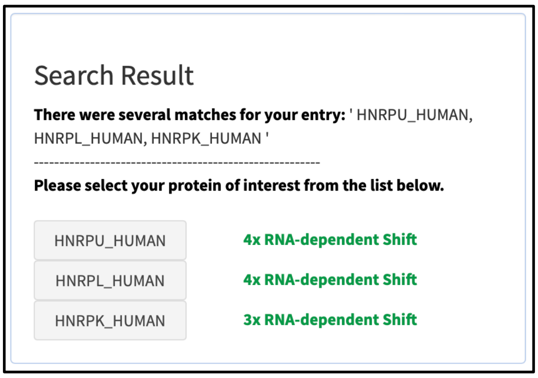


**Illustration 6**. Advanced search for a list of proteins.

In addition, if the search result displays a list of proteins, it directly indicates whether data is available for the protein and whether it is shifting. However, since the search occurs in all datasets at one time, there are multiple options (**Illustration 7**):

- **4x, 3x or 2x RNA-dependent Shift**: RNA-dependent shift in 4, or in at least 3 or 2 datasets (green comment next to the protein name, clickable button).
- **Shift HeLa**: RNA-dependent shift in HeLa S3 unsynchronized dataset. Similarly, there are proteins with "**Shift A549**", "**Shift HeLa Mitosis**" and "**Shift HeLa Interphase**" (green comment next to the protein name, clickable button). In that case, there is a shift observed in only one of the datasets. For the other datasets, it can be "data available" or "no data" (see below).
- **4x, 3x or 2x Data**: no RNA-dependent shift in 4, 3 or 2 datasets. There is no data available for the other datasets (yellow comment, clickable button).
- **Data HeLa Mitosis**: no RNA-dependent shift in HeLa S3 synchronized in mitosis (yellow comment, clickable button) and no data in any of the other experiment, *i.e.* not detected. Similarly, there are proteins with "**Data HeLa**", "**Data A549**" and "**Data HeLa Interphase**" (yellow comment, clickable button).
- **No data**: no data available for this protein in either dataset (red comment, no clickable button (**Illustration 7**).

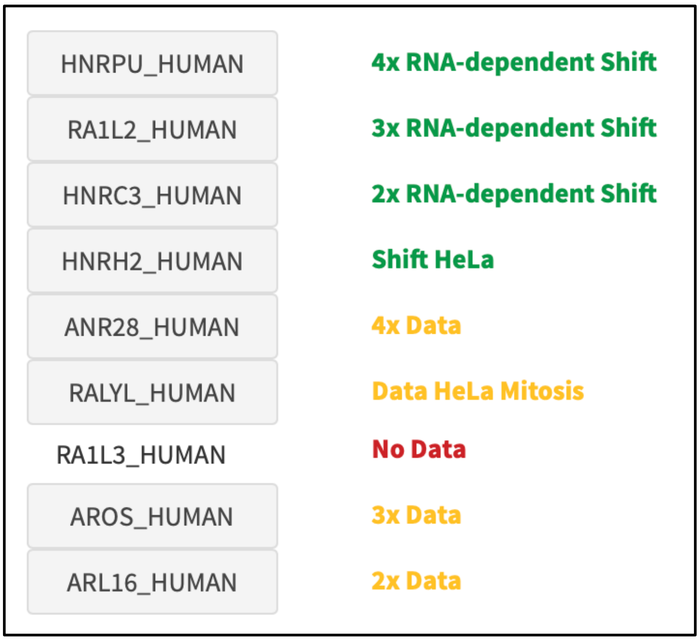

**Illustration 7.** Example of search results in the Advanced search option.

The search returns an error message in case there are no results (**Illustration 8**).

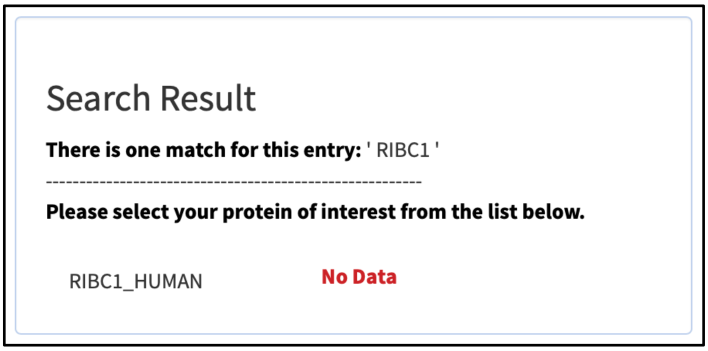


**Illustration 8**. Example of protein that is not available in the mass spectrometry data.

If there are more than 100 proteins found, the search returns the results for the first 100 matches. Alternatively, the search needs to be refined (**Illustration 9**).

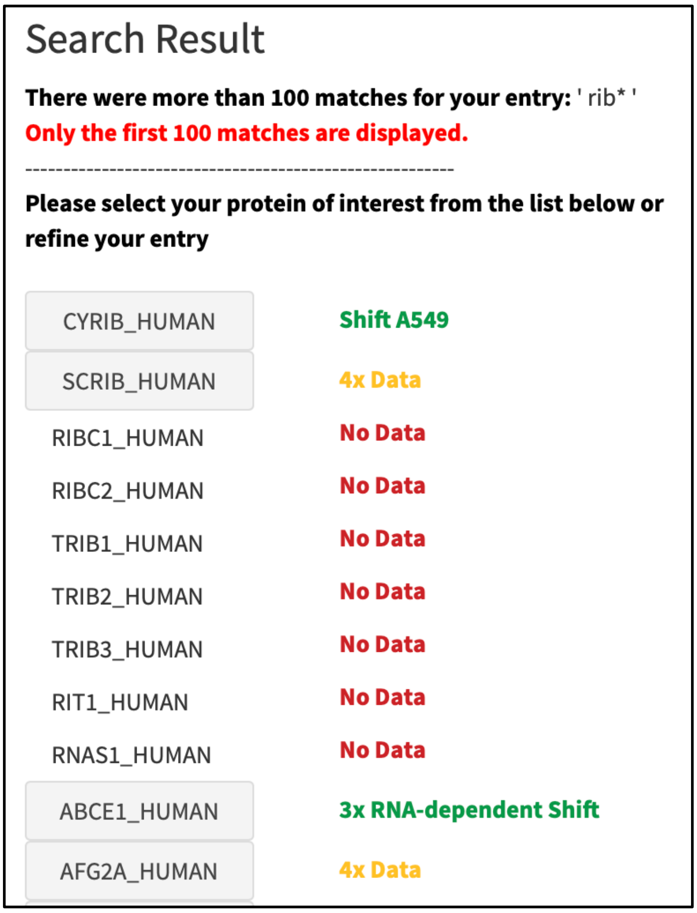


**Illustration 9**. Search result for more than 100 matches.

By clicking on a specific protein (button), the complete R-DeeP 3.0 analysis results (**Illustration 10**) appear in a new panel, which is composed of several elements.

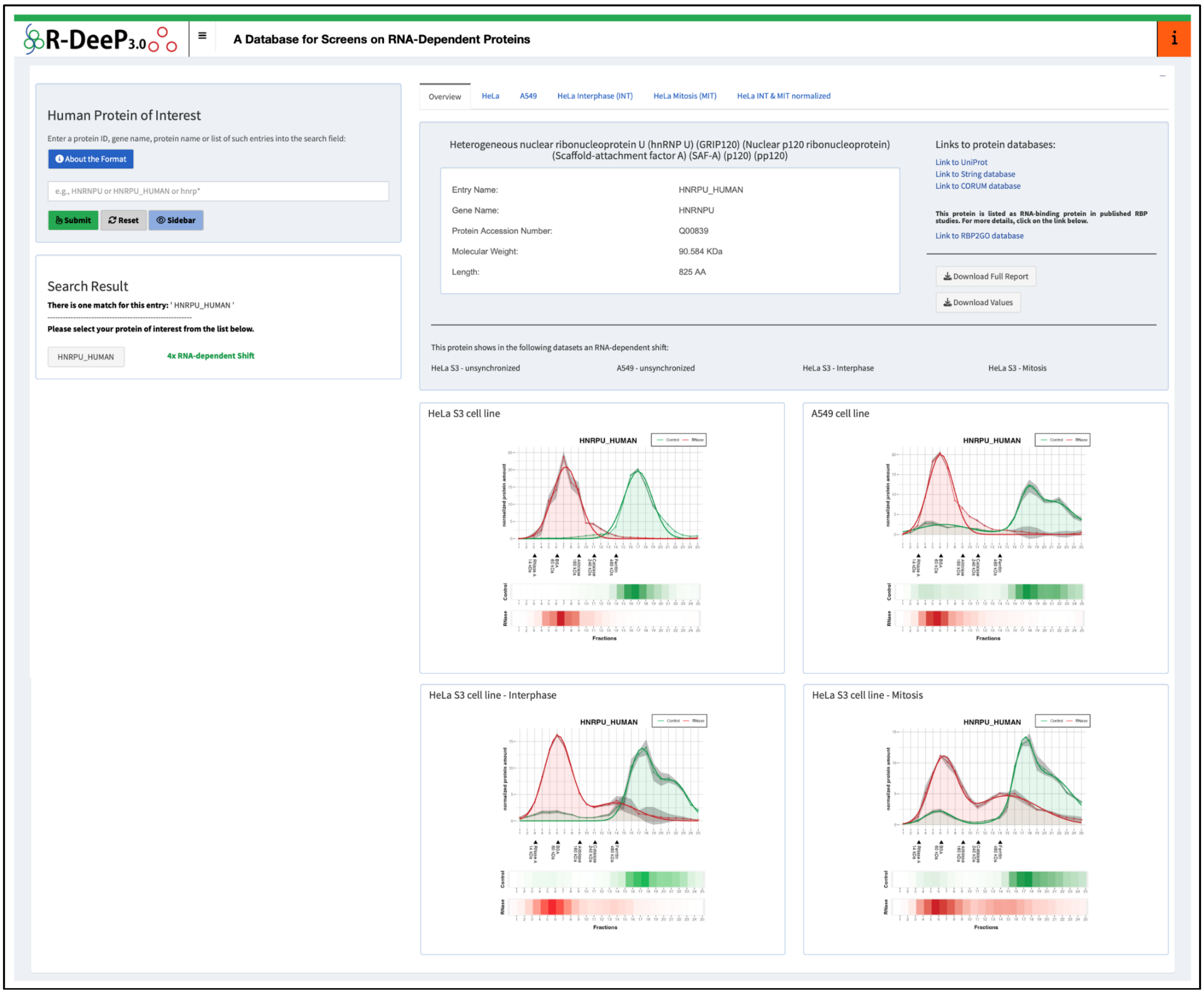


**Illustration 10**. R-DeeP 3.0 Advanced search analysis results for hnRNP U. On the left side, the "Search Result" panel indicates that the protein depicts four RNA-dependent shifts (*i.e.*, in all four datasets). On the right side, details on the protein are given in various tabs.

For each protein in the "Advanced search", the following information is depicted, if available (additional panels):

- **Overview**: general information about the protein and if available, graphical representation for each dataset. Downloads options are offered.
- **For each additional tab (HeLa, A549, HeLa Interphase (INT)**, **HeLa Mitosis (MIT)**, **HeLa INT & MIT normalized)**: On the left-hand side, the top panel provides the graphical representation of the distribution of the protein in the gradients loaded with control (in green) and RNase-treated (in red) lysates. In the line graph, the respective green and red thin lines are the mean raw data curves for three replicates and the stronger lines are the Gaussian fits of the mean raw data. The grey areas around the curves represent the standard deviation for the raw data calculated from the three replicates. The graph also indicates (black arrows) the position of the standard proteins that were used to calibrate the gradient for the HeLa S3 cell line (**Illustration 11**). Three buttons at the top provide various download options:

- "Download Plot": a PDF of the graphical representation of the distribution of the protein

- “Download Values”: a comma-separated values (CSV) file of the raw data and standard deviation for each fraction
- Download “Full Report”: a complete analysis report in html format

The user can open a larger view of the graphical representation by clicking the “Zoom” button (top left corner of the graph panel).

On the lower left-hand side, the results of the statistical analysis are displayed. Lists of the maxima as well as their position and corresponding amount of protein for the control and RNase gradients are shown. If significant shifts were detected, the parameters of the shifts are indicated.

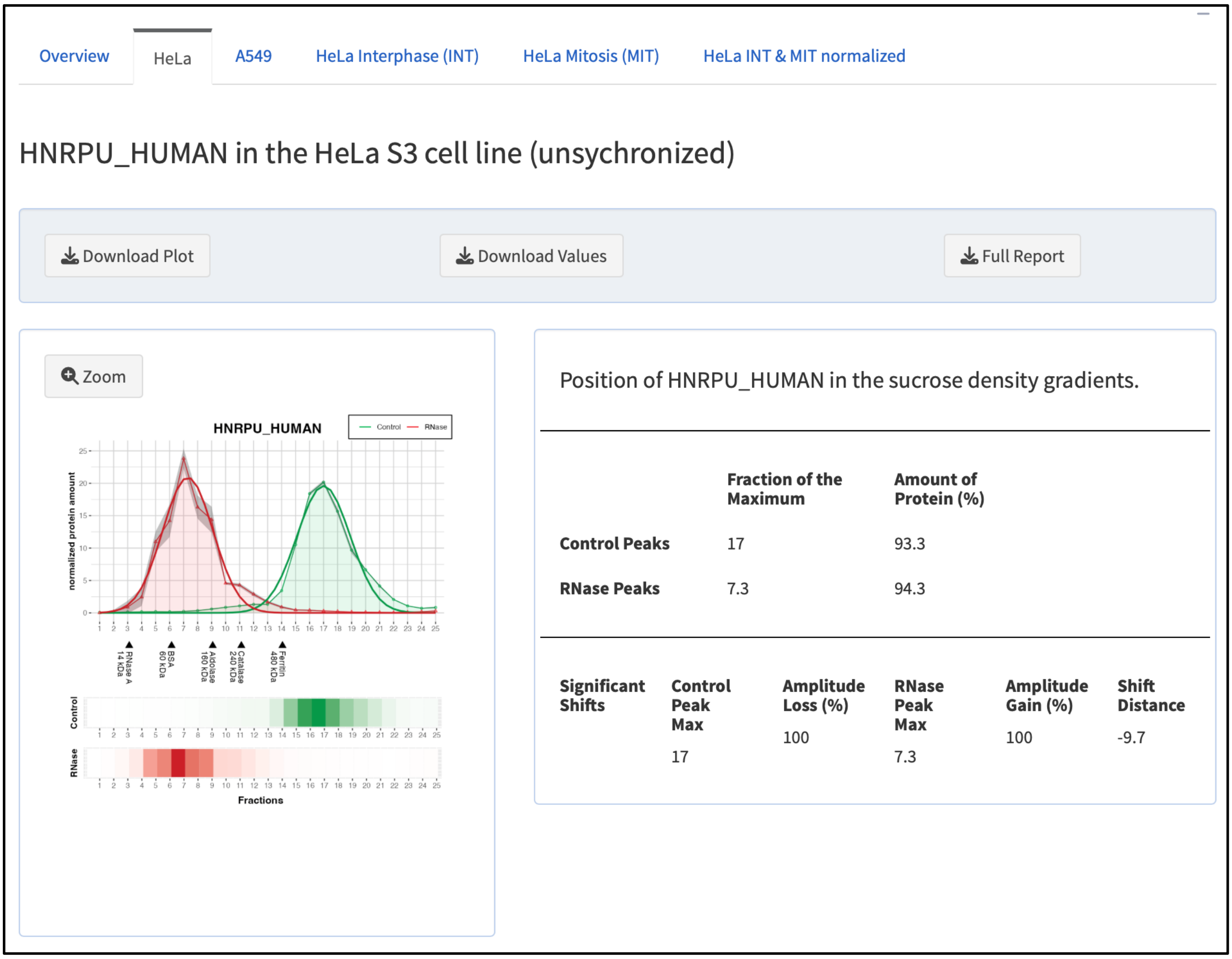


**Illustration 11**. R-DeeP 3.0 results for hnRNP U in HeLa S3 cells.

#### 2. Single search

This search option allows searching the database for one protein of interest in either of the datasets: HeLa S3 or A549 unsynchronized, HeLa S3 synchronized in Interphase or Mitosis (**Illustration 12**). Once the dataset is selected, it is possible to obtain information for a protein of interest by using the input field (**Illustration 13**).

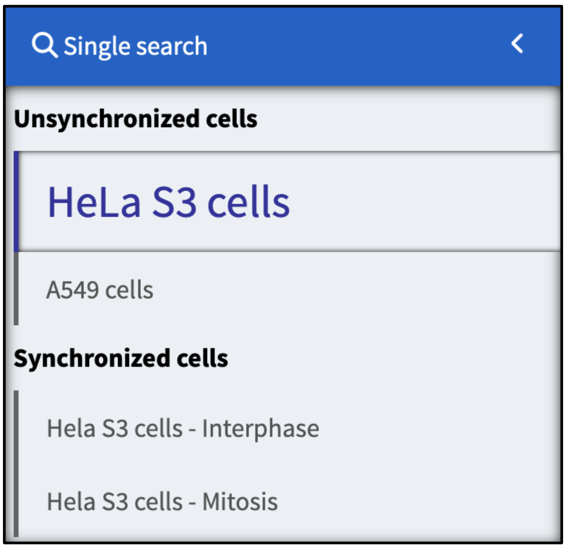


**Illustration 12.** Dataset selection.

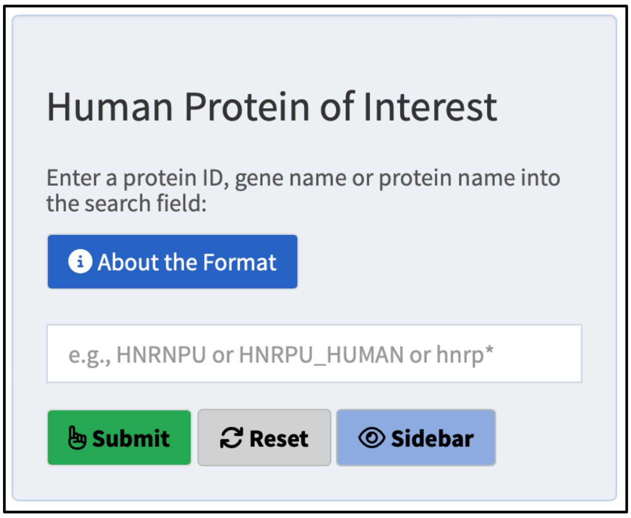


**Illustration 13**. Single protein search input field.

In the "About the Format" pop-up window, more detailed information about the input format is provided. (**Illustration 14**). The pop-up window contains a link to the UniProt database, that could be helpful to find the proper protein or gene name in case they are not known.

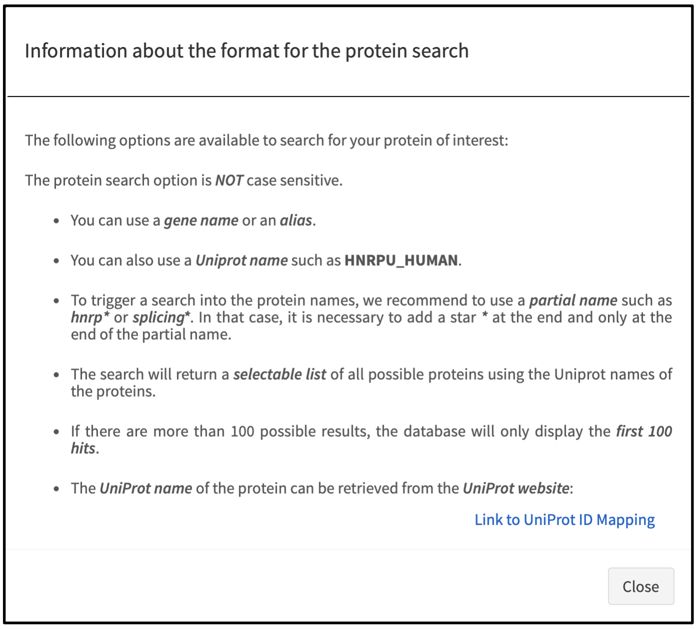


**Illustration 14**. Information about the search format.

In case of a successful search, a new panel with search results appears with either a unique result (**Illustration 15**) or a list of suggested hits (**Illustration 16**). In case of more than 100 hits, the list contains the first 100 hits (**Illustration 16**). The search result already indicates whether data is available for this protein and whether the protein is shifting (green comment next to the protein name, clickable button), whether there is data available but the protein does not shift (yellow comment, clickable button) or whether there is no data available, *i.e.* the protein has not been detected by mass spectrometry (red comment, no clickable button, **Illustration 17**). The protein of interest, if available, can be selected to obtain the corresponding comprehensive R-DeeP 3.0 analysis.

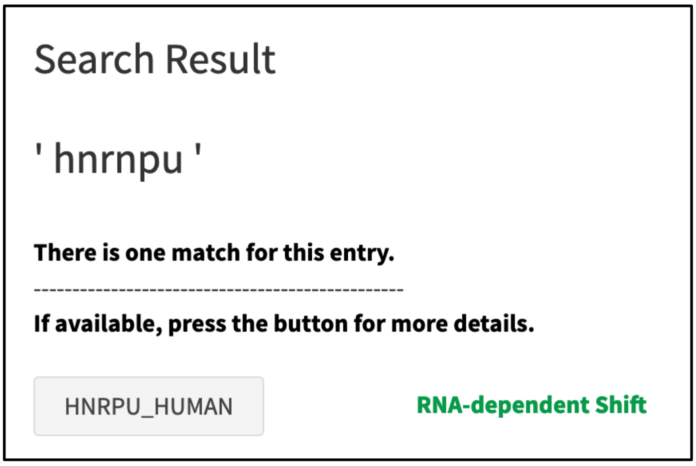


**Illustration 15.** Result of a search with one match.

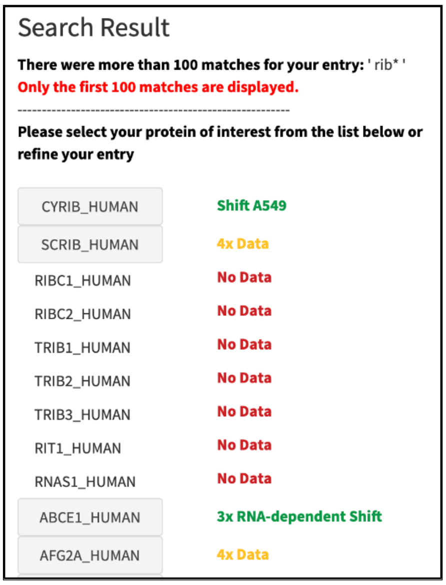


**Illustration 16**. Result list with multiple hits including message indicating a restriction to the first 100 hits.

The search returns an error message in case there are no results (**Illustration 18**).

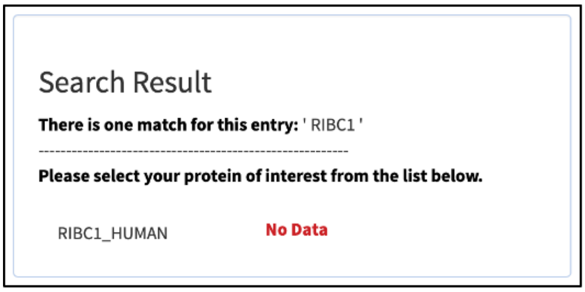


**Illustration 17**. Example of a protein that is not available in the mass spectrometry data.

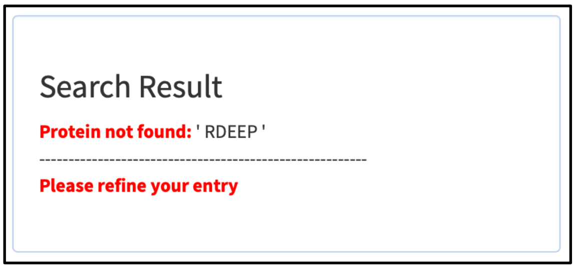


**Illustration 18**. Error message if no hits are found.

The complete R-DeeP 3.0 analysis results (**Illustration 19**) appear in a new panel, which is composed of several elements.

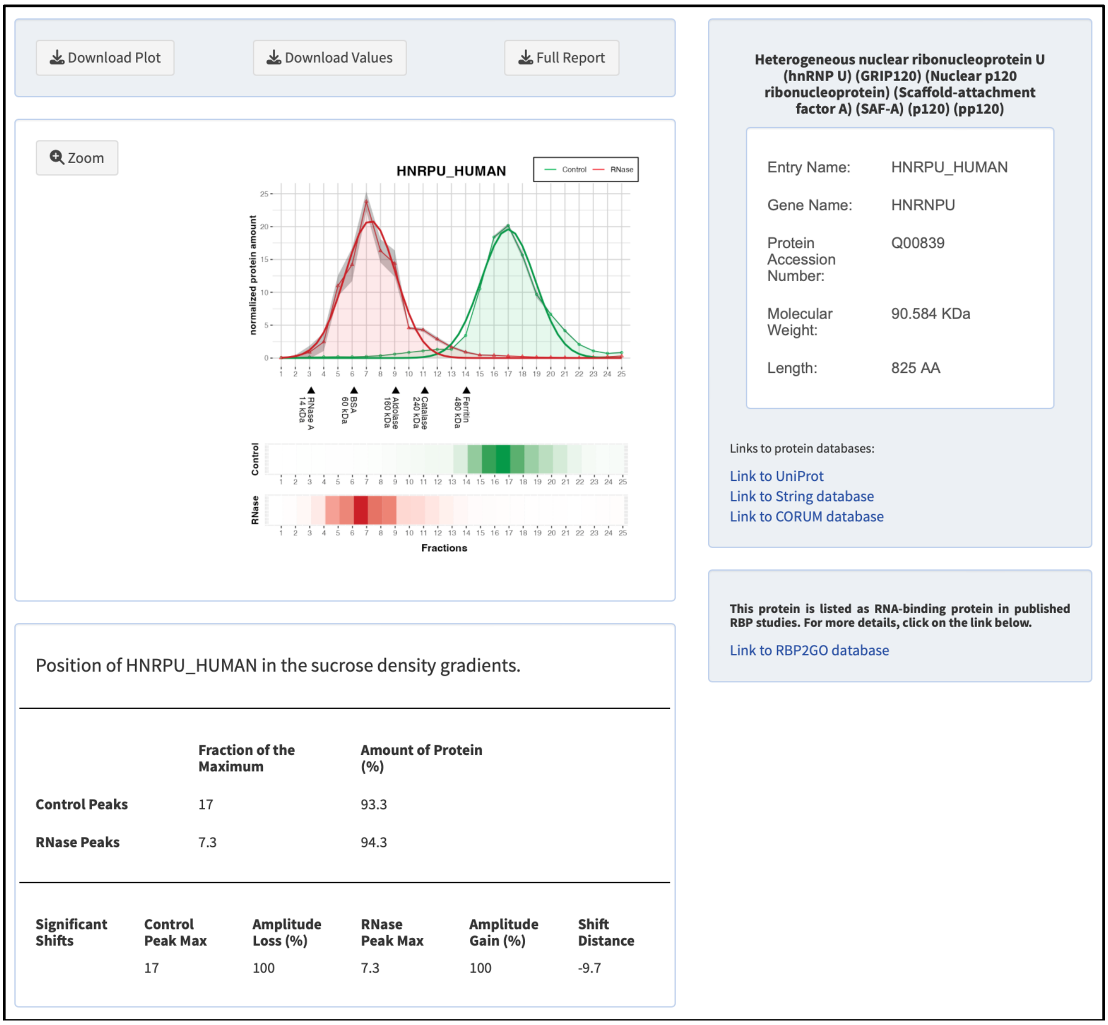


**Illustration 19**. R-DeeP 3.0 analysis results for hnRNP U.

On the left-hand side, the top panel provides the graphical representation of the distribution of the protein in the gradients loaded with control (in green) and RNase-treated (in red) lysates. In the line graph, the respective green and red thin lines are the mean raw data curves for three replicates and the stronger lines are the Gaussian fits of the mean raw data. The grey areas around the curves represent the standard deviation for the raw data calculated from the three replicates. The graph also indicates (black arrows) the position of the standard proteins that were used to calibrate the gradient for the HeLa S3 cell line.

Three buttons at the top provide various download options:

- “Download Plot”: a PDF of the graphical representation of the distribution of the protein
- “Download Values”: a comma-separated values (CSV) file of the raw data and standard deviation for each fraction
- Download “Full Report”: a complete analysis report in html format

The user can open a larger view of the graphical representation by clicking the “Zoom” button (top left corner of the graph panel).

On the lower left-hand side, the results of the statistical analysis are displayed. Lists of the maxima as well as their position and corresponding amount of protein for the control and RNase gradients are shown. If significant shifts were detected, the parameters of the shifts are indicated.

On the right-hand side, a panel provides information about the protein itself. The top panel offers basic information on the protein as found on the UniProt database as well as direct links to the UniProt page of the protein and the CORUM and STRING database for further information about this protein and its interaction partners (**Illustration 20**). In addition, the lower panel indicates whether the protein of interest has already been listed as potential RNA-binding protein (RBP) in previous studies. A link brings the user directly to the RBP2GO database [4][5] in a new tab.

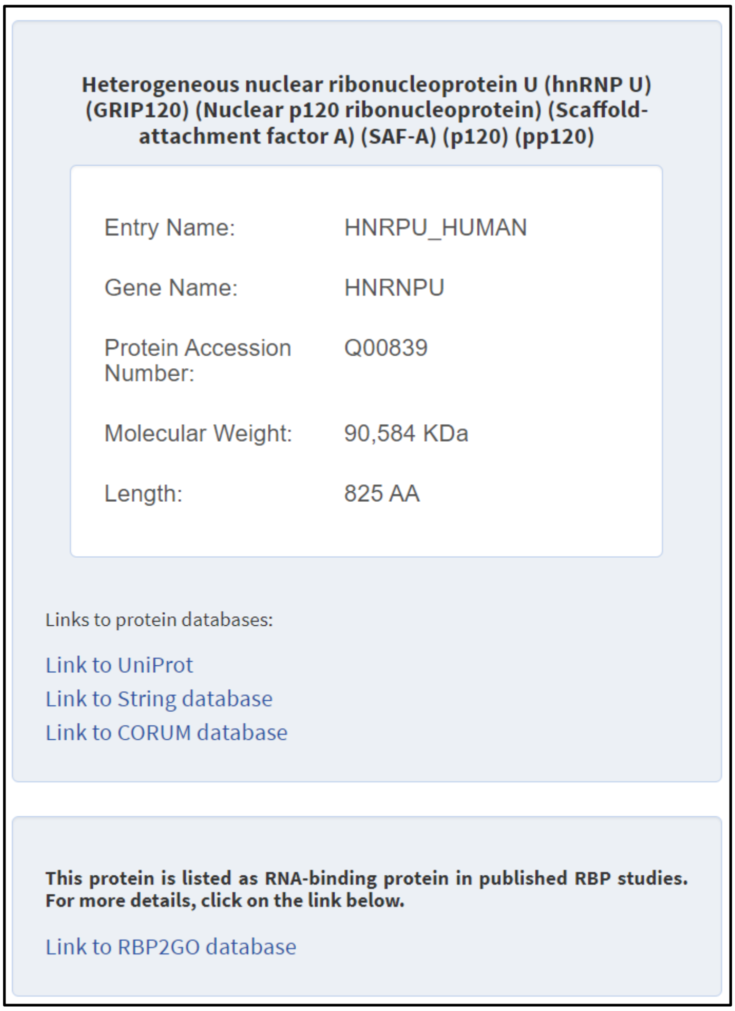


**Illustration 20**. Panel with specific information on the protein and link to protein databases as well as R-DeeP 3.0 tab for RBP resources.

For the HeLa S3 datasets synchronized in interphase or mitosis, the search results offer the possibility to visualize both results in separate panels at the same time if the protein is available in both datasets (**Illustration 21**). For this visualization, the protein amounts in the graphs were normalized to the cycle-phase for which the protein had the highest amount (**Illustration 22**).

Info Mitosis | Info Mitosis & Interphase

Download Plot | Download Values | Full Report

Zoom

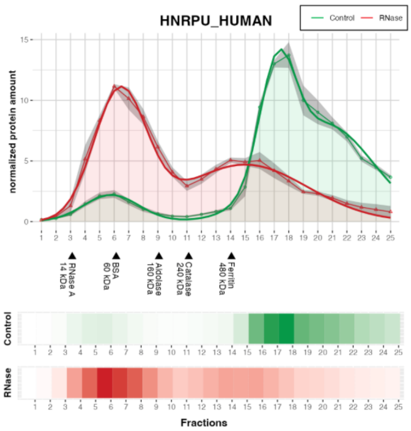


Position of HNRPU_HUMAN in the sucrose density gradients.

| | Fraction of the Maximum | Amount of Protein (%) |
|---|---|---|
| **Control Peaks** | 17.2 | 27.7 |
| | 20.5 | 60.8 |
| **RNase Peaks** | 6.4 | 47.3 |
| | 15 | 50.2 |

| Significant Shifts | Control Peak Max | Amplitude Loss (%) | RNase Peak Max | Amplitude Gain (%) | Shift Distance |
|---|---|---|---|---|---|
| | 17.2 | 70.7 | 6.4 | 81.2 | -10.8 |
| | 20.5 | 74.7 | 6.4 | 81.2 | -14.1 |
| | 17.2 | 70.7 | 15 | 21.1 | -2.2 |
| | 20.5 | 74.7 | 15 | 21.1 | -5.5 |

**Heterogeneous nuclear ribonucleoprotein U (hnRNP U) (GRIP120) (Nuclear p120 ribonucleoprotein) (Scaffold-attachment factor A) (SAF-A) (p120) (pp120)**

| | |
|---|---|
| Entry Name: | HNRPU_HUMAN |
| Gene Name: | HNRNPU |
| Protein Accession Number: | Q00839 |
| Molecular Weight: | 90.584 KDa |
| Length: | 825 AA |

Links to protein databases:

Link to UniProt

Link to String database

Link to CORUM database

**This protein is listed as RNA-binding protein in published RBP studies. For more details, click on the link below.**

Link to RBP2GO database

**Illustration 21**. Panel with specific information on the protein and link to protein databases as well as R-DeeP 3.0 tab for RBP resources.

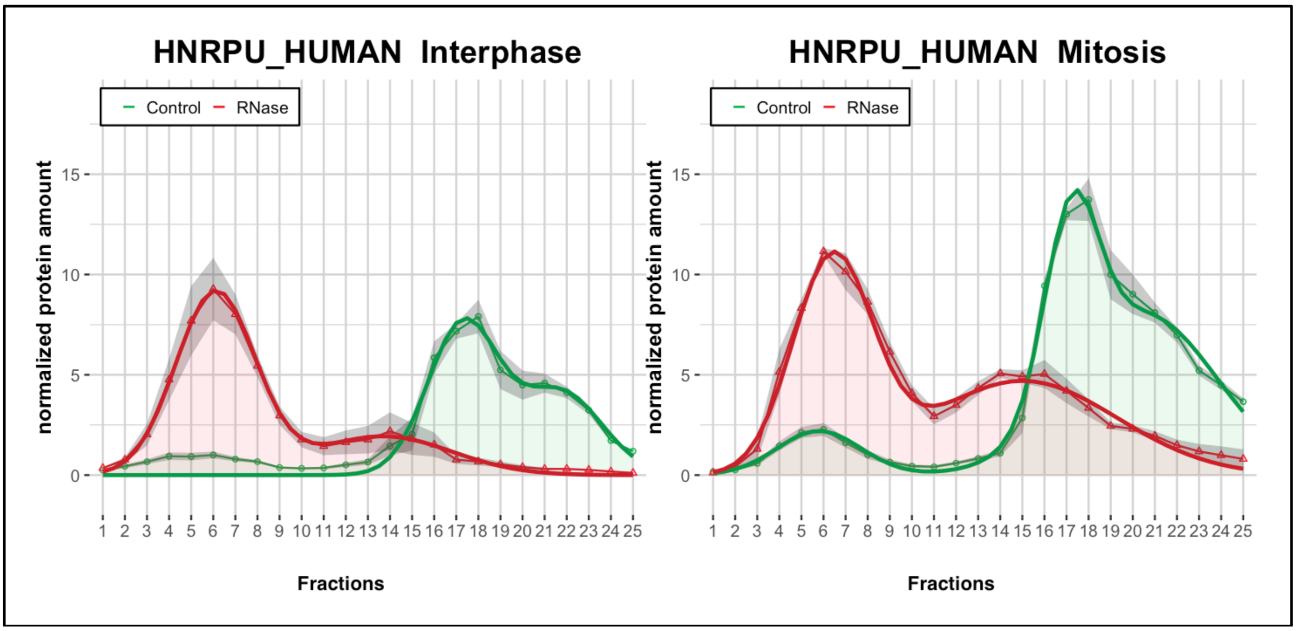


**Illustration 22**. Graphical representation of the normalized protein amounts in HeLa S3 interphase and mitosis.

#### 3. Other tabs: Links / Documentation / About

The remaining three tabs on the R-DeeP 3.0 homepage (**Illustration 23**) summarize important links, the documentation including this User Guide as well as general information about the R-DeeP 3.0 database including reference to the publication and contact information.

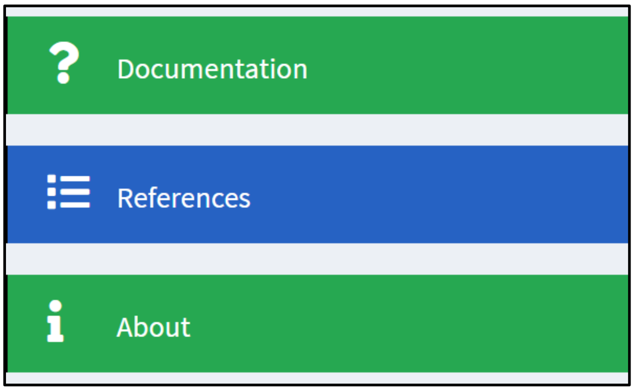


**Illustration 23**. Documentation, References and About tabs.

### 4. Example of an R-DeeP 3.0 full report for hnRNP U

#### R-DeeP 3.0 Analysis report for interphase or mitosis synchronized HeLa S3 cell line

**Research Group "RNA-Protein Complexes & Cell Proliferation"**

German Cancer Research Center (DKFZ) - Heidelberg - Germany

**July 22, 2024**

http://r-deep3.dkfz.de
https://rbp2go-2-beta.dkfz.de

##### UniProt name of the protein: HNRPU_HUMAN

Gene Name: HNRNPU
Gene Name Synonyms: HNRNPU C1orf199 HNRPU SAFA U21.1
Protein Accession Number: Q00839
Molecular Weight: 90.584 KDa
Length: 825 AA
Description: Heterogeneous nuclear ribonucleoprotein U (hnRNP U) (GRIP120) (Nuclear p120 ribonucleoprotein) (Scaffold-attachment factor A) (SAF-A) (p120) (pp120)

##### RNA-Interaction of the protein: HNRPU_HUMAN

This protein is listed as RNA-binding protein in published RBP studies. For more details, click on the link below.
https://rbp2go-2-beta.dkfz.de

This protein shows in the following R-DeeP 3.0 datasets an RNA-dependent shift:

HeLa S3 (unsynchronized), A549 (unsynchronized), HeLa S3 - Interphase, HeLa S3 - Mitosis

##### Position of the protein in the sucrose density gradient - HeLa S3 (unsynchronized)

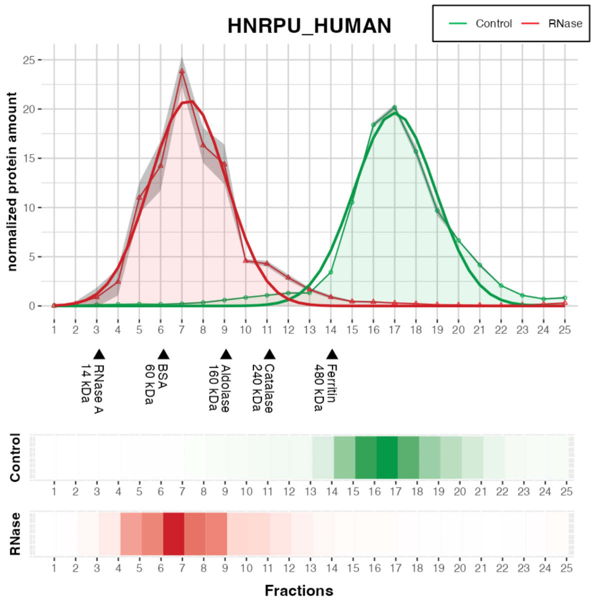

#### Peaks and shifts information - HeLa S3 (unsynchronized)

Control Peaks

| Fraction of the Maximum | Amount of Protein (%) |
|---|---|
| 17 | 93.3 |

RNase Peaks

| Fraction of the Maximum | Amount of Protein (%) |
|---|---|
| 7.3 | 94.3 |

Significant Shifts

| Control Peak Max | Amplitude Loss (%) | RNase Peak Max | Amplitude Gain (%) | Shift Distance |
|---|---|---|---|---|
| 17 | 100 | 7.3 | 100 | -9.7 |

#### Position of the protein in the sucrose density gradient - A549 (unsynchronized)

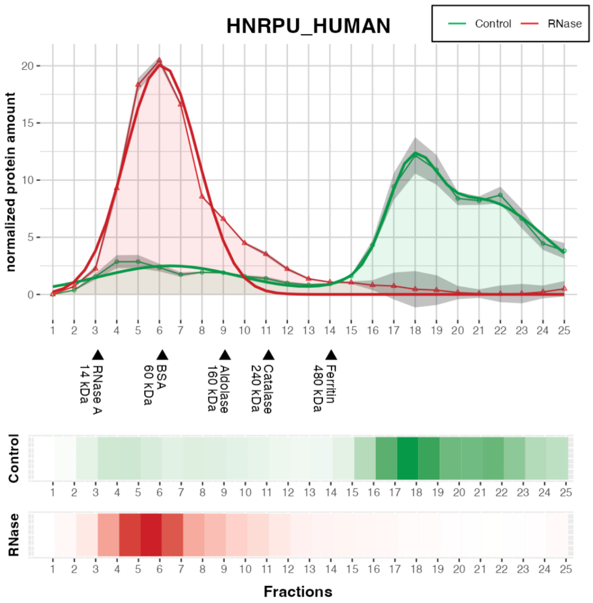


#### Peaks and shifts information - A549 (unsynchronized)

Control Peaks

| Fraction of the Maximum | Amount of Protein (%) |
|---|---|
| 6.5 | 20.2 |
| 17.8 | 20.1 |
| 21.0 | 58.5 |

RNase Peaks

| Fraction of the Maximum | Amount of Protein (%) |
|---|---|
| 6.1 | 85.6 |

Significant Shifts

| Control Peak Max | Amplitude Loss (%) | RNase Peak Max | Amplitude Gain (%) | Shift Distance |
|---|---|---|---|---|
| 17.8 | 100 | 6.1 | 87.6 | -11.7 |
| 21.0 | 100 | 6.1 | 87.6 | -14.9 |

#### Position of the protein in the sucrose density gradient - HeLa S3 - Interphase

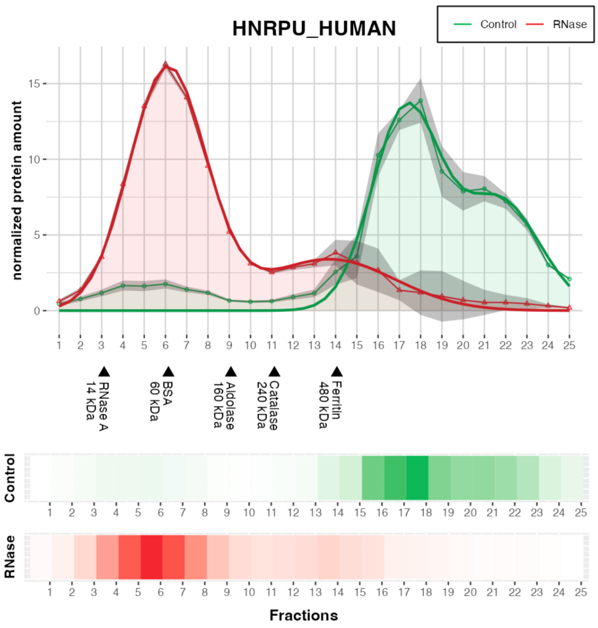


#### Peaks and shifts information - HeLa S3 - Interphase

Control Peaks

| Fraction of the Maximum | Amount of Protein (%) |
|---|---|
| 17.3 | 52.9 |
| 21.7 | 33.5 |

RNase Peaks

| Fraction of the Maximum | Amount of Protein (%) |
|---|---|
| 6.1 | 72.1 |

Significant Shifts

| Control Peak Max | Amplitude Loss (%) | RNase Peak Max | Amplitude Gain (%) | Shift Distance |
|---|---|---|---|---|
| 17.3 | 87.4 | 6.1 | 100 | -11.2 |
| 21.7 | 98.4 | 6.1 | 100 | -15.6 |

Position of the protein in the sucrose density gradient - HeLa S3 - Mitosis

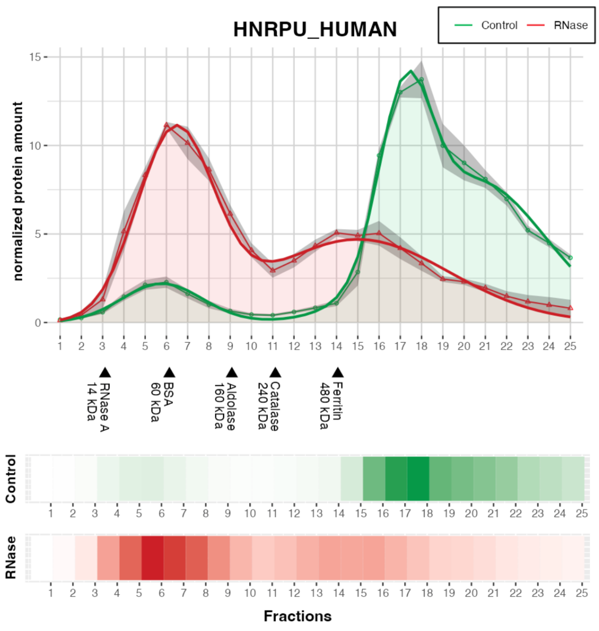


Peaks and shifts information - HeLa S3 - Mitosis

Control Peaks

| Fraction of the Maximum | Amount of Protein (%) |
|---|---|
| 17.2 | 27.7 |
| 20.5 | 60.8 |

RNase Peaks

| Fraction of the Maximum | Amount of Protein (%) |
|---|---|
| 6.4 | 47.3 |
| 15.0 | 50.2 |

Significant Shifts

| Control Peak Max | Amplitude Loss (%) | RNase Peak Max | Amplitude Gain (%) | Shift Distance |
|---|---|---|---|---|
| 17.2 | 70.7 | 6.4 | 81.2 | -10.8 |
| 20.5 | 74.7 | 6.4 | 81.2 | -14.1 |
| 17.2 | 70.7 | 15.0 | 21.1 | -2.2 |
| 20.5 | 74.7 | 15.0 | 21.1 | -5.5 |

### 5. Contact information

The authors of the R-DeeP 3.0 database can be contacted at the following email address:
